# Structural Organization of the Retriever-CCC Endosomal Recycling Complex

**DOI:** 10.1101/2023.06.06.543888

**Authors:** Daniel J. Boesch, Amika Singla, Yan Han, Daniel A. Kramer, Qi Liu, Kohei Suzuki, Puneet Juneja, Xuefeng Zhao, Xin Long, Michael J. Medlyn, Daniel D. Billadeau, Zhe Chen, Baoyu Chen, Ezra Burstein

**Affiliations:** Roy J. Carver Department of Biochemistry, Biophysics & Molecular Biology, Iowa State University, 2437 Pammel Drive, Ames, IA 50011, USA; Department of Internal Medicine, University of Texas Southwestern Medical Center, 5323 Harry Hines Boulevard, Dallas, TX 75390, USA; Department of Biophysics, University of Texas Southwestern Medical Center, 6001 Forest Park Road, Dallas, TX 75390, USA; Cryo-EM facility, Office of Biotechnology, Iowa State University, 2437 Pammel Drive, Ames, IA 50011, USA; Research IT, College of Liberal Arts and Sciences, Iowa State University, 2415 Osborn Dr, Ames, IA 50011, USA; Division of Oncology Research, College of Medicine, Mayo Clinic, Rochester MN, 55905, USA; Department of Molecular Biology, University of Texas Southwestern Medical Center, 5323 Harry Hines Boulevard, Dallas, TX 75390, USA

**Keywords:** Endosome recycling, Retromer, Retriever, VPS35L, VPS29, VPS26C, COMMD, CCDC22, CCDC93, CCC complex, Commander

## Abstract

The recycling of membrane proteins from endosomes to the cell surface is vital for cell signaling and survival. Retriever, a trimeric complex of VPS35L, VPS26C and VPS29, together with the CCC complex comprising CCDC22, CCDC93, and COMMD proteins, plays a crucial role in this process. The precise mechanisms underlying Retriever assembly and its interaction with CCC have remained elusive. Here, we present the first high-resolution structure of Retriever determined using cryogenic electron microscopy. The structure reveals a unique assembly mechanism, distinguishing it from its remotely related paralog, Retromer. By combining AlphaFold predictions and biochemical, cellular, and proteomic analyses, we further elucidate the structural organization of the entire Retriever-CCC complex and uncover how cancer-associated mutations disrupt complex formation and impair membrane protein homeostasis. These findings provide a fundamental framework for understanding the biological and pathological implications associated with Retriever-CCC-mediated endosomal recycling.

## Main

The plasma membrane (PM) proteome comprises a diverse array of components, including receptors, transporters, channels, and numerous other factors, which altogether account for nearly 11% of all proteins present in human cells^1, 2^. With the constant flux of membranes to and from the cell surface, PM proteins are frequently internalized into the endosomal system. Subsequently, these proteins require either recycling back to the cell surface for reuse or routing to lysosomes for degradation. Given the crucial roles of many PM proteins in cellular functions, the process of endosomal recycling is vital for maintaining cellular homeostasis. Multiple pathways exist in the cell to ensure the proper regulation of this recycling process.

One ancient pathway conserved from humans to yeast relies on a trimeric complex of ∼150 kDa known as Retromer, which is composed of VPS35, VPS26A/B, and VPS29^3–5^. Retromer facilitates the recycling of cargo proteins by directly binding to the cytoplasmic tails of certain receptors, including CI-M6PR and DMT1, or by interacting with various sorting nexin proteins, including SNX1, SNX2, SNX5, SNX6, SNX3, and SNX27. These sorting nexins in turn recruit various cargo proteins such as the copper transporters ATP7A and ATP7B, the glucose transporter GLUT1, and others^6–9^. Aside from cargo recognition, Retromer is essential for recruiting several cytosolic factors required for endosomal recycling. Among these is the WASH complex, a heteropentameric protein assembly containing a member of the Wiskott-Aldrich Syndrome Protein family named WASH, which stimulates the Arp2/3 complex to polymerize actin at the cytosolic side of endosomal membranes^10–13^. Without WASH-dependent actin polymerization, cargoes are trapped in the endosomal compartment, leading to their default trafficking to lysosomes for degradation.

Another crucial component involved in endosomal recycling is the COMMD/CCDC22/CCDC93 complex^14^ (CCC), a multiprotein assembly that has co-evolved with the WASH complex. The CCC complex was identified through proteomics as an assembly containing members of the COMMD protein family^15^, ranging from COMMD1 to COMMD10, together with two coiled-coil containing proteins, CCDC22 and CCDC93, which share distant homology with the kinetochore proteins Nuf2 and Ndc80^16^. The founding member of the COMMD family of proteins, COMMD1, contains an area of homology at its C-terminus, termed the COMM domain which mediates homo and heterodimer formation^15, 17^, and an N-terminal globular domain of tightly packed helices^18^. COMMD1 was initially discovered for its role in copper homeostasis in mammals^19, 20^. This function was soon linked to the role of CCC in regulating endosomal recycling of copper transporters ATP7A and ATP7B^14, 21^. Subsequent studies have revealed the involvement of CCC in the recycling of numerous other PM proteins, including those regulated by Retromer (such as Glut1 and CI-MPR) and proteins independent of Retromer (such as LDLR and Notch2)^22–24^. Defects in recycling have been observed in cultured cells, mouse models of CCC deficiency^25^, and patients carrying *CCDC22* mutations^22, 24^. Through interactome analysis, FAM45A (now known as DENND10) was identified as a significant CCC-interacting protein^22^. DENND10 contains a DENN domain characteristic of a family of Rab guanine exchange factors (GEF), which activate Rab GTPases crucial for endomembrane trafficking^26, 27^. While DENND10 appears to be tightly associated with CCC, its precise function, potential Rab targets, and contributions to the endosomal sorting process remain to be fully defined^22, 26^.

Unbiased proteomic screenings of CCC-interacting proteins and SNX17-associated factors also led to the discovery of another crucial component for endosomal recycling called Retriever^23^. Retriever is a trimeric complex of ∼160 kDa exhibiting a distant relationship with Retromer. Both Retriever and Retromer contain VPS29, while the other two subunits of Retriever, VPS35L and VPS26C, share less than 25% sequence identity with the corresponding subunits of Retromer, VPS35 and VPS26A/B, respectively. Similar to Retromer, Retriever is responsible for endosomal recycling of myriad client proteins, including integrins, members of the LDL receptor family, and likely numerous others^23^. Furthermore, recycling events involving Retriever are also dependent on proper activation of the WASH complex, which when dysregulated, prevents the recycling of LDL receptor family members and other Retriever cargoes^22, 24^.

The precise mechanisms underlying the assembly of Retriever and its regulation of endosomal recycling are still largely unknown. Proteomic analyses indicate that the CCC and Retriever subunits are closely linked^22^. Other studies examining protein assemblies across distinct organisms confirmed that CCC subunits (CCDC22, CCDC93 and 10 COMMD proteins) form a large conserved protein assembly as had been originally described^14^ and renamed the assembly as Commander^28^. Subsequently, through review of prior studies examining protein-protein interaction data, the same research group proposed that CCC and Retriever may form a larger assembly and utilized the term “Commander” to describe this proposed model^29^. At the same time, blue native gel electrophoreses have indicated that Retriever, defined as the trimer of VPS35L, VPS26C, and VPS29, migrates as an assembly with an apparent mass of ∼240 kDa, much like Retromer. On the other hand, CCC migrates with an apparent mass of ∼700-720 kDa. When the CCC complex fails to assemble in *COMMD3* deficient cells, the Retriever complex migrating at ∼240 kDa remains unaffected^22^. Therefore, while Retriever and CCC are closely linked molecular entities, the exact nature of their interaction and whether they function as one entity or as dynamically regulated individual entities in specific aspects of endosomal regulation are still unclear.

In this study we report the first high-resolution structure of Retriever determined by cryogenic electron microscopy (cryo-EM) and present an overall architecture of the Retriever-CCC complex derived from a combination of computational, biochemical, cellular, and proteomic approaches. These studies provide a comprehensive foundation to understand how these essential regulatory proteins work together to carry out their functions.

## Results

### Cryo-EM structure of Retriever reveals an assembly mechanism distinct from Retromer

To determine the structure of human Retriever, we co-expressed its three subunits, VPS29, VPS26C and VPS35L, in Sf9 cells using individual baculoviruses, in which only VPS29 contained a His_6_ tag at its C-terminus (CT) to facilitate purification. The purified Retriever eluted as a single peak in size-exclusion chromatography (Extended Data Fig. 1A), which gave rise to high-quality cryo-EM grids with homogeneous single particle distributions (Extended Data Fig. 1B). We then determined the cryo-EM structure of Retriever to a resolution of 2.9 Å using single particle reconstruction (Table 1). We applied local refinement to further improve the map quality in the region encompassing VPS29 and its interaction with VPS35L (Fig. 1A; Extended Data Fig. 1C-G).

**Table 1.** Cryo-EM data collection, refinement, and validation statistics.

|  | VPS35L-VPS29-VPS26C<br>(EMDB: 40885)<br>(PDB: 8SYN) | VPS35L(partial)-<br>VPS29<br>(EMDB: 40884)<br>(PDB: 8SYM) | Composite Map<br>(EMD: 40886)<br>(PDB: 8SYO) |
| --- | --- | --- | --- |
| **Data collection and processing** |  |  |  |
| Magnification | 105,000 |  |  |
| Voltage (kV) | 300 |  |  |
| Electron exposure (e<sup>-</sup>/Å<sup>-2</sup>) | 60 |  |  |
| Defocus range (μm) | -1.2 to -2.4 |  |  |
| Pixel size (Å) | 0.83 |  |  |
| Symmetry imposed | C1 | C1 |  |
| Initial particle images (no.) | 1,221,095 | 426,624 |  |
| Final particle images (no.) | 426,624 | 83,654 |  |
| Map resolution (Å) | 2.9 | 3.2 |  |
| FSC threshold | 0.143 | 0.143 |  |
| Map pixel size (Å) | 1.0624 | 1.0624 |  |
| **Refinement** |  |  |  |
| Initial model used (PDB code) | - | - |  |
| **Model composition** |  |  |  |
| Nonhydrogen atoms | 10,070 | 4,487 | 10,070 |
| Protein residues | 1,259 | 560 | 1,259 |
| Ligands | 0 | 0 | 0 |
| **R.m.s. deviations** |  |  |  |
| Bond lengths (Å) | 0.005 | 0.004 | 0.005 |
| Bond angles (°) | 0.550 | 0.454 | 0.550 |
| **Validation** |  |  |  |
| MolProbity score | 1.66 | 1.45 | 1.66 |
| Clashscore | 7.56 | 8.24 | 7.56 |
| Poor rotamers (%) | 0 | 0 | 0 |
| **Ramachandran plot** |  |  |  |
| Favored (%) | 96.4 | 98.0 | 96.4 |
| Allowed (%) | 3.6 | 2.0 | 3.6 |
| Disallowed (%) | 0 | 0 | 0 |
| Protein residues included in the model | VPS35L: 3-37, 110-139, 175-254, 268-924<br>VPS29: 3-186<br>VPS26C: 3-29, 38-56, 61-81, 86-127, 132-222, 225-297 | VPS35L: 3-37, 580-602, 607-924<br>VPS29: 3-186 | VPS35L: 3-37, 110-139, 175-254, 268-924<br>VPS29: 3-186<br>VPS26C: 3-29, 38-56, 61-81, 86-127, 132-222, 225-297 |

**Fig. 1.**
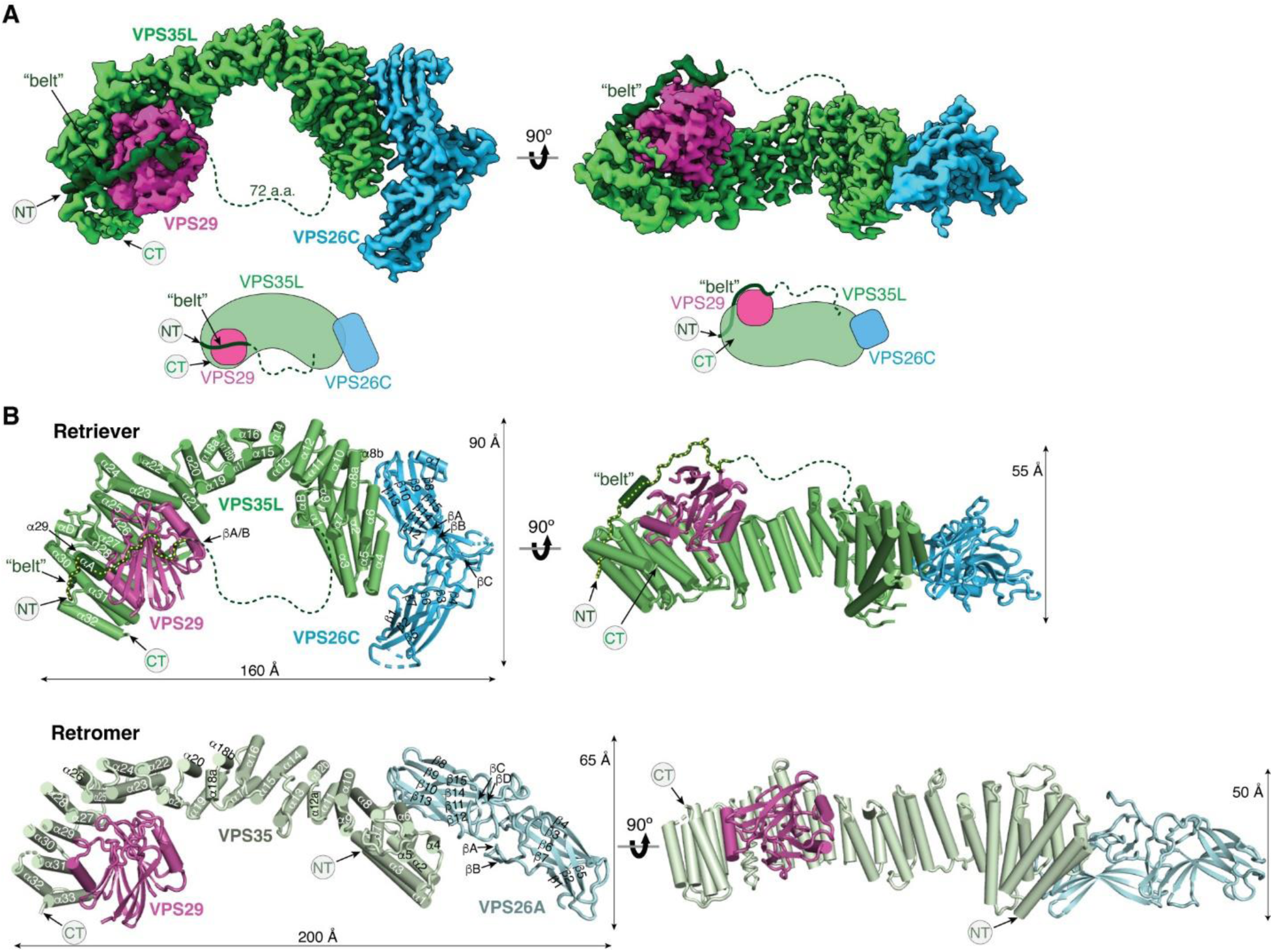
Cryo-EM structure of Retriever reveals a unique assembly mechanism. **(A)** Cryo-EM map (EMD: 40886; PDB: 8SYO) and schematic of the Retriever complex. Dotted lines represent the putative flexible linker sequence in VPS35L not observed in the map. **(B)** Structural comparison between Retriever (top) and Retromer (bottom, PDB: 7U6F). Secondary structural elements of the remotely homologous proteins, including VPS35L vs. VPS35 and VPS26C vs. VPS26A, are labeled. The “belt” sequence unique to VPS35L is traced by yellow dotted lines.

The overall structure of Retriever assumes a semicircular configuration measuring ∼55 x 90 x 160 Å, which is dominated by the extended conformation of VPS35L adopting an α/α solenoid fold that comprises 32 α−helices. Within the complex, the globular structure of VPS29 is partially embraced within an extensive pocket formed by the CT region of VPS35L, while the VPS26C binds to the outer ridge of the solenoid at the opposite end of VPS35L (Fig. 1A, B). Another key feature of the complex is the first 37 amino acids (a.a.) at the N-terminus (NT) of VPS35L. This NT peptide, hereafter referred to as the “belt” sequence due to its resemblance to a seatbelt, wraps around the CT region of VPS35L and the bound VPS29 (Fig. 1, dark green). Following the NT “belt” is a long, unstructured peptide linker of ∼72 residues, which extends toward the opposite end of the complex (Fig. 1A, B, dashed green line).

In many aspects, Retriever exhibits similar, yet distinct structural features compared to Retromer (Fig. 1B; Extended Data Fig. 2). The overall conformation of Retriever is more compact and twisted than Retromer, shorter by ∼40 Å in its longest dimension (Fig. 1B). Moreover, the overall molecular surface of Retriever is less negatively charged than Retromer (Extended Data Fig. 2A). Although VPS35L and VPS26C in Retriever share only ∼15% and 23/24% sequence identity with VPS35 and VPS26A/B in Retromer, respectively, their secondary structures exhibit remarkable similarities. Both VPS35L and VPS35 adopt the α/α solenoid fold with a similar number and organization of helices (Fig. 1B; Extended Data Fig. 2B). Both VPS26C and VPS26A consist of two domains formed by a similar number and arrangement of β-strands, which pack into a deeply curved β-sandwich resembling an arrestin fold (Fig. 1B; Extended Data Fig. 2B). Despite these similarities, however, VPS35L exhibits a more compact structure than VPS35 and contains several unique features absent from VPS35. These include the NT “belt” peptide, which makes extensive contacts with VPS29 and the CT region of VPS35L, and several additional short helices and a β-hairpin, which are inserted between the common solenoid helices (Fig. 1B). Similarly, VPS26C is also more compact than VPS26A and contains several distinct short β-strand insertions compared to VPS26A (Fig. 1B; Extended Data Fig. 2B). In contrast, the VPS29 subunit, which adopts a globular, phosphodiesterase/nuclease-like fold, has nearly identical structures in both Retriever and Retromer, with a root mean square deviation (RMSD) of ∼1 Å for all Cα atoms between the two complexes (Extended Data Fig. 2B).

**Fig. 2.**
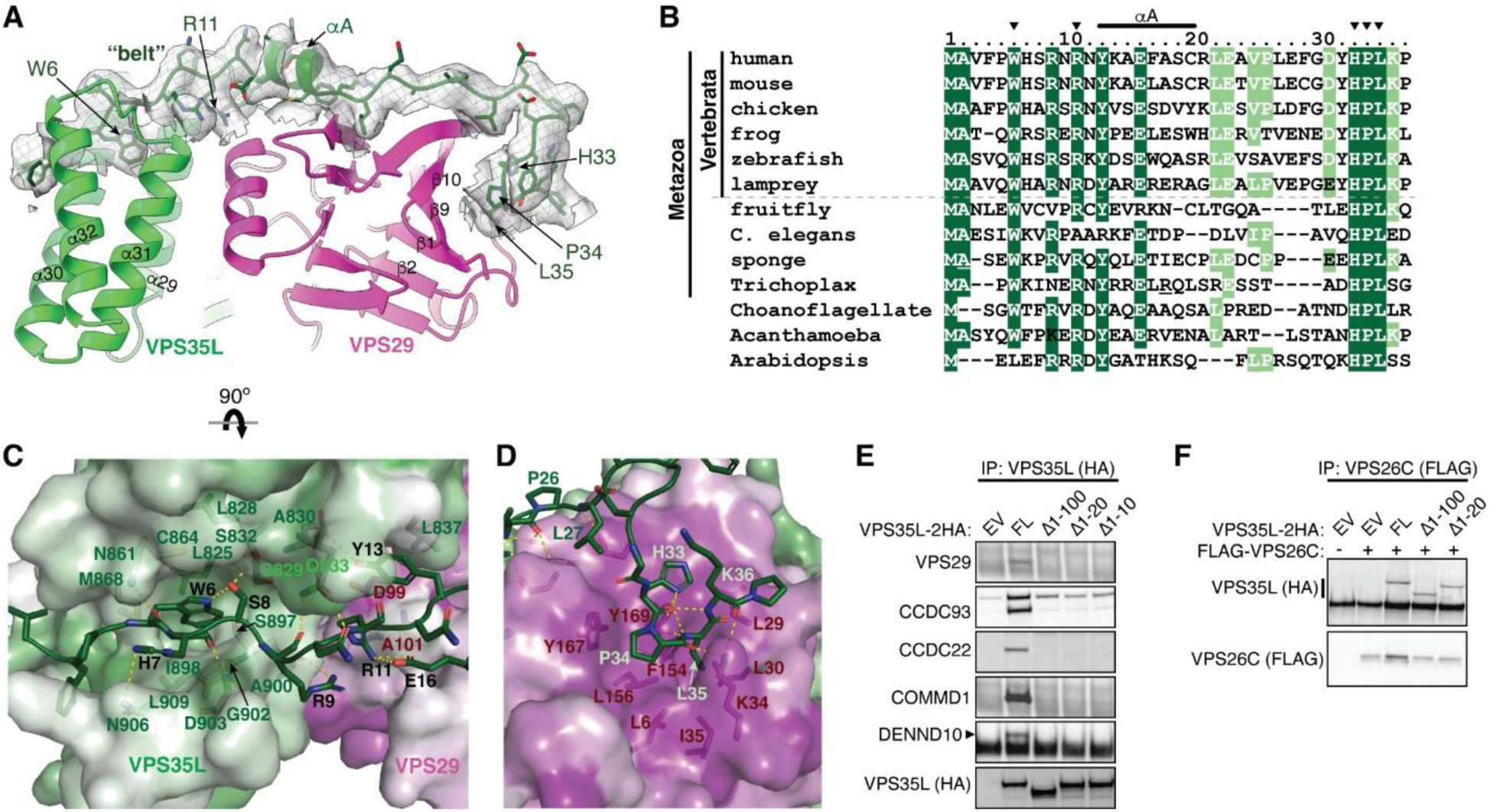
The N-terminal “belt” sequence unique to VPS35L is key to Retriever assembly. **(A)** Cryo-EM density of the “belt” sequence interacting with VPS35L and VPS29. **(B)** Alignment of the “belt” sequences from representative species from animal to amoeba and plants. Residues shown in (A) are marked with arrowheads. **(C-D)** Key interactions between the “belt” sequence (represented in cartoon, with carbon in green, oxygen in red, and nitrogen in blue) and its binding surface on VPS35L (C) and VPS29 (D). The binding surface is colored based on conservation score calculated by Consurf^71^, with color to white gradients representing the most (ConSurf score = 9) to the least conserved residues (ConSurf score = 1). Contacting residues are shown as sticks. Yellow dashed lines indicate polar interactions. **(E-F)** Immunoprecipitation of VPS35L NT-deletion mutants expressed in HEK293T cells. Interactions with indicated components of Retriever and CCC were assessed by immunoblotting.

The interface between VPS35L and VPS29 in Retriever buries a solvent accessible area of ∼2,400 Å^2^, which is significantly larger than the interface between VPS35 and VPS29 (∼1,400 Å^2^) in Retromer (Extended Data Fig. 2C). This difference is largely contributed by the interface between VPS29 and the NT “belt” peptide of VPS35L, which alone accounts for a buried solvent accessible area of ∼700 Å^2^. Even when considering only the interface between the CT of VPS35L and VPS29, it is still ∼20% larger (∼1,700 Å^2^) than the corresponding interface in Retromer, likely due to the more compact architecture adopted by VPS35L (Extended Data Fig. 2B, C). Moreover, the interface between VPS35L and VPS26C in Retriever (∼1,000 Å^2^) is ∼50% larger than the interface between VPS35 and VPS26A in Retromer (∼670 Å^2^). This further underscores the more compact nature of the Retriever complex (Extended Data Fig. 2D).

### The NT “belt” sequence of VPS35L plays a key role in stabilizing Retriever

Given the distinctive feature of the NT “belt” sequence in VPS35L and its extensive contact with both the CT region of VPS35L and the bound VPS29, we asked whether the “belt” sequence could play an important role in stabilizing Retriever assembly. Close inspection of the structure reveals two major anchoring points in the “belt” sequence (Fig. 2A). First, the NT 11 residues of the “belt” sequence winds through a deep trough on the CT region of VPS35L formed by the ends of helix α29, α30, α31, and α32, largely through structural complementarity (Fig. 2A, C). Consequently, this interaction makes the “belt” sequence an integral part of the CT region of VPS35L. The interaction is centered around W6, a highly conserved residue in orthologs ranging from amoeba to humans (Fig. 2B). W6 inserts into a deep pocket formed by L825, L828, S829, C864, M868, I898, G902, and L909 from VPS35L (Fig. 2C). This interface is stabilized by extensive *van der Waals* interactions and a few hydrogen bonds. At the boundary of this extensive interaction surface, the conserved residue R11 of the “belt” sequence is supported by salt bridges with two conserved residues, E16 of the “belt” sequence itself and D99 from the bound VPS29 (Fig. 2B, C).

The second conserved anchoring point of the “belt” sequence is located at its C-terminus, where it interacts with VPS29 largely through a conserved “HPL” motif in Retriever (Fig. 2C-D). This interaction is unique to Retriever and absent between VPS29 and VPS35 in Retromer (Fig. 1B). It is remarkable that the HPL motif is virtually 100% conserved in all examined organisms (Fig. 2B). The motif adopts a type-I β-turn structure through a network of intrapeptide hydrogen bonds (Fig. 2D). At the tip of the β-turn, P34 and L35 of the “HPL” motif insert into a conserved and largely hydrophobic pocket on VPS29 formed by β1, β9, β10 and the linker connecting α1 and β2, consisting of L6, L29, L30, K34, I35, F154, L156, Y167, and Y169 (Fig. 2D, 3A). This interaction is further stabilized by a hydrogen bond network involving K34 and Y169 from VPS29 and H33, P34, and L35 from VPS35L (Fig. 2D).

Consistent with the observation that the belt is an integral component of the CT region of VPS35L and essential for VPS29 binding, deleting the first 10 amino acids of VPS35L was sufficient to abrogate VPS35L-VPS29 interaction in cells, as noted in co-immunoprecipitation (co-IP) experiments (Fig. 2E). In contrast, complete deletion of the “belt” sequence or even the first 100 amino acids, which include the unstructured linker sequence, had no effect on the binding between VPS35L and VPS26C (Fig. 2F), in agreement with the presented Retriever structure. Surprisingly, disrupting the interaction between VPS29 and VPS35L eliminated the interaction between VPS35L and the CCC subunits CCDC22, CCDC93, and COMMD1, as well as DENND10 (Fig. 2E). This suggests an interdependence between VPS29-VPS35L and Retriever-CCC interactions, as will be examined in later parts of the paper.

### Conserved surfaces in VPS35L that bind to VPS26C and VPS29 are mutated in cancer

In addition to the NT “belt” sequence contacting VPS29, the CT region of VPS35L interacts with VPS29 using a slightly concave and extensive surface (Fig. 3A, Extended Data Fig. 2C). This interface between VPS35L and VPS29 in Retriever is analogous to the binding interface between VPS35 and VPS29 in Retromer. On VPS29, it involves the four extended loops connecting β1 and α1 (herein referred to as Loop1), β3 and β4 (Loop2), β5 and α3 (Loop3), and β8 and β9 (Loop4) (Fig. 3A, B). On VPS35L, it involves the solenoid helices α21, α23, α25, α27, α29, and α31, as well as the β-hairpin inserted between α26 and α27 (Fig. 3A, B). Many residues in the CT region of VPS35L establish extensive polar and non-polar contacts with VPS29 through this broad interaction surface (Fig. 3B).

**Fig. 3.**
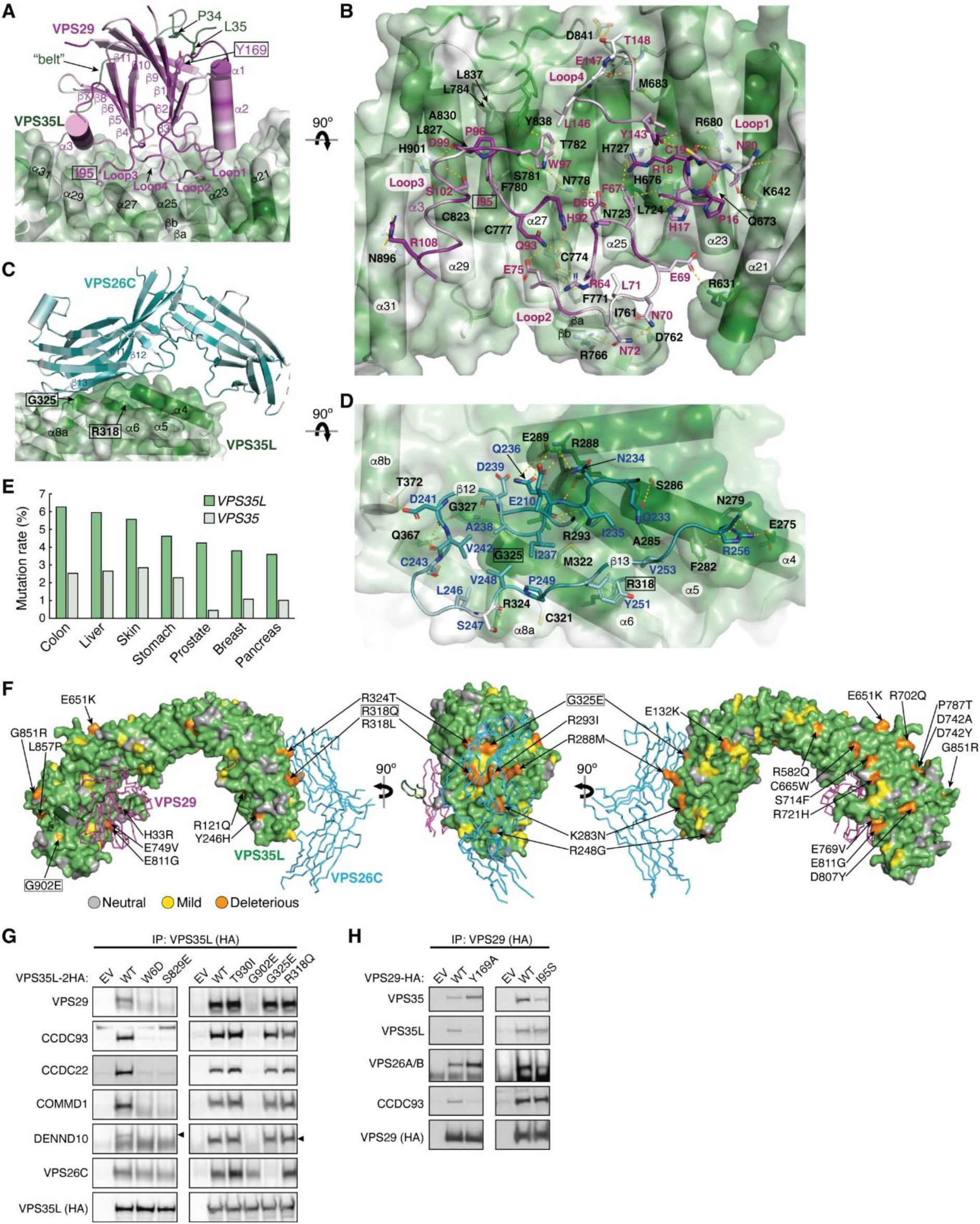
VPS35L bridges VPS26C and VPS29 through conserved surfaces. **(A-D)** Interaction surface of VPS35L with VPS29 (A-B) and VPS26C (C-D). The binding surface is colored based on conservation score using the same scheme shown in Fig. 2. Contacting residues are shown as sticks. Yellow dashed lines indicate polar interactions. For clarity, the backbones of VPS29 and VPS26C in (B) and (D) are shown as loops. **(E)** Mutation rates (%) for *VPS35L* and *VPS35* across multiple tumor types. **(F)** Overall structural model of Retriever showing the location of cancer-associated mutations on the surface of VPS35L. Residues mutated in this study are outlined with a black box. For clarity, VPS29 and VPS26 are shown as ribbons. **(G-H)** Immunoprecipitation of VPS35L (G) or VPS29 (H) carrying indicated point mutations expressed in HEK293T cells. Interactions with various components of Retriever and CCC were assessed by immunoblotting.

The similarity between VPS29-VPS35L interface in Retriever and VPS29-VPS35 interface in Retromer poses a challenge in the design of a mutation that can specifically disrupt one interaction without affecting the other. To specifically disrupt VPS29-VPS35L interaction in Retriever, instead of mutating this extensive surface, we introduced Y169A to VPS29 to disrupt the interaction between VPS29 and the “HPL” motif in the “belt” sequence (Fig. 3A). Y169 is located at the base of the hydrophobic pocket forming hydrogen bonds and a π-π interaction with the “HPL” β-hairpin (Fig. 2D). As expected, Y169A significantly decreased the binding of VPS29 to VPS35L (Fig. 3H). Interestingly, this mutation simultaneously increased the binding to Retromer components VPS35 and VPS26A/B (Fig. 3H), suggesting a potential competition between Retriever and Retromer for the same pool of VPS29 in cells. Next, we tested the effect of a mutation in VPS29, I95S, which was previously shown to disrupt the VPS29-VPS35 interaction in Retromer^30^. Interestingly, although I95 in VPS29 has a close contact with both VPS35 and VPS35L, this mutation selectively reduced the binding to VPS35, but preserved the interaction with VPS35L (Fig. 3H). This result highlights the differences in the binding mechanism of VPS29 between Retromer and Retriever.

The interaction between VPS35L and VPS26C is mediated by an extensive and conserved interface involving β12 and β13 of VPS26C and α4, α5, α6, and α8 of VPS35L (Fig. 3C-D, Extended Data Fig. 2D). A similar set of β-strands and solenoid helices are also involved in the VPS26A-VPS35 interaction in Retromer (Extended Data Fig. 2D). The interaction surface comprises a largely hydrophobic core region surrounded by a series of polar interactions in the periphery involving E210, Q233, N234, Q236, D241, and R256 in VPS26C and E275, N279, S286, R288, E289, R293, Q367, and T372 in VPS35L (Fig. 3D, Extended Data Fig. 2D).

Previous studies reported that the rate of mutation in VPS35L exceeds random mutation burden when the gene length is considered^31^. Our review of the COSMIC database (https://cancer.sanger.ac.uk/cosmic) also indicates that the rate of somatic mutations in VPS35L exceeds that of its closest paralog, VPS35, across all tumor types (Fig. 3E). We first used the SNAP2 tool to assess the potential impact of the missense mutations^32^, through which we identified a number of somatic mutations with high likelihood of functional impairment, accounting for 25 – 52% of total 235 missense mutations, depending on the evaluation stringency (Extended Data Table 2). When projected onto the cryo-EM structure of Retriever, several of these mutations were found to potentially disrupt the interaction between the NT “belt” and the CT region of VPS35L, while others were clustered over the binding surfaces for VPS29 and VPS26C (Fig. 3F).

We then selected several mutations, including a few derived from our structural analyses, to test how they may impact Retriever assembly. We found that mutations predicted to disrupt the interaction between the NT “belt” and the CT region of VPS35L, including W6D, S829E, and the cancer-derived mutation G902E, abolished the binding to VPS29 without affecting VPS26C binding (Fig. 3G). In addition, these mutations simultaneously disrupted the binding to CCC components, including CCDC93, CCDC22, and COMMD1, as well as the binding to DENND10. The same effects were observed when the “belt” sequence was deleted, as shown earlier (Fig. 2E, VPS35L D10). In contrast, the cancer-derived mutation G325E specifically disrupted VPS35L binding to VPS26C, without affecting the binding to VPS29 or CCC components (Fig. 3G). This suggests that, unlike VPS29, the association of VPS26C with VPS35L does not contribute to the Retriever-CCC interaction. Other mutations in the NT “belt” or the CT region of VPS35L did not exhibit appreciable effects on complex assembly under our experimental conditions, when they were transiently expressed and mutated in isolation (Fig. 3G, Extended Data Fig. 3D).

We proceeded with the four mutations that had profound effects on Retriever assembly to further examine how they may impair Retriever function in cells. For this, we used CRISPR/Cas9 mediated gene editing to knock out *VPS35L* from liver cancer Huh-7 cells and then stably reconstituted VPS35L expression using an empty vector (EV), or VPS35L variants, including wild-type (WT), W6D, S829E, G902E, and G325E (Extended Data Fig. 3A). The stable expression of VPS35L variants produced similar results obtained from the transient transfections shown in Fig. 3G, in which mutants of VPS35L affecting the NT “belt” interaction (W6D, S829E, and G902E) failed to bind VPS29 or CCC components in co-IP, while G325E specifically abrogated VPS26C binding while not affecting VPS29 or CCC binding (Extended Data Fig. 3A).

Using these stable cell lines, we purified VPS35L-associated complexes using HA tag-mediated immunoprecipitation. The native complexes were eluted by HA peptide competition in a non-denaturing, physiological buffer and then resolved through blue native gel electrophoresis. Consistent with previous findings^22^, we found that VPS35L WT partitioned in two distinct complexes: a smaller complex with an apparent Mw of ∼240 kDa corresponding to Retriever, and a larger complex with an apparent Mw of over 720 kDa, which contained CCC, as confirmed by immunoblotting for COMMD1 (Extended Data Fig. 3B). Interestingly, CCC has a unique band at ∼500 kDa devoid of VPS35L, suggesting that the interaction between Retriever and CCC is likely not constitutive, leading to their dissociation during electrophoresis (Extended Data Fig. 3B). In contrast to VPS35L WT, all mutations that abolished VPS29 binding (W6D, S829E, and G902E) failed to precipitate Retriever or CCC, while the mutation that disrupted VPS26C binding (G325E) resulted in the absence of the Retriever complex at ∼240 kDa, while still maintaining the binding to CCC (Extended Data Fig. 3B).

To further investigate the impact of the four mutations in VPS35L on protein-protein interactions, we conducted semiquantitative unbiased proteomics. Compared to the empty vector control, VPS35L WT bound most intensely to VPS26C and CCC complex components (with VPS29 binding not detectable above background signals in this approach), followed by weaker binding to several additional known interactors, such as WASHC5, and many previously unreported interactors (Extended Data Fig. 3C). In contrast, compared to VPS35L WT, the W6D, S829E, and G902E mutations were devoid of CCC binding while preserving the interaction with VPS26C. Conversely, G325E showed a complete loss of VPS26C binding, while preserving the interaction with CCC (Extended Data Fig. 3C). These proteomic results aligned well with the co-IP experiments (Fig. 3G, Extended Data Fig. 3A-B) and provided further validation for the cryo-EM structure of Retriever. It is worth noting that none of these mutations affected other interactions, such as the binding to WASHC5, a component of the WASH complex, suggesting that the mutations were specific and did not interfere with protein folding or disrupt other surfaces involved in additional protein-protein interactions.

### Disrupted Retriever assembly alters PM homeostasis

The specific effects of structure-guided and cancer-associated mutations in VPS35L allowed us to examine the physiological function of Retriever assembly in cell models. First, we observed that all VPS35L variants maintained endosomal localization, irrespective of their ability to interact with CCC, which is evident from their co-localization with the WASH subunit FAM21 (Extended Data Fig. 4A). We confirmed our prior observation that loss of the CCC complex, as a result of *COMMD3* or *CCDC93* deficiency, increased VPS35L cytosolic staining^22^, but did not completely abrogate endosomal recruitment of VPS35L (Extended Data Fig. 4B). Thus, Retriever recruitment to endosomes, while enhanced by CCC, is not fully dependent on it, thus explaining the similar localization of VPS35L mutants on FAM21-positive endosomes (Extended Data Fig. 4A). While VPS35L is predominantly endosomal, we observed that a small amount of the protein is detectable in LAMP1+ vesicles. Interestingly, mutants that lost the ability to bind to VPS29 and CCC (i.e., W6D, S829E and G902E) had reduced localization to this compartment, while the G325E mutation disrupting VPS26C binding had no significant effect (Fig. 4A, B).

**Fig. 4.**
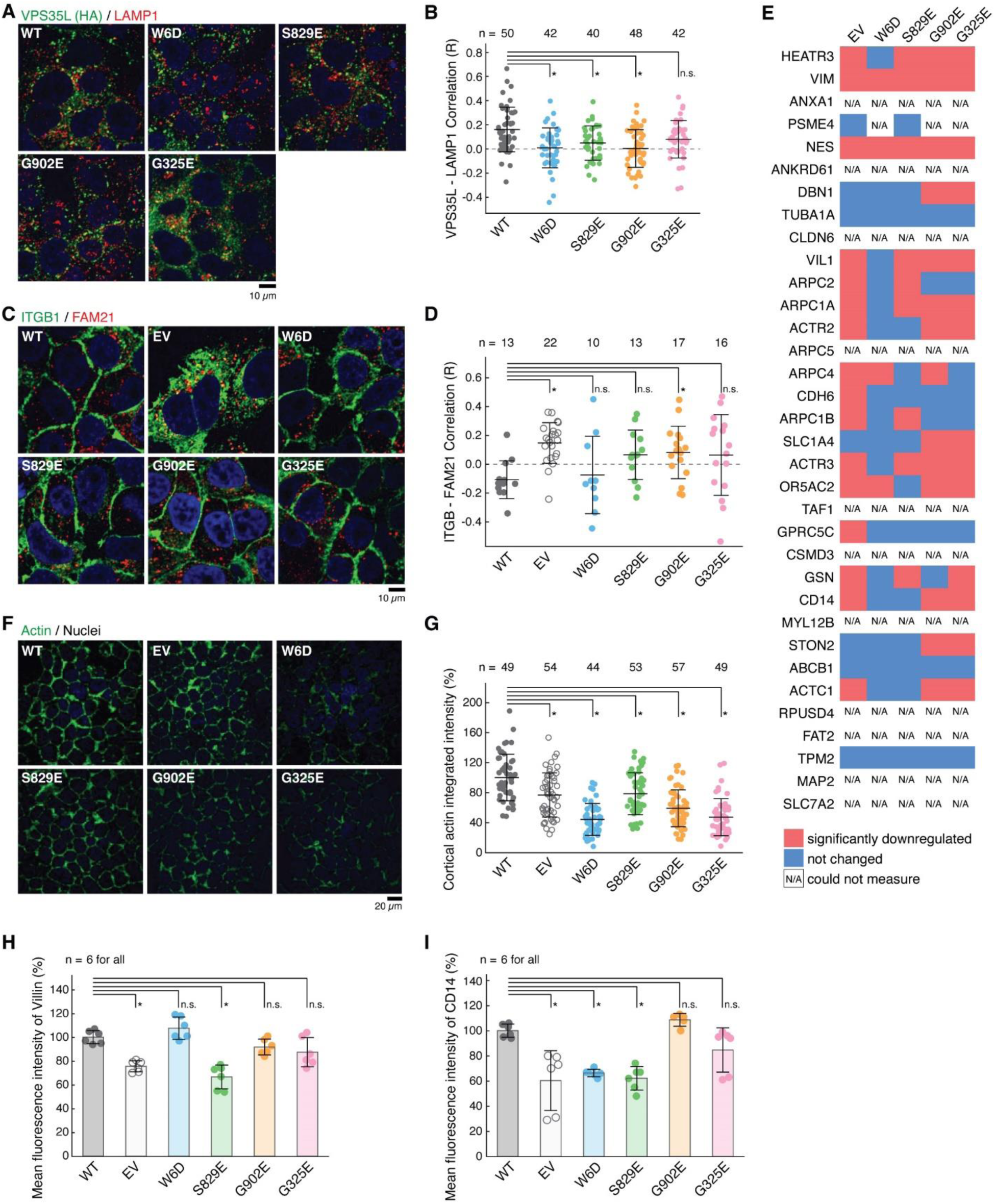
Disruption of Retriever assembly affects membrane protein homeostasis. **(A)** Immunofluorescence staining for VPS35L (green channel, using HA antibody), LAMP1 (red channel), and nuclei (DAPI, blue channel) in the indicated stable Huh-7 cell lines. **(B)** Quantification of the correlation coefficient for VPS35L and LAMP1 localization for the images shown in (A). Each dot represents an individual cell. **(C)** Immunofluorescence staining for ITGB1 (green channel), FAM21 (red channel), and nuclei (DAPI, blue channel) in the indicated stable Huh-7 cell lines. **(D)** Quantification of the correlation coefficient for ITGB1 and FAM21 localization for the images shown in (C). Each dot represents an individual cell. **(E)** Surface biotinylation and protein isolation, followed by proteomic quantification was performed and protein abundance was compared against VPS35L WT in the indicated cell lines stable Huh-7 cell lines. Red indicates values for proteins with at least 50% reduction compared to VPS35L WT cells, blue represents values that were not significantly reduced, while N/A represents proteins that could not be quantified. **(F)** Phalloidin staining for F-Actin (green channel) and nuclei (DAPI, blue channel) in the indicated stable Huh-7 cell lines. **(G)** Quantification of the cortical actin staining in the images shown in (F). Each dot represents an individual cell. **(H-I)** Quantification of Villin (H) and CD14 (I) fluorescence staining intensity as determined by FACS, expressed as % compared to VPS35L WT cells.

Next, we assessed the impact of disrupting Retriever assembly on the trafficking of a well-established cargo protein, Integrin-β1 (ITGB1). Loss of VPS35L is known to impair ITGB1 endosomal recycling^22, 23^, which was also observed in Huh-7 *VPS35L* knockout cells rescued by empty vector (EV), where we observed significant endosomal trapping of ITGB1 (Fig. 4C, D).

Compared to EV, however, the impact on ITGB1 recycling was not as profound for other mutants in VPS35L, with G902E showing significant endosomal trapping, while other mutants showing a milder and statistically insignificant effect (Fig. 4C, D). Thus, these data suggest that the mutations did not fully abrogate the function of Retriever.

To further delineate the functional effects of these mutations we used surface biotinylation followed by mass spectrometry analysis to examine the PM proteome in cells expressing various VPS35L mutants. First, when using tandem mass tagging (TMT)-based proteomics to compare isogenic *VPS35L* knockout cells re-expressing EV versus VPS35L WT, we identified 236 proteins showing significant changes in abundance within the biotinylated pool (p < 0.05 and fold change greater than 1.5 or lower than 0.65). When more stringent abundance cutoff values were applied (greater than 2-fold or lower than 0.5-fold), 67 proteins showed statistically significant changes in abundance. When we repeated the quantification using spectral counts instead of TMT, 23 of 34 proteins with 2-fold reduced PM expression in the EV condition showed similarly reduced surface expression (Fig. 4E). Between the two quantification methods, the largest discrepancies were accounted for by low expression and absent quantification when using spectral counts for quantification. It is remarkable that various VPS35L mutants, particularly the cancer-associated ones (G902E and G325E), recapitulated many of the changes caused by the deletion of *VPS35L* (EV column) (Fig. 4E).

Among these proteins were various membrane anchored proteins (e.g., CD14, SLC7A2) as well as membrane proximal proteins (e.g., ACTR1, ACTR2, ARPC1A, ARPC2, ARPC4). Prominent in the latter group were several components of the Arp2/3 complex. It was previously shown that Arp2/3 is more extensively recruited to endosomes in *CCC* and *VPS35L* deficient cells^22^. This is consistent with our observations here that Arp2/3 was correspondingly reduced from the PM (Fig. 4E). In agreement with the proteomic findings, we observed significant reduction measurement in the cortical actin compartment after knockout of *VPS35L* (EV) and in all the VPS35L mutants, (Fig. 4F, G). Moreover, VIL1 (Villin1), an actin binding protein localized to the brush border of epithelial cells, also displayed decreased expression in mutant cells by flow cytometry analysis, correlating well with the reduction in Arp2/3 seen by proteomics and with the reduction in cortical actin (Fig. 4E-H). Finally, surface levels of CD14, another PM protein noted in our proteomic analysis, were also reduced in mutant cells (Fig. 4I). Altogether, these studies demonstrated that disruption in Retriever assembly has a profound impact in PM homeostasis.

### Structural model of Retriever – CCC complex formation

In the aforementioned results, it is intriguing that all mutations disrupting the VPS29-VPS35L interaction also led to the dissociation of Retriever from CCC and DENND10 (Fig. 3, Extended Data Fig. 3). To understand the structural mechanisms underlying this observation, we employed ΑlphaFold 2 multimer (AFM) to predict how Retriever and CCC interact with each other^33, 34^. Through extensive rounds of iterations where we systematically examined various combinations of subunits, we were able to obtain highly reliable models that depict the architecture of the entire Retriever-CCC complex. These models were further validated using our biochemical and cellular assays. In the following sections, we will describe the structural models in separate segments.

### The CCDC22-CCDC93 dimer binds to the outer ridge at the CT of VPS35L

We first evaluated the reliability of AFM predictions by examining its capability to predict the structure of Retriever itself, for which no homologous structures were yet available. Remarkably, all predicted models exhibited a near perfect alignment with our cryo-EM structure, with an average RMSD of ∼2 Å (Extended Data Fig. 5A-C). It is interesting that the variations among the predicted models mainly arose from the subtle differences in model compactness (Extended Data Fig. 5A). This phenomenon mirrors what we observed during cryo-EM structure determination, where the cryo-EM particles displayed some heterogeneity in compactness, and larger motions were observed near both ends of the elongated VPS35L, leading to reduced resolution in these areas (Extended Data Fig. 1C-G). Equally reliable AFM models were obtained for Retromer, which also aligned near perfectly with the published cryo-EM structure (not shown). Hence, AFM can reliably predict unknown structures of large complexes. In all our AFM predictions, we applied three criteria to evaluate the reliability of the predicted models^33, 34^. These included the predicted local difference distance test (pLDDT) scores to assess the accuracy of the local structure, the predicted aligned error (PAE) scores to evaluate the distance error between every pair of residues, and the visual consistency of at least 25 solutions when aligned to evaluate the convergence of predictions. In most cases, we found that the visual consistency of the 25 aligned models agreed well with the PAE and pLDDT criteria.

In all the AFM predictions involving different subunits of CCC and Retriever, only CCDC22 and CCDC93 always bound to VPS35L in a highly consistent manner, while none of the other subunits were able to establish a reliable contact between CCC and Retriever. In all solutions, CCDC22 and CCDC93 form an extended heterodimer containing four coiled coils. The last two and a half-coiled coils (CC2b, CC3, and CC4) at the C-termini were consistently predicted to interact with a conserved surface at the outer ridge of the CT region of VPS35L, with CC3 and CC4 adopting a sharp V-shaped configuration (Fig. 5A-B, Extended Fig. 5D-F). These prediction results were not affected by the presence or absence of other CCC or Retriever components (not shown). We hereafter refer to the CC2b, CC3, and CC4 regions as the VPS35L binding domain (VBD). In addition to the VBD, the small globular domain located at the N-terminus of CCDC22, known as the NDC80-NUF2 calponin homology (NN-CH) domain^16^, consistently occupies the space between the V-shaped CC3 and CC4, irrespective of the presence of VPS35L (Fig. 5A-B). The NN-CH domain does not directly contact VPS35L and likely plays a structural role in stabilizing the CC3-CC4 conformation.

**Fig. 5.**
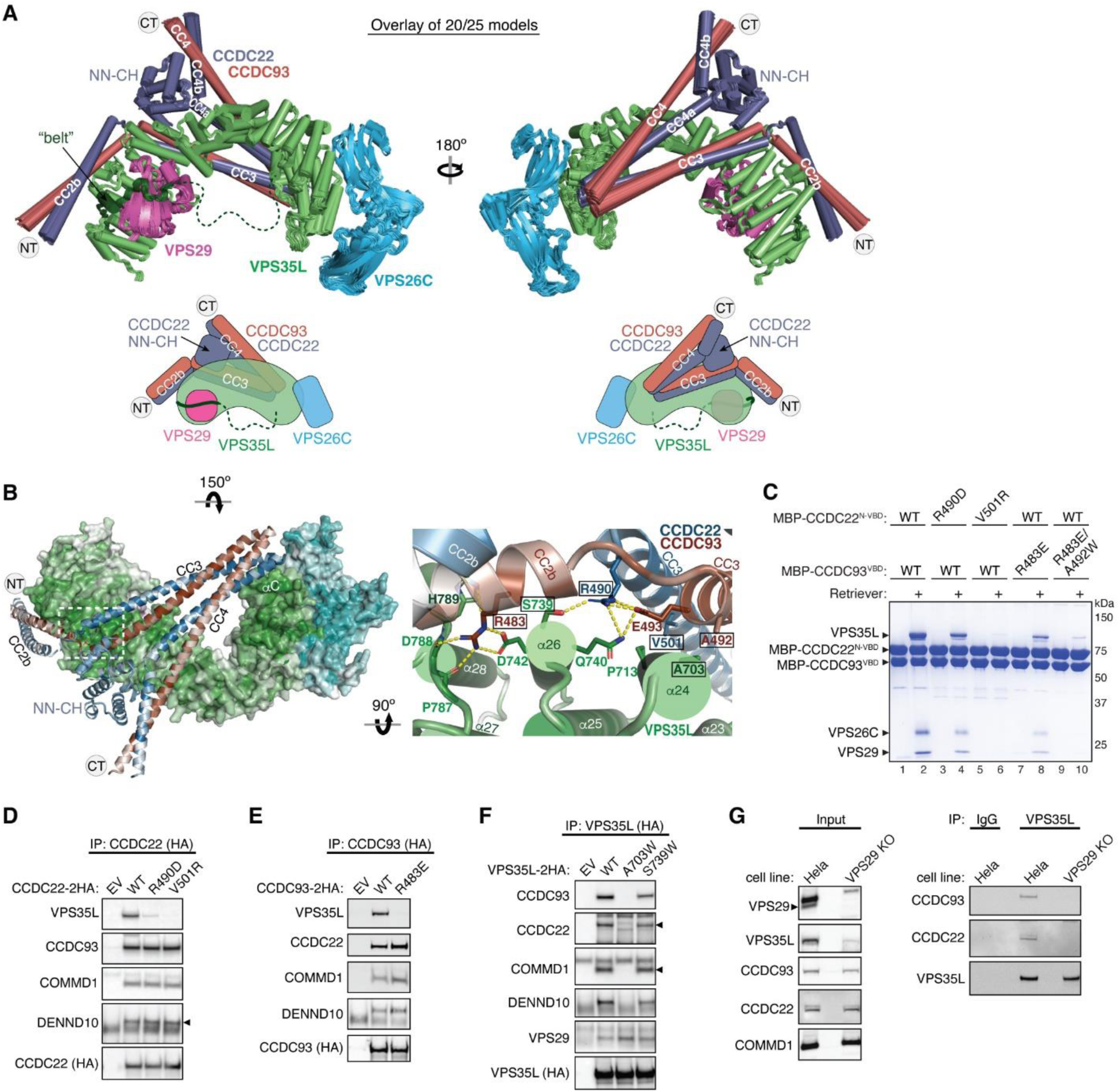
Structural model of CCDC22-CCDC93 binding to Retriever. **(A)** Overlay of AlphaFold Multimer models and schematic showing Retriever binding to CCDC22-CCDC93. For clarity, inconsistent models (5 out of 25 total models) are excluded. Unreliable structural regions showing inconsistency between models and high PAE scores are removed, including the peptide linker following the “belt” sequence in VPS35L (dotted green line). **(B)** Interaction surface between Retriever and CCDC22-CCDC93 colored by conservation score using the same scheme shown in Fig. 2. Key interactions are shown as sticks and polar interactions are represented with a dashed yellow line. Residues mutated in this study are outlined with a black box. **(C)** Coomassie blue-stained SDS PAGE gel showing indicated variants of MBP-CCDC22 NN-CH-VBD/MBP-CCDC93 VBD dimers (200 pmol) pulling down Retriever (60 pmol). **(D-F)** Immunoprecipitation of indicated mutants of CCDC22 (D), CCDC93 (E), and VPS35L (F) expressed in HEK293T cells and immunoblotting of indicated proteins. **(G)** Immunoprecipitation and immunoblotting of VPS35L from parental HeLa cells and a *VPS29* knockout line derived from these cells.

The VBD interacts with VPS35L at two conserved surfaces. The first surface encompasses helix α24 and the connecting loops between α25 and α26, α27 and α28, and α29 and α30 (Fig. 5B). The second surface is contributed by helix αC, which precedes the solenoid helix α1 (Fig. 5B). This αC helix is absent in VPS35 and is not visible in our cryo-EM map of Retriever. Interestingly, the first VBD binding surface is located at the opposite side of the same solenoid region of VPS35L that binds to VPS29. In addition, the coiled coil CC2b is in close proximity to the “belt” peptide (Fig. 5B). This configuration provides a plausible explanation for why disrupting the “belt” peptide or VPS29 binding impacted the binding to CCC (Fig. 3, Extended Data Fig. 3). We propose that loss of the “belt” peptide or VPS29 disturbs the local conformation of the CT region of VPS35L, which in turn allosterically destabilizes the binding of CCDC22-CCDC93 binding to Retriever.

To validate the predicted model, we purified the recombinant CCDC22-CCDC93 VBD dimer tagged with MBP (maltose binding protein) and used an MBP pull-down assay to test if the VBD directly interacts with purified Retriever. Given the apparent importance of the NN-CH domain of CCDC22 for the VBD structure, we introduced a flexible linker, (GGSK)_6_, to connect the NN-CH domain to the N-terminus of CCDC22 CC2b helix. The linker length used was of adequate to connect the C-terminus of the NN-CH domain to the N-terminus of CCDC22 CC2b helix if the AFM model is correct.

Consistent with the predicted model, the MBP-tagged VBD dimer robustly retained Retriever in pull-down assays (Fig. 5C, lane 2). To test if the interaction was specific, we introduced mutations in conserved residues that were predicted to be crucial for binding. These residues include R490 in CCDC22, which forms hydrogen bonds with S739 and Q740 in VPS35L, V501 in CCDC22, which interacts with a hydrophobic surface near P713 on α24 of VPS35L, R483 in CCDC93, which engages in a network of polar and π-π interactions with D742, P787, D788, and H789 on VPS35L, and A492 in CCDC93, which forms a close hydrophobic contact with A703 on α24 of VPS35L (Fig. 5B). All of the mutations impacted the interaction between the purified VBD and Retriever, although to different extents (Fig. 5C). R490D in CCDC22 and R483E in CCDC93 partially impaired the binding, while V501R in CCDC22 and the R483E/A492W double mutation in CCDC93 completely abolished Retriever binding (Fig. 5C).

Consistent with the in vitro pull-down results, when CCDC22 or CCDC93 bearing the same mutations were expressed in mammalian cells, they lost the interaction with VPS35L in co-immunoprecipitation experiments, but still maintained the interaction with other CCC components and DENND10 (Fig. 5D-E). Conversely, reciprocal mutations in VPS35L, including A703W and S739W, also affected the interaction. A703W completely abolished the binding to CCC complex and DENND10 (but not VPS29), while S739W exhibited a milder effect (Fig. 5F). Together, the above results provide strong validation of the AFM predicted model depicting how CCDC22-CCDC93 interacts with VPS35L.

Finally, we tested if VPS35L alone is sufficient for binding to CCDC22-CCDC93 VBD or if the presence of VPS29 is necessary for the interaction. Our earlier results demonstrated that mutations in VPS35L impacting VPS29 binding also affected the binding to CCC (Fig. 3, Extended Data Fig. 3), suggesting that VPS29 plays a role in VPS35L binding to CCDC22-CCDC93. To test this hypothesis, we used CRISPR/Cas9 to knock out *VPS29* and asked if the remaining VPS35L could immunoprecipitate CCC. The loss of VPS29 expression led to reduced expression of VPS35L, but did not affect the levels of CCC components, including CCDC93, CCDC22, and COMMD1 (Fig. 5G, left). Consistent with our mutagenesis data, VPS35L alone could not bind to CCC in VPS29 knockout cells (Fig. 5G, right), confirming that VPS29 is indeed necessary for the interaction between VPS35L and CCC. As VPS29 is not expected to have a direct contact with CCDC22-CCDC93, this result further supports our hypothesis that VPS29 facilitates a favorable conformation of VPS35L that is required for the interaction with CCC; mutations that disrupts VPS29 binding to VPS35L destabilizes this conformation, leading to the loss of CCC interaction.

### DENND10 binds directly to the CCDC22-CCDC93 dimer

During our AFM predictions, we included DENND10 to explore its relationship with the overall assembly of Retriever-CCC. We found that all AFM predictions involving CCDC22, CCDC93, and DENND10 consistently yielded a highly coherent model showing that DENND10 binds specifically to the CC1 and CC2a coiled coils of the CCDC22-CCDC93 heterodimer (Fig. 6A, Extended Data Fig. 6A-C), but not to any other components of the CCC or Retriever (not shown). This result suggests that the interaction between Retriever and DENND10 is indirect and is mediated by CCDC22-CCDC93. This model explains our experimental observations. First, Retriever could only co-immunoprecipitate DENND10 whenever CCC was also co-precipitated (Fig. 3G). Second, mutations in VPS35L specifically disrupting the binding to CCDC22-CCDC93 VBD similarly affected the binding to DENND10 (Fig. 5F). Third, mutations in CCDC22-CCDC93 VBD specifically disrupted the binding to VPS35L without affecting the interaction with DENND10 (Fig. 5D-E).

**Fig. 6.**
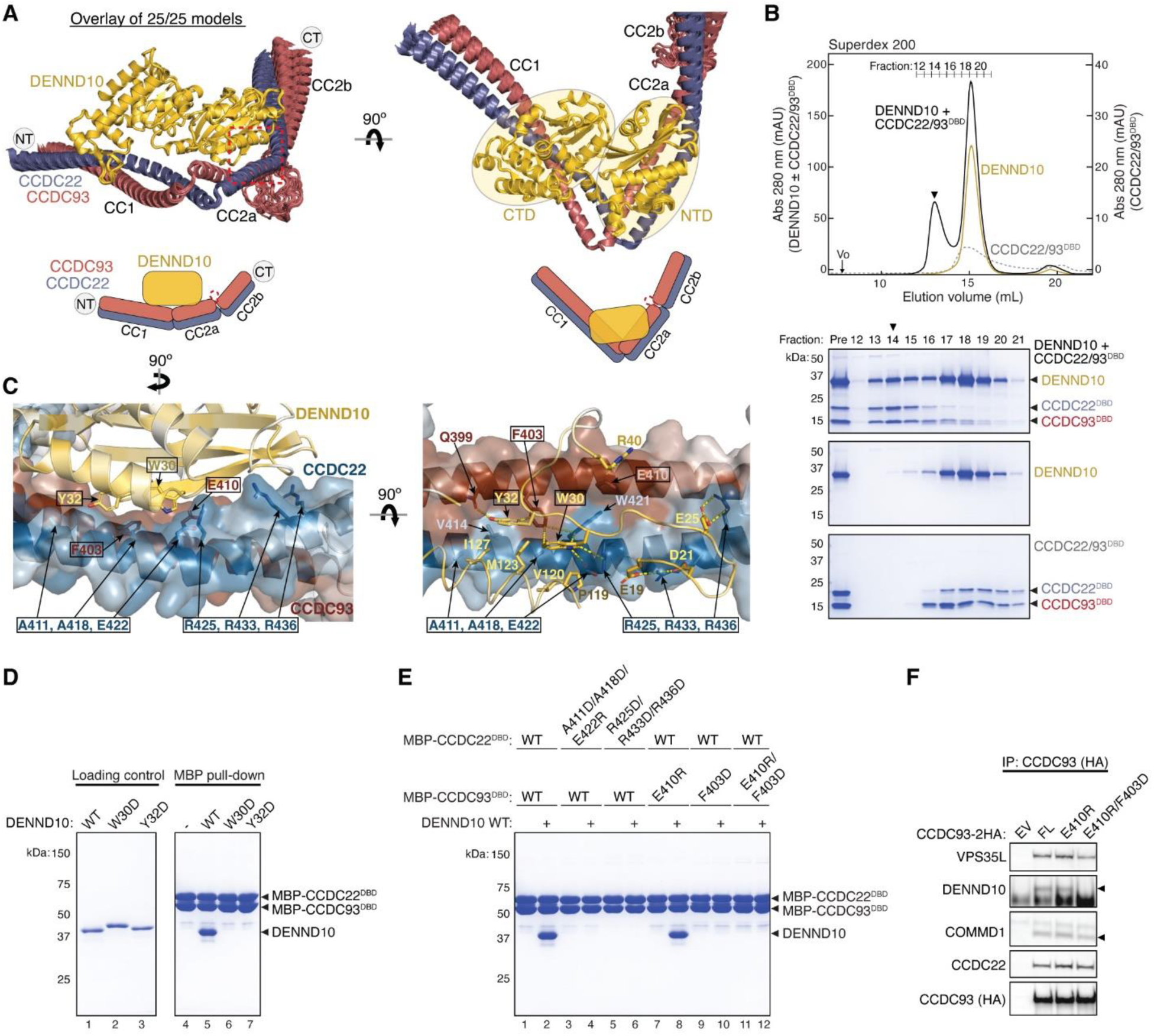
Structural model of CCDC22-CCDC93 binding to DENND10. **(A)** Overlay of all 25 AlphaFold Multimer models and schematic showing DENND10 binding to CCDC22-CCDC93. **(B)** Gel filtration of DENND10 and CCDC22-CCDC92 DBD, individually and in combination. Coomassie blue-stained SDS-PAGE gels of the indicated fractions are shown. The arrowhead indicates the peak fraction of the trimer. **(C)** Interaction surface between DENND10 and CCDC22-CCDC93 DBD colored by conservation score using the same scheme shown in Fig. 2. Key interactions are shown as sticks and polar interactions are represented with a dashed yellow line. Residues mutated in this study are outlined with a black box. **(D-E)** Coomassie blue-stained SDS PAGE gels showing MBP-tagged CCDC22-CCDC93 DBD (200 pmol) pulling down DENND10 (500 pmol). Mutations in corresponding constructs are indicated. **(F)** Immunoprecipitation of CCDC93 carrying indicated point mutants expressed in HEK293T cells and immunoblotting for the indicated proteins.

The AFM model reveals that DENND10 consists of two closely packed domains, the N-terminal domain (NTD) and the C-terminal domain (CTD), similar to the crystal structure of the DENN domain from DENND1^35^ (Fig. 6A, Extended Data Fig. 6D). DENND10 is positioned above the junction between the CC1 and CC2a coiled coils, where the coiled coils make a sharp turn to adopt a V-shaped configuration (Fig. 6A). We hereafter refer to the CC1 and CC2a coiled coils as the DENND10 binding domain (DBD). While it remains unknown whether DENND10 possesses Rab GEF activity like other DENN-domain containing proteins, the probable GTPase binding surface is partially blocked by the interaction with the CCDC22-CCDC93 DBD (Extended Data Fig. 6D).

To validate the predicted structure, we purified the DBD heterodimer and DENND10 recombinantly and used size-exclusion chromatography to test whether they can directly interact with each other. Individually, the untagged DBD dimer and DENND10 eluted at ∼15 mL, corresponding to their similar molecular weight of ∼40 kDa. When the two components were combined, a new peak appeared at ∼13 mL, indicating the formation of a complex (Fig. 6B). The peak contained all three proteins in near 1:1:1 stoichiometry, confirming the direct interaction and stable complex formation between the DBD and DENND10 (Fig. 6B).

To further validate the predicted structure, we used MBP pull-down assays and co-immunoprecipitation to test if mutations in the conserved residues predicted to be crucial for the interaction could disrupt the binding. Consistent with the AFM model, the W30D and Y32D mutations in DENND10 completely abolished its binding to CCDC22-CCDC93 DBD (Fig. 6D). Both residues are located at the center of the interaction surface between the NTD of DENND10 and the CC2a coiled coil of CCDC22-CCDC93, where they form extensive *van der Waals*, polar, and π-π interactions with residues from both CCDC22 (V414, A418, W421, E422, and R425) and CCDC93 (Q399 and F403) (Fig. 6C). Similarly, mutating surface residues in CCDC22 DBD (A411D/A418D/E422R or R425D/R433D/R436D) or CCDC93 DBD (F430D or E410R/F403D) also abolished the interaction (Fig. 6C, E). The E410R mutation in CCDC93 DBD did not cause an appreciable effect, likely due to its location at the periphery of the interaction surface (Fig. 6C, E). These results are consistent with our co-immunoprecipitation data when these mutant proteins were expressed in cells, showing that while E410R had minimal impact on the binding to DENND10, the E410R/F403D double mutation completely abolished the interaction (Fig. 6F).

### CCDC22-CCDC93 binds to COMMD oligomers

While our models had unraveled the roles of the coiled coils at the C-termini of CCDC22-CCDC93 heterodimer in the binding to Retriever and DENND10, respectively, prior work had identified the significance of the N-terminal sequences of the CCDC22-CCDC93 dimer in binding to COMMD proteins^14^. The ten members of the COMMD protein family are known to readily form heterodimers both in vitro and in cells through their conserved C-terminal COMM domains^15, 17^. However, the precise stoichiometry and assembly of COMMDs in vivo remain unknown. To address these questions, we used AFM to predict how the COMMD proteins associate with each other in different combinations and stoichiometries, in the presence or absence of different fragments of CCDC22-CCDC93.

Remarkably, we could obtain a highly convergent model when we included one copy of each of the ten COMMD proteins, with or without the CCDC22-CCDC93 heterodimer. This model is consistent with our quantitative proteomic analyses of the native CCC-Retriever complex isolated from blue native gels, where the ratio of all 10 COMMD proteins, except for COMMD7, is nearly equimolar (Extended Data Table 1). The resulting AFM model consistently suggested the formation of a ring-like structure resembling a pentagram, with a height of ∼90 Å and a diameter of ∼100 Å (Fig. 7A-C, Extended Data Fig. 7). The structure consists of five COMMD-COMMD heterodimers assembled mainly through interactions between COMM domains. The ten COMMDs are arranged in a highly organized manner, following the sequence of (1/6)-(4/8)-(2/3)-(10/5)-(7/9). Subunits within the same parentheses form a heterodimer through a face-to-face “hand shaking” interaction between their COMMD domains. These heterodimers further pack into a ring through a back-to-back stacking of β-sheets with neighboring heterodimers (Fig. 7A). The resulting ring formed by the COMMD domains provides a flat pentagram base of ∼20 Å in thickness, while the N-terminal globular domains are positioned on alternating sides of the ring (Fig. 7B). The five COMMDs facing the same clockwise direction, 1-4-2-10-7 or 6-9-5-3-8, orient their N-terminal globular domains towards the same side of the ring, approximately above the COMMD domains of the subsequent heterodimer (Fig. 7B). When viewed from the top or bottom of the COMMD ring, the globular domains project in a counterclockwise order from their corresponding COMM domain. Because COMMD6 in humans lacks a globular domain, the COMMD ring has five globular domains on one side and four on the other side.

**Fig. 7.**
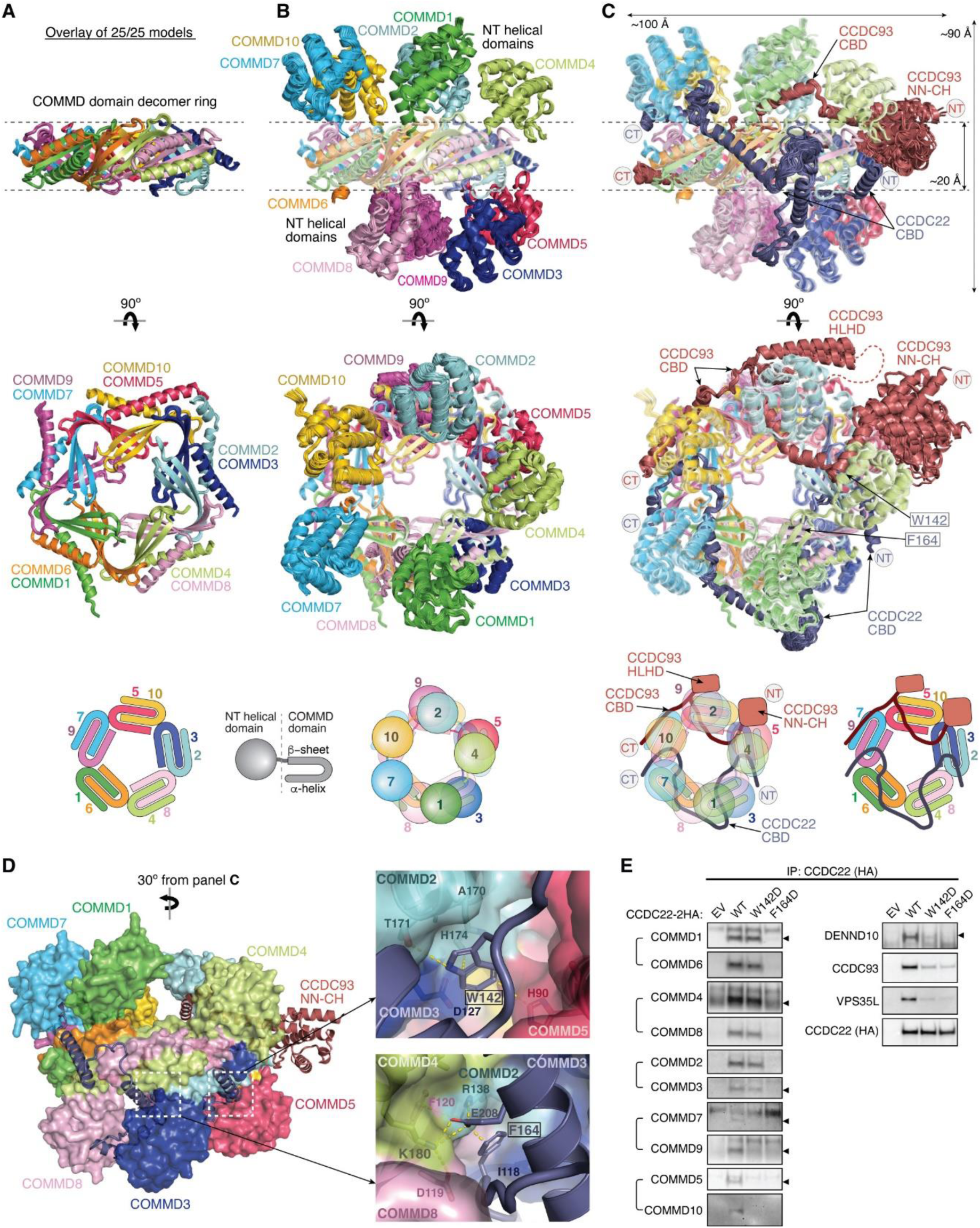
Structural model of CCDC22-CCDC93 binding to COMMD. **(A-C)** Overlay of all 25 AlphaFold Multimer models and schematic showing COMMD decamer ring binding to CCDC22-CCDC93, with (A) highlighting the central ring of the COMM domain, (B) highlighting the globular domains on the two sides of the ring, and (C) highlighting the conformation of CCDC22 and CCDC93 CBDs. **(D)** Interaction surface between the COMMD ring (surface representation) with CCDC22-CCDC93 CBDs (cartoon). Key interactions are shown as sticks and polar interactions are represented with a dashed yellow line. Residues mutated in this study are outlined with a black box. **(E)** Immunoprecipitation of CCDC22 carrying indicated point mutations expressed in HEK293T cells and immunoblotting for the indicated proteins.

The NT regions of CCDC22 (a.a. 125-261) and CCDC93 (a.a. 130-304) separately make extensive contacts with two opposite sides of the COMMD ring, resembling “tweezers” pinching into a “donut” (Fig. 7C). We hereafter refer to these regions as the COMMD binding domain (CBD) of CCDC22 and CCDC93, respectively. The CCDC93 CBD binds to one side of the ring encompassed by (2/3)-(10/5)-(7/9), while the CCDC22 CBD binds to the opposite side of the ring encompassed by (3/2)-(8/4)-(6/1)-(9/7). The N-terminal NN-CH domain of CCDC93 (a.a. 1-130) does not directly interact with the COMMD ring, for which AFM could not identify a consistent position relative to the ring and showed poor alignment relative to the rest of the structure (Fig. 7C). Immediately following the NN-CH domain of CCDC93 is the CBD, which adopts several helices that wind through the space between the globular domains and the central COMM domain base. Within the CBD is a region containing two short β strands, a short helix, and a long helix, separated by an extensive unstructured loop. We name this region the helix-loop-helix domain (HLHD) of CCDC93. The majority of the HLHD domain does not directly interact with the COMMD ring (Fig. 7C). As for CCDC22, its NN-CH domain is already involved in stabilizing the VBD domain (Fig. 5A) and is separated from the CBD by a short, flexible linker of ∼ 15 residues. The CBD of CCDC22 adopts a more extended conformation than CCDC93, winding through a larger area on other side of the COMMD ring.

To validate the AFM model, we extended our predictions of the COMMD ring in other species, including fish and amoeba, which possess all the 10 COMMD proteins, as well as CCDC22 and CCDC93. Strikingly, the positioning of COMMD orthologs within the ring follows the exact sequence predicted for human proteins (Extended Data Fig. 7). Similarly, the regions of CCDC22 and CCDC93 that interact with the ring and the points of contact on the ring itself are also highly consistent across these species (Extended Data Fig. 7). It is interesting to note that the globular domains of COMMD6 and COMMD9 show variations between species. While humans lack the globular domain of COMMD6, zebrafish possess it, but amoeba lacks the globular domains from both COMMD6 and COMMD9, reflecting the evolutionary divergence in the structure and composition of the COMMD ring across species.

Using the AFM model as a guide, we introduced specific mutations to the residues in the CBD of CCDC22 that are predicted to make critical contacts with the COMMD ring and therefore are likely crucial for CCC assembly. These mutations include W142D and F164D (Fig. 7D). W142 is located within a cavity formed between COMMD2, COMMD3, and COMMD5, making *van der Waals*, polar, and π-π interactions with A170, T171, and H174 in COMMD2 and H90 in COMMD5. F164 is sandwiched between a helix of CCDC22 itself and a composite surface formed by COMMD2/4/3/8, where it forms a cation-π interaction with R138 in COMMD2 and hydrophobic interactions with I118 in COMMD3. The conformation of R138 is further stabilized by a cation-π interaction with F120 from COMMD4 and E208 from the CBD of CCDC22. When these mutant proteins were expressed in cells, they indeed impacted the binding to COMMD proteins (Fig. 7E). CCDC22 F164D failed to bind to all tested COMMD proteins, supporting the accuracy of the AFM model. In contrast, CCDC22 W142D showed an intriguing patter of disruption: it retained normal binding to the COMMDs present on the same side of the ring as CCDC22, including COMMD dimers (1/6), (4/8), and (2/3), but exhibited poor binding to COMMDs on the opposite side of the ring contacted by CCDC93 CBD, including (9/7) and (5/10) (Fig. 7E). Notably, both mutations reduced the binding of CCDC22 to Retriever, DENND10 and CCDC93 (Fig. 7E), even though the mutated residues are not expected to directly interact with these proteins. These results suggest that the binding to the COMMD ring creates a supra-structure that may be critical to support other protein-protein interactions involved in the proper assembly and function of the Retriever-CCC complex.

### Overall model of the Retriever-CCC assembly

With the above knowledge of the individual components of the Retriever-CCC assembly, it becomes evident that the CCDC22-CCDC93 dimer serves as a scaffold to connect three separate components, Retriever, DENND10, and the COMMD ring, with each section of the CCDC22-CCDC93 dimer forming a stable subcomplex with the corresponding component (Fig. 8A). We next wondered what configuration the entire Retriever-CCC complex would adopt when fully assembled.

**Fig. 8:**
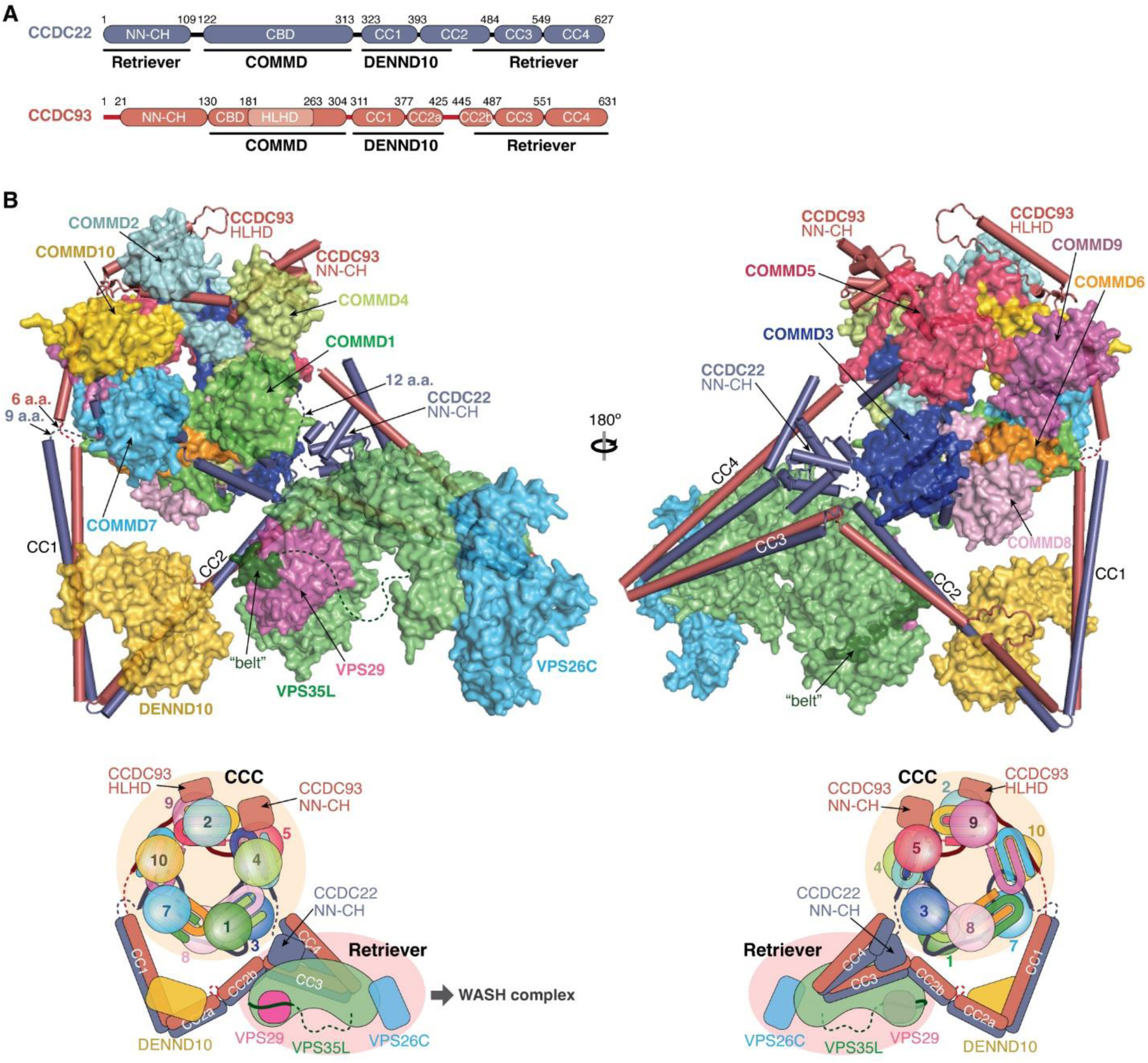
Overall model of the Retriever-CCC complex. **(A)** Schematic showing the domain organization and the corresponding interaction partners of CCDC22 and CCDC93 derived from AlphaFold Multimer prediction. **(B)** Overall structural model and schematic of the Retriever-CCC complex derived from AlphaFold Multimer prediction of individual subcomplexes. The peptide linkers in CCDC22 and CCDC93 serving as distance constraints are shown as dashed lines.

To assemble the complete Retriever-CCC complex, we “stitched” the three subcomplexes, including VBD-Retriever, DBD-DENND10, and CBD-COMMD ring, by aligning their overlapping regions. The assembly must satisfy two constraints: first, the N-terminal NN-CH domain of CCDC22 needs to interact with the C-terminal VBD, for which CCDC22 has to adopt a “looped” configuration, and second, there are short peptide linkers between the NN-CH domain of CCDC22 and its CBD (12 a.a.), between the CBD and DBD in CCDC22 (9 a.a.), and between CBD and DBD in CCDC93 (6 a.a.). These short peptide linkers limit the placement of the two opposite sides of the CBD-COMMD ring relative to VBD-Retriever and DBD-DENND10 (Fig. 8B, dashed lines). With these constraints, the final Retriever-CCC complex, which comprises 16 polypeptide chains, unambiguously adopts a compact configuration. When seen from the side, the complex resembles a “scorpion”, with Retriever forming the body, and the COMMD ring appearing like a curled tail. Within this complex, the COMMD ring is locked into a position where the globular domains of COMMD1 and COMMD3 and the heterodimeric COMM domains of COMMD8 and COMMD2 are in close proximity to the back ridge of VPS35L. As there is no evidence that the COMMD proteins can directly interact with Retriever, the location of the COMMD ring is mainly determined by the peptide linkers in CCDC22 and CCDC93, rather than through a specific interaction surface on Retriever. Therefore, while the relative position and orientation of the COMMD ring with respect to the rest of the complex are expected to remain stable, the internal components of the Retriever-CCC complex are likely to exhibit a certain degree of flexibility. This flexibility may play a role in the binding to regulatory molecules or cargo proteins.

## Discussion

Retriever and CCC play crucial roles in cellular and organismal physiology, as evident by myriad phenotypes identified thus far when these systems are disturbed, ranging from developmental alterations in humans and mouse models^36–41^, to changes in copper^25, 42–44^ and lipid metabolism^24, 45, 46^, and alterations in immune signaling^47–49^ and function^50–53^. However, despite two decades of work in this area, the structural mechanism underlying the assembly of these complexes remained elusive. Here, our study fills this knowledge gap by presenting the first high-resolution structure of Retriever and an experimentally validated model of Retriever-CCC assembly. Our data provide important structural and molecular insights into their implications in cell biology and human disease.

One key observation reported here is that cancer-associated missense mutations in VPS35L can dramatically affect Retriever assembly. The precise mechanism by which the loss of Retriever assembly may affect the oncogenic process remains to be determined. Our findings open the door to further investigate the potential client proteins whose disrupted recycling and cellular localization may be advantageous during cancer development and progression. More broadly, the structural model provides a comprehensive roadmap to understand how other complexes and cofactors may interact with this assembly and how inherited mutations in CCC and Retriever subunits, which result in Ritscher-Schinzel/3C syndrome^37, 38^, may disrupt protein function.

The model of Retriever highlights key distinctions from Retromer, in both structure and regulation. Retriever has a more compact structure and a unique mechanism of intramolecular interaction between the NT and CT portions of VPS35L. In particular, the “belt” sequence at the NT of VPS35L provides a direct binding interface for VPS29, a feature completely absent in Retromer. Another feature unique to Retriever is the long unstructured peptide linker in VPS35L that follows the “belt” sequence. The primary sequence of this serine-rich unstructured linker is highly conserved in vertebrates, suggesting that it may be a site for regulatory interactions or post-translational modifications, which remain to be elucidated.

Our study also uncovered how Retriever encounters CCC to assemble a larger complex. We find that the CCDC22-CCDC93 heterodimer is the essential scaffold around which all the components, including DENND10, are assembled. Rather than being “beads on a string”, the structure of Retriever-CCC is highly compact and selectively oriented, with only limited internal flexibility. If we extrapolate a potential orientation for the Retriever-CCC complex guided by the orientation of Retromer on endosomal membranes^54^, in which the concave aspect of VPS35 faces the membrane, the prediction would orient the COMMD ring toward the cytosolic environment where it can potentially interact with regulatory molecules or phospholipid-containing vesicles. Within this assembly, the contributions of its individual components remain to be fully understood. For example, the role of DENND10 and its putative GEF activity in this complex is unclear. While Retriever-CCC is localized to WASH-positive early endosomes, DENN10 has been reported to localize to late endosomes and multivesicular bodies, and to act on Rab27a/b^26^, which have not been implicated as cellular or molecular targets of Retriever-CCC. Therefore, these observations remain to be reconciled and expanded.

One crucial question to be addressed is the functional importance of the COMMD ring and the exquisite conservation of its assembly through evolution. Our studies show that mutations that prevent binding of CCDC22 to the ring also impaired its ability to dimerize with CCDC93 or bind to DENND10 and VPS35L. Based on these observations, we propose that the COMMD ring helps assemble or stabilize the CCDC22-CCDC93 heterodimer and is therefore essential to the assembly of the entire complex. This could explain why the CCDC22-CCDC93 heterodimer is destabilized whenever any given COMMD protein is knocked out in cells or in tissues^14, 25, 46^.

The COMMD ring assembly may have intermediate states. We previously reported^41^ that when COMMD9 is knocked out, CCDC22 cannot bind COMMD5 or COMMD10 but retains its ability to bind to COMMD6, COMMD4, COMMD8. Our structural model of the CCC complex provides a plausible explanation for this selective binding, when taking into account the points of contact made by CCDC22 and CCDC93 with the ring. According to the model, CCDC22 contacts specific COMMD dimers, (1/6), (4/8), and (2/3), while CCDC93 CBD mainly contacts (9/7) and (5/10). Therefore, deleting COMMD9 would disrupt the CCDC93 part of the ring, including COMMD5 and COMMD10, without affecting the other half of the ring stabilized by CCDC22, including COMMD6, COMMD4 and COMMD8 (Fig. 7E). In this study, we observed similar phenomena using point mutations in the CBD of CCDC22. CCDC22 F164D destabilized the binding to all COMMD proteins, likely because this residue is in the middle of the CBD contacting a combined surface created by COMMD2/4/3/8. In contrast, the disruption caused by CCDC22 W142D is highly selective. The W142D mutant binds to all the COMMD proteins mainly contacted by CCDC22 CBD, including (1/6), (4/8), and (2/3), but not to any of the COMMD proteins mainly contacted by CCDC93 CBD, including (9/7) and (5/10), analogous to the effect of COMMD9 deficiency. This is likely because W142D is located at the NT of the CBD, contacting an interface composed by COMMD (2/3)-5.

Based on these results, we propose that the ring may be formed by two half intermediate precursors, with each half stabilized independently by CCDC22 or CCDC93. The formation of the ring promotes dimerization and stabilization of CCDC22-CCDC93, providing a platform for VPS35L and DENND10 binding. Evidence of these precursor complexes is present in our proteomic data, where CCDC93 is detected at about 240 kDa Mw in blue native gels in ∼1:1 equimolar ratio with its associated COMMD proteins (Extended data Table 1). Altogether, these observations suggest that the complex may be dynamically regulated, rather than being a static entity. The mechanism by which the ring is fully assembled from precursor complexes will likely play a key role in the function of the CCC complex.

Another intriguing feature of the COMMD ring is the highly conserved order of assembly of its ten COMMD proteins. Presumably, the high conservation of each of the COMMD family members is essential to yield this arrangement. In the structure, the central part of the ring is created by COMM domain-mediated heterodimers, which are then the building blocks to assemble the ring. Therefore, we postulate that unique sequence variability in the COMM domains favors specific heterodimers over others. The model also shows that these heterodimers are further “glued” together by the CBDs of CCDC22 and CCDC93, as well as the N-terminal globular domains of each COMMD through conserved interfaces. Since the globular domains of COMMD proteins provide most of the exposed surface of the ring, we speculate that they likely provide key interfaces for regulatory interactions between the ring and other proteins.

Finally, concurrent with our effort, two other groups independently provided other structural aspects of this assembly^55, 56^. These studies are complementary to ours, as they did not resolve the experimental structure of Retriever but determined the cryo-EM structure of the CCC ring, which turned out to be highly similar to our predicted and tested models. Therefore, our study is unique in providing the high-resolution cryo-EM structure of Retriever and uncovering the presence of cancer-associated mutations in VPS35L that impair Retriever assembly. Together, our work opens the door to answer many questions related to Retriever-CCC in biology and disease, including endosomal membrane recruitment, client protein recognition, interaction with various regulators, dynamic assembly of the holo-complex, and function of distinct parts of these complexes.

## Methods

### Plasmids

All constructs used here were created by standard molecular biology procedures, verified by Sanger sequencing. Constructs used for recombinant protein production are described in detail in Supplementary Table 1, along with the resulting recombinant proteins, (Supplementary Table 2), and the DNA oligonucleotides used for construct generation (Supplementary Table 3). The ORF for VPS35L that was PCR amplified using the IMAGE clone 6452778 as a template, and codes for a 963 amino acid protein listed as isoform 1 in NCBI Gene (Gene ID: 57020). The ORF for CCDC22 and CCDC93 have been previously described^14, 49^. VPS29 and DENND10 ORF’s were PCR amplified using the IMAGE clones 3461977 and 4688412, respectively.

Site-Directed Mutagenesis was performed using Platinum SuperFi II DNA polymerase (Thermo Fisher Scientific, Waltham, MA) to generate different mutant constructs. Human full-length VPS35L (untagged), full-length VPS26C (untagged), and full-length VPS29 (isoform 2) containing a C-terminal (GGS)_2_-His_6_ tag were cloned in a modified pFastBac^TM^ vector for insect cell expression (PMID: 19363480). Fragments of human CCDC22 and CCDC93 and full-length human DENND10 were ordered as GeneStrings (ThermoFisher) with the gene codon optimized for *E. coli* expression. The Head-VPS35L-binding domain (VBD) of CCDC22 and the VBD of CCDC93 contains MBP-CCDC22 (1-118)-(GGSK)_6_-(436-727) and MBP-CCDC93 (442-631), respectively. The DENND10-binding domain (DBD) of CCDC22 and CCDC93 contains MBP-CCDC22 (280-446) and MBP-CCDC93 (305-433), respectively.

### E. coli strains for protein expression

Standard, commercial *E. coli* strains used in this study include Mach1^T1R^ (Thermo Fisher), BL21 (DE3)^T1R^ (Sigma), and ArcticExpress^TM^ (DE3)RIL cells (Stratagene), and are grown in Luria-Bertani or Terrific Broth medium using standard molecular biology conditions.

### Insect cell lines for protein expression

Standard, commercial insect cell lines used this study include *Sf9* cells (Invitrogen) and *Tni* (High-Five) cells (Expression System). *Sf9* cells were used for baculovirus preparation and maintained in Sf900 II serum free medium (Thermo Fisher). *Tni* cells were used for large-scale protein expression and are maintained in ESF 921 serum free medium (Expression Systems).

### Cell culture

HEK293T and HeLa cell lines were obtained from the American Type Culture Collection (Manassas, VA). Huh-7 cell lines were a gift from Dr. Jay Horton (University of Texas Southwestern Medical Center, Dallas, TX, USA). All cell lines were cultured in high-glucose Dulbecco’s modified Eagle’s medium (DMEM) containing 10% fetal bovine serum (FBS) and 1% penicillin/streptomycin, in incubators at 37⁰C, 5% CO2. Periodic testing for Mycoplasma spp. using PCR based detection was performed to exclude infection of the cultures. HeLa cells with VPS26C deficiency (generated using CRISPR/Cas9) and complemented with HA-tagged VPS26C have been previously reported^23^. A HeLa line with COMMD1 deficiency was previously reported^57^; these cells were complemented using a lentiviral vector with HA-tagged COMMD1.

### CRISPR/Cas9-mediated gene deletion

VPS35L and VPS29 knockout cell lines were generated using CRISPR/Cas9, as previously described. Briefly, in vitro assembled Cas9-ribonucleotide complexes were transfected onto Huh7 cells for VPS35L and onto HeLa cells for VPS29. The degree of VPS35L or VPS29 protein expression was examined in this polyclonal population pool through immunoblotting, and if deemed greater than 50% decreased from the parental cells, the transfected pool was subjected to clonal isolation. Clones were isolated through limiting dilution and screened by immunoblot for protein expression. CRIPSR guide RNA sequences used in this study are listed in Supplementary Table 4.

### Transfection and lentiviral methods

Lipofectamine 2000 (Life Technologies) was used to transfect plasmids in HEK 293T cells and cultured for either 24 or 48 hours before analysis. VPS35L Huh-7 knockout cells generated by CRISPR/Cas9 as detailed above were reconstituted with either HA empty vector or HA-tagged VPS35L wild type or mutants, using a lentivirus system. All lentivirus experiments were performed with a standard viral vector production and selection protocol as previously described.

### Immunofluorescence staining

We followed the protocols that we have previously reported^14, 22^. In brief, cells were fixed in cold fixative (4% paraformaldehyde in phosphate-buffered saline; PBS) and incubated for 18 min at room temperature in the dark, followed by permeabilization for 3 min with 0.15% Surfact-Amps X-100 (28314, Thermofisher Scientific, Rockford, IL) in PBS. Samples were then incubated overnight at 4 °C in a humidified chamber with primary antibodies in immunofluorescence (IF) buffer (Tris-buffered saline plus human serum cocktail). After three washes in PBS, the samples were incubated with secondary antibodies (1:500 dilution in IF buffer) for 1 h at room temperature or overnight at 4 °C in a humidified chamber. After four washes in PBS, coverslips were rinsed in water and affixed to slides with SlowFade Anti-fade reagent (Life Technologies, Grand Island, NY). The primary and secondary antibodies used for staining are detailed in Supplementary Table 5. Alexa Fluor 488– phalloidin (A12379, Life Technologies, Grand Island, NY) was used to visualize F-actin. Images were obtained using an A1R confocal microscope (Nikon, ×60 /1.4 oil immersion objective). Fluorescence signal values were quantified using Fiji (ImageJ, NIH). Data were processed with Excel (Microsoft) and plotted with Prism6 (GraphPad). Each dot represents the value from a single cell; the horizontal bar in these graphs represents the mean and the error bars correspond to the standard deviation (SD). Pearson’s correlation coefficient was measured using Colocalization Threshold Fiji Plugin within manually outlined regions of interest (ROIs).

### Flow cytometry

Cells were detached from plates using a cell scraper in 1 x PBS and spun at 3,000 RPM for 5 min. The cells were resuspended in fresh PBS and rinsed one time with a repeat spinning step. For CD14 staining cells were immediately processed and resuspended in FACS buffer (PBS, 1% BSA) containing CD14 antibody for 30 min on ice protected from light. After this, cells were rinsed by spinning and resuspension in FACS buffer, 3 times. For Villin staining, cells were fixed and permeabilized using BD Cytofix/Cytoperm solution kit according to the manufacturer’s instructions (BD Biosciences). Thereafter, they were incubated with Villin antibody overnight at 4⁰C in BD Perm/wash buffer, followed by 3 washes using the same buffer.

Thereafter, secondary antibody incubation was performed in BD Perm/wash buffer, followed by 3 washes using the same buffer. The primary and secondary antibodies used for staining are detailed in Supplementary Table 5. Samples were processed by the Flow Cytometry core at UT Southwestern using a Cytek Aurora instrument. Data analysis was performed using FlowJo software.

### Mammalian protein extraction, immunoblotting and immunoprecipitation

For most experiments, whole cell lysates were prepared by adding Triton X-100 lysis buffer (25 mM HEPES, 100 mM NaCl, 10 mM DTT, 1 mM EDTA, 10% Glycerol, 1% Triton X-100) supplemented with protease inhibitors (Roche). Immunoprecipitation, SDS-PAGE, and immunoblotting experiments were performed as previously described. The antibodies used for immunoprecipitation and immunoblotting are detailed in Supplementary Table 5.

### Blue native electrophoresis and immunoblotting

Cell lysate preparation was performed using MRB buffer as lysis buffer (20 mM HEPES pH 7.2, 50 mM potassium acetate, 1 mM EDTA, 200 mM D-Sorbitol, 0.1% Triton X-100). After immunoprecipitation using HA affinity beads (Roche), associated complexes bound to 2xHA-VPS35L, HA-COMMD1 or HA-VPS26C were eluted using lysis buffer containing HA peptide (1 mg/mL). The eluted complexes were then loaded to NativePAGE^TM^ 3-12% Bis-Tris protein gels, with one lane containing NativeMark^TM^ Unstained protein standard. For immunoblotting, the proteins in the gel were transblotted to PVDF membranes. After transfer, the proteins were fixed by incubating the membrane in 8% acetic acid for 15 minutes, followed by immunoblotting as described above. For proteomic experiments, gels were stained with Coomassie blue and gel slices of specific apparent mass were cut and submitted for analysis.

### Cell surface biotinylation

Cell surface biotinylation was performed as previously reported^14^. Briefly, cells were incubated at 4⁰C with Sulfo-NHS-SS-biotin (Pierce, Rockford, IL) in biotinylation buffer (10 mM triethanolamine, 150 mM NaCl, 2 mM CaCl2, pH 8.0). After 30 min of labeling, cells were lysed in Tris-lysis buffer (50 mM Tris-HCl, pH 7.4, 150 mM NaCl, 1% NP-40, 0.5% Na deoxycholate, 5 mM EDTA, 5 mM EGTA) supplemented with protease inhibitors (Halt Protease/Phosphatase inhibitor, Thermofisher). Biotinylated proteins were precipitated using nanolink Streptavidin magnetic beads (Solulink) and washed 3 times with the same lysis buffer, 1 time in high salt buffer (50 mM Tris-HCl, pH 7.4, 500 mM NaCl), and 1 time in low salt buffer (10 mM Tris-HCl, pH 7.4, 5 μM Biotin). Precipitated protein-containing beads were eluted using 3 x LDS/DTT gel loading buffer at 95⁰C. The samples were loaded on an SDS/PAGE gel and the bands cut from the stacker portion of the gel to be submitted to the Proteomics core at UT Southwestern for further processing. For TMT proteomics, the eluted proteins in solution were directly submitted to the core.

### Protein affinity purification

HA-, HA-VPS35L wild-type or mutants transduced knockout cells were grown on cell culture dishes and lysed in Triton-X lysis buffer. Cell lysates were cleared, and equal amounts of protein were then added to HA-resin to capture HA-tagged proteins. HA beads were washed using lysis buffer and proteins were eluted using HA peptide (1 mg/mL). Eluted material was resuspended in 3x LDS sample buffer with DTT. Proteins were then subjected to SDS-PAGE and analyzed by LC-MS/MS mass spectrometry at the UT Southwestern Proteomics core.

### Proteomic interactome and cell surface analysis

Protein identification, abundance (based on spectral index) and enrichment ratios (compared to empty vector) were utilized to identify putative interacting proteins. This process consisted first of overnight sample digestion with trypsin (Pierce) following reduction and alkylation with DTT and iodoacetamide (Sigma–Aldrich). The samples then underwent solid-phase extraction cleanup with an Oasis HLB plate (Waters) and the resulting samples were injected onto an Orbitrap Fusion Lumos mass spectrometer coupled to an Ultimate 3000 RSLC-Nano liquid chromatography system. Samples were injected onto a 75 μm i.d., 75-cm long EasySpray column (Thermo) and eluted with a gradient from 1-28% buffer B over 90 min. Buffer A contained 2% (v/v) ACN and 0.1% formic acid in water, and buffer B contained 80% (v/v) ACN, 10% (v/v) trifluoroethanol, and 0.1% formic acid in water.

The mass spectrometer operated in positive ion mode with a source voltage of 1.8-2.4 kV and an ion transfer tube temperature of 275 °C. MS scans were acquired at 120,000 resolution in the Orbitrap and up to 10 MS/MS spectra were obtained in the ion trap for each full spectrum acquired using higher-energy collisional dissociation (HCD) for ions with charges 2-7. Dynamic exclusion was set for 25 s after an ion was selected for fragmentation.

For the plasma membrane and interaction proteomics samples, raw MS data files were analyzed using Proteome Discoverer v3.0 (Thermo), with peptide identification performed using Sequest HT searching against the human protein database from UniProt. Fragment and precursor tolerances of 10 ppm and 0.6 Da were specified, and three missed cleavages were allowed. Carbamidomethylation of Cys was set as a fixed peptide modification, with oxidation of Met set as a peptide variable modification. The false-discovery rate (FDR) cutoff was 1% for all peptides.

For protein complex composition in native gel samples, raw MS data files were analyzed using MaxQuant v.2.0.3.0, with peptide identification performed against the human protein database from UniProt. Fragment and precursor tolerances of 20 ppm and 0.5 Da were specified, and three missed cleavages were allowed. Carbamidomethylation of Cys was set as a fixed peptide modification, oxidation of Met was set as a peptide variable modification, and N-terminal acetylation was set as a protein variable modification. iBAQ quantitation was used for performing protein quantitation within each sample.

### TMT proteomics

Samples used for TMT-based proteomic quantification were processed as follows. 25 μL of 10% SDS, 100 mM triethylammonium bicarbonate (TEAB) was added to each sample and vortexed to mix. 2 μL of 0.5M tris(2-carboxyethyl)phosphine (TCEP) was added to reduce the disulfide bonds and the samples were incubated for 30 min at 56⁰C. The free cysteines were alkylated by adding 2 μL of 500 mM iodoacetamide (IAA) to a final IAA concentration of 20 mM, and the samples were incubated in the dark at room temperature for 30 min. 5.4 μL of 12% phosphoric acid was added to each sample along with 300 μL of S-Trap (Protifi) binding buffer, and the samples were loaded on to an S-Trap column. 1 μg of trypsin was added and the sample was digested overnight at 37⁰C. Following digestion, the peptide eluate was dried and reconstituted in 21 μL of 50 mM TEAB buffer. Equal amounts of the samples based on NanoDrop A(205) reading were labelled with TMT 6plex reagent (Thermo), quenched with 5% hydroxylamine, and combined. The peptide mixture was dried in a SpeedVac, desalted using an Oasis HLB microelution plate (Waters), and dried again in a SpeedVac. 50 μL of 2% acetonitrile, 0.1% TFA was added to the sample, and this was injected onto an Orbitrap Eclipse mass spectrometer coupled to an Ultimate 3000 RSLC-Nano liquid chromatography system. Samples were injected onto a 75 μm i.d., 75-cm long EasySpray column (Thermo) and eluted with a gradient from 1-28% buffer B over 180 min, followed by 28-45% buffer B over 25 minutes. Buffer A contained 2% (v/v) ACN and 0.1% formic acid in water, and buffer B contained 80% (v/v) ACN, 10% (v/v) trifluoroethanol, and 0.1% formic acid in water. The mass spectrometer operated in positive ion mode with a source voltage of 2.0 kV and an ion transfer tube temperature of 300°C. MS scans were acquired at 120,000 resolution in the Orbitrap over a mass range of m/z = 400-1600, and top speed mode was used for SPS-MS3 analysis with a cycle time of 2.5 s. MS2 was performed using collisionally-induced dissociation (CID) with a collision energy of 35% for ions with charges 2-6. Dynamic exclusion was set for 25 s after an ion was selected for fragmentation. Real-time search was performed using the reviewed human protein database from UniProt, with carbamidomethylation of Cys and TMT 6plex modification of Lys and peptide N-termini set as static modifications and oxidation of Met set as a variable modification. Two missed cleavages and up to 3 modifications per peptide were allowed. The top 10 fragments for MS/MS spectra corresponding to peptides identified by real-time search were selected for MS3 fragmentation using high-energy collisional dissociation (HCD), with a collision energy of 65%. Raw MS data files were analyzed using Proteome Discoverer v3.0 (Thermo), with peptide identification performed using both Sequest HT and Comet searching against the human protein database from UniProt. Fragment and precursor tolerances of 10 ppm and 0.6 Da were specified, and two missed cleavages were allowed. The same modifications were used in the search as for the real-time search. The false-discovery rate (FDR) cutoff was 1% for all peptides.

### Recombinant protein purification

Retriever was expressed by co-infecting Sf9 cells at 2 M/mL cell density (Expression Systems) with individual baculoviruses prepared by the Bac-to-Bac system used in other studies^58, 59^ (Invitrogen). Retriever was purified by Ni-NTA agarose beads (Qiagen), followed by cation exchange chromatography using a 2-mL Source 15S column, anion exchange chromatography using a 1-mL Capto^TM^ HiRes Q 5/50, and size exclusion chromatography using a 24-mL Superdex Increase 200 column. CCDC22 and CCDC93 fragments and DENND10 were expressed in BL21 (DE3)^T1R^ cells (Sigma) at 18 °C overnight after induction with 1mM IPTG. MBP-tagged CCDC22 and CCDC93 proteins were purified using Amylose beads (New England Biolabs), mixed in approximately 1:1 stoichiometry, incubated overnight at 4°C to promote dimer formation, and were further purified by anion exchange chromatography using a Source 15Q column and size exclusion chromatography using a Superdex 200 column. His_6_-Tev-DENND10 was purified using Ni-NTA agarose resin (Qiagen), followed by anion exchange chromatography using a Source 15Q column, and size exclusion chromatography using a 24-mL Superdex Increase 200 column. All ion exchange and gel filtration chromatography steps were performed using columns purchased from Cytiva on an ÄKTA^TM^ Pure protein purification system.

### Size exclusion chromatography analysis

MBP-CCDC22 DBD and MBP-CCDC93 DBD proteins were assembled into a dimer after amylose purification and further purified using a Source 15Q column, followed by TEV cleavage overnight to remove the MBP tags. His_6_-DENND10 was purified using a Source 15Q column and treated with TEV protease overnight to remove the His_6_ tag. The cleaved CCDC22-CCDC93 DBD dimer and DENND10 were further purified using a Source 15Q column. The concentration was determined using the absorbance at 280 nm and the extinction coefficient generated by Expasy ProtParam. The DBD dimer and DENND10 proteins were mixed and purified over a Superdex 200 column equilibrated in 100 mM NaCl, 10 mM HEPES pH 7.0, 5% (w/v) glycerol, and 1 mM DTT. The individual dimer and DENND10 proteins were diluted to the same concentration as they were in the trimer complex with buffer and the same amount of protein was purified over the same Superdex 200 column, in the same buffer as the trimer.

### In vitro pull-down assays

MBP pull-down experiments were performed as previously described^60^. Briefly, bait (100-200 pmol of MBP-tagged proteins) and prey (60 pmol for Retriever or 500 pmol for DENND10) were mixed with 20 µL of Amylose beads (New England Biolabs) in 1 mL of binding buffer [10 mM HEPES pH 7, 150 mM NaCl, 5% (w/v) glycerol, 0.05% (w/v) Triton-X100, and 5 mM β-mercaptoethanol] at 4 °C for 30 min. The beads were washed three times with 1 mL of the binding buffer each time. Bound proteins were eluted with binding buffer supplemented with 2% (w/v) maltose and were examined by SDS-PAGE and Coomassie blue staining.

### Sample preparation for electron microscopy

The Retriever complex (3 µL at 0.25 mg/ml) in a buffer containing 10 mM HEPES (pH 7.0), 150 mM NaCl, 2 mM MgCl_2_, 2 mM DTT, and 5% (w/v) glycerol was applied to a Quantifoil 300-mesh R1.2/1.3 grid (Quantifoil, Micro Tools GmbH, Germany), which had been pre-treated with glow discharge using Pelco EasiGlow immediately before use. Following a 30-second preincubation, the grid was blotted for 4 sec under 100% humidity at 4°C and then plunged into liquid ethane using Vitrobot Mark IV (Thermo Fisher Scientific).

### Electron microscopy data acquisition

A pilot dataset was collected using a 200 kV Glacios microscope (Thermo Fisher Scientific) equipped with a K3 camera (Gatan). For large data set collection, sample grids were screened on a 200 kV Talos Artica microscope (Thermo Fisher Scientific). The final cryo-EM data were acquired on a Titan Krios microscope (Thermo Fisher Scientific) operated at 300 kV and equipped with a post-column energy filter (Gatan) and a K3 direct detection camera (Gatan) in CDS mode using SerialEM^61^. Movies were acquired at a pixel size of 0.415 Å in super-resolution counting mode, with an accumulated total dose of 60 e-/Å2 over 60 frames. The defocus range of the images was set between -1.2 to -2.4 μm. In total, 3,594 movies were collected and used for data processing.

### Electron Microscopy data processing

The cryo-EM data were processed using cryoSPARC^62^ v4.2.1. To correct for beam induced motion and compensate for radiation damage over spatial frequencies, the patch motion correction algorithm was employed using a binning factor of 2, resulting in a pixel size of 0.83 Å/pixel for the micrographs. Contrast Transfer Function (CTF) parameters were estimated using patch CTF estimation. After manual curation, a total of 2,892 micrographs were selected for further processing from the initial 3,594 micrographs. For particle picking, a 4.3 Å map of the Retriever complex obtained from the pilot dataset was used as a template, resulting in the identification of 1,221,095 particles. After 2D classification, 1,105,321 particles were selected and subjected to *ab initio* 3D reconstruction, followed by heterogeneous refinement (Extended Data Fig. 1). The best resolved 3D class, containing 426,624 particles, was selected for the final non-uniform refinement followed by the CTF refinement, producing a full map with an overall resolution of 2.94 Å with a binned pixel size of 1.0624 Å/pixel. DeepEMhancer^63^ was then used with the two unfiltered half maps to generate a locally sharpened map (EMD-40885/PDB-8SYM). To better resolve the interaction between VPS29 and VPS35L, a mask was applied around VPS29 and the adjacent C-terminal region of VPS35L and signals outside the mask were subtracted (Extended Data Fig. 1G). Next, 3D classification without alignment was applied to the subtracted particle stack, resulting in a class containing 83,654 particles with better resolved density of the “belt” sequence. Local refinement of this class resulted in a map with an overall resolution of 3.18 Å, which was further sharpened by DeepEMhancer. This map was then aligned with the full map and combined using the “*vop maximum*” function in UCSF Chimera based on the maximum value at each voxel^64^. This composite map (EMD-40886/PDB-8SYO) is used to show the overall features of the Retriever complex in Fig. 1A. All reported resolutions followed the gold-standard Fourier shell correlation (FSC) using the 0.143 criterion^65^.

### Atomic model building

A Retriever model predicted by AlphaFold Multimer 2.2.3 was used as the initial model^34^ for model building using COOT^33, 66^ with the DeepEMhancer sharpened maps. The model was built through iterations of real-space refinement in Phenix^67^ with secondary structure restraints. Model geometries were assessed using the MolProbity module in Phenix, the Molprobity server^68^ (http://molprobity.biochem.duke.edu/), and the PDB Validation server^69^ (www.wwpdb.org). Figures were rendered using PyMOL (The PyMOL Molecular Graphics System, Schrödinger) or ChimeraX^70^. Interface area calculation was done using the PISA server (https://www.ebi.ac.uk/pdbe/pisa/).

### AlphaFold prediction and analysis

Versions 2.1.1, 2.1.2, 2.2.3, 2.2.4, 2.3.0, and 2.3.1 of AlphaFold (https://github.com/deepmind/alphafold) were installed on local NVidia A100 80GB GPU computers hosted by Iowa State University ResearchIT or High-Performance Computing for performing AlphaFold Multimer prediction. The standard AlphaFold procedures were followed^33, 34^, with the following specifics. The full genetic database was used for multiple sequence alignment. For each multimeric complex, five models were generated, and five predictions were performed for each model, resulting in a total of 25 unrelaxed models. These unrelaxed structures were relaxed using Amber energy minimization and ranked based on the predicted template modeling (pTM) scores. The typical memory allocations are 128-256 GB for CPU and 80 GB for GPU. The command option “maximum template release date” was set to be 2021-11-01, as we used multiple versions of Alphafold (2.1.1, 2.1.2, 2.2.3, 2.2.4, 2.3.0, and 2.3.1) to predict structures for the same sequence since 2021, and the three databases used in AlphaFold were older than 2021-11-01 and could be obtained from the following sources: SOURCE_URL=http://wwwuser.gwdg.de/∼compbiol/data/hhsuite/databases/hhsuite_dbs/old-releases/pdb70_from_mmcif_200401.tar.gz, and SOURCE_URL=https://storage.googleapis.com/alphafold-databases/v2.3/UniRef30_2021_03.tar.gz The reliability of the predicted models was assessed by three different means, including the predicted local difference distance test (pLDDT) scores, the PAE scores, and the manual examination of the consistency of 25 solutions when they were aligned in Pymol.

### Reproducibility and statistical analysis

To assess statistical significance, one-way ANOVA with Dunnett’s post-hoc test was applied to compare multiple groups with one control group. Statistical analyses were performed using Prism 9.5.1. An error probability below 5% (p < 0.05; * in Fig. panels) was considered to imply statistical significance. All imaging, FACS, and co-precipitation experiments were performed at least in two to four independent iterations. Large scale proteomics were performed once, and key results confirmed with other methods.

## Data availability

Cryo-EM maps have been deposited in the EMDB and PDB (accessions noted in Table 1). Mass spectrometry data have been deposited at the MassIVE repository (accession numbers MSV000092100, MSV000092102, MSV000092103, MSV000092104). Source data are available for all uncropped western blots and all image and FACS quantification used in these studies. Further information and requests for resources and reagents including DNA constructs, cell lines, and structural models should be directed to and will be fulfilled by Ezra Burstein and/or Baoyu Chen. Any additional information required to reanalyze the data reported here will be shared upon reasonable request. This paper does not report original code.

## Figures and Legends

## Supporting information

Tables with reagents and materials used for the study

## Acknowledgements

We thank the ResearchIT at Iowa State University for installing AlphaFold Multimer. We also want to thank Andrew Lemoff and the Proteomics core as well as Angela Mobley and the Flow Cytometry core at UT Southwestern. Electron Microscopy data were collected in collaboration with the Structural Biology Laboratory with the help from Yang Li and the Cryo Electron Microscopy Facility at UT Southwestern Medical Center which are partially supported by grant RP220582 from the Cancer Prevention & Research Institute of Texas (CPRIT) for cryo-EM studies, and at the Iowa State University Cryo-EM Facility (supported by the Roy J. Carver Structural Initiative). Research was supported by funding from the National Institutes of Health (R35 GM128786), the National Science Foundation CAREER award (CDF 2047640), and Roy J. Carver Charitable Trust seed funds to B.C., the National Institutes of Health (R01 DK107733) to E.B. and D.D.B.

## Author Contributions

E.B., B.C., and D.D.B. conceived the project. E.B. oversaw cell biological and proteomic experiments performed by A.S. with the help from Q.L., K.S and X.L. B.C. oversaw protein purification, biochemical experiments, and AlphaFold predictions performed by D.J.B. with the help from D.A.K. and X.Z. Z.C. and Y.H. oversaw cryo-EM grid preparation, data collection, single particle reconstruction and atomic-model building. P.J. supervised initial cryo-EM grid preparation and data collection performed by D.J.B. at Iowa State. M.J.M and D.D.B. helped with cellular experiments and data interpretation. B.C., Z.C. D.J.B., and Y.H. analyzed structures. E.B., B.C., and Z.C. drafted the manuscript and prepared the Figs with assistance from all other authors.

## Ethics Declarations

The authors declare no competing interests.

## Additional Information

Correspondence and request for materials should be addressed to.

## Extended Data Figures

**Extended Data Fig. 1.**
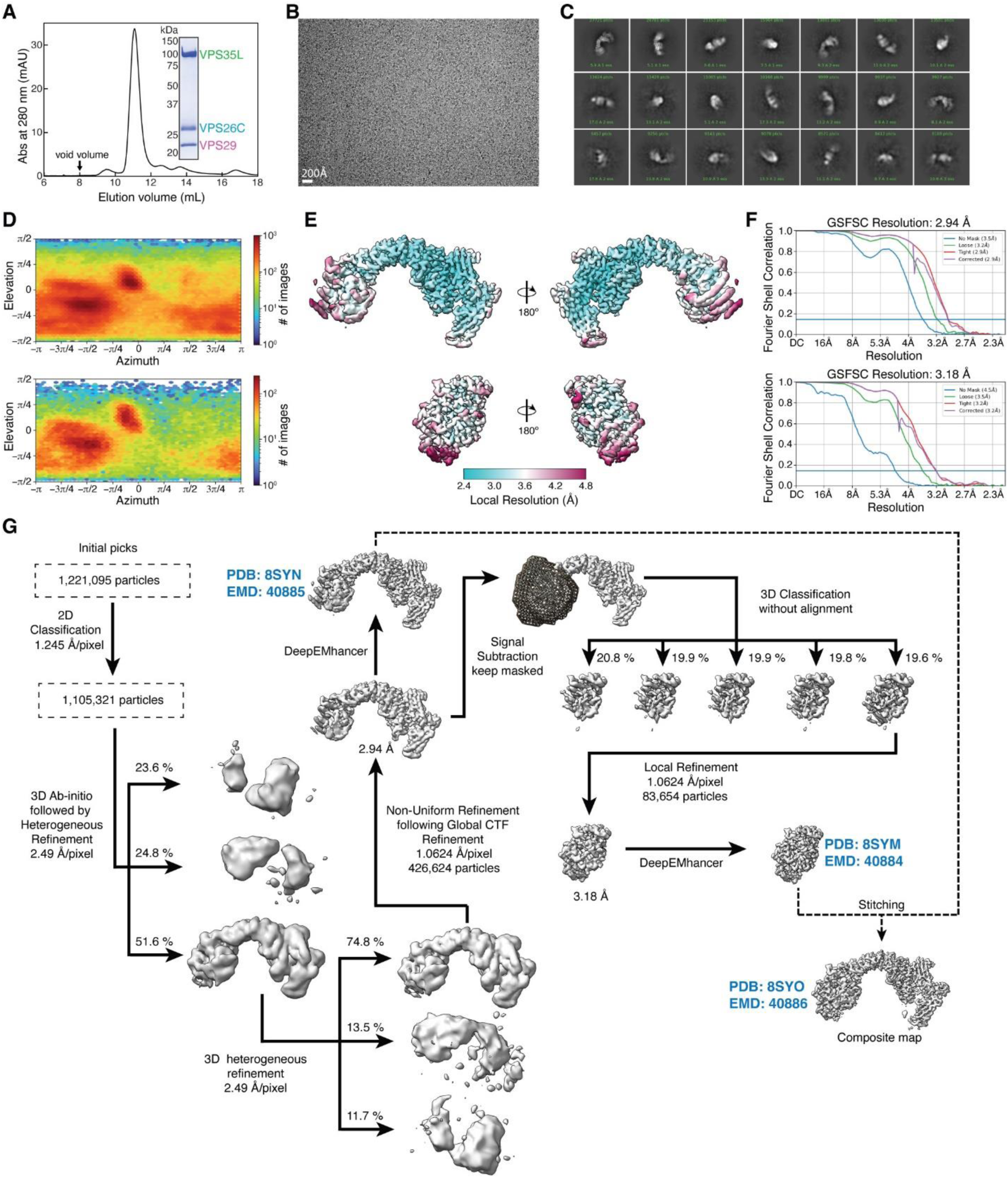
Purification and cryo-EM structural determination of Retriever. **(A)** Gel filtration chromatography of the purified Retriever complex. **(B)** Representative cryo-EM micrograph. **(C)** Representative 2D class averages. **(D)** Euler angle distribution plots for Retriever (upper) and the locally refined VPS29 with the NT “belt” peptide and the CT region of VPS35L (lower). **(E)** Local resolution map of Retriever (upper) and the locally refined VPS29 with the NT “belt” peptide and the CT region of VPS35L (lower). **(F)** Fourier Shell Correlation (FSC) plot for Retriever (upper) and the locally refined VPS29 with the NT “belt” peptide and the CT region of VPS35L (lower). **(G)** Schematic showing cryo-EM data processing steps for obtaining 3D reconstruction of Retriever. The three maps deposited to PDB/EMDB are labeled.

**Extended Data Fig. 2.**
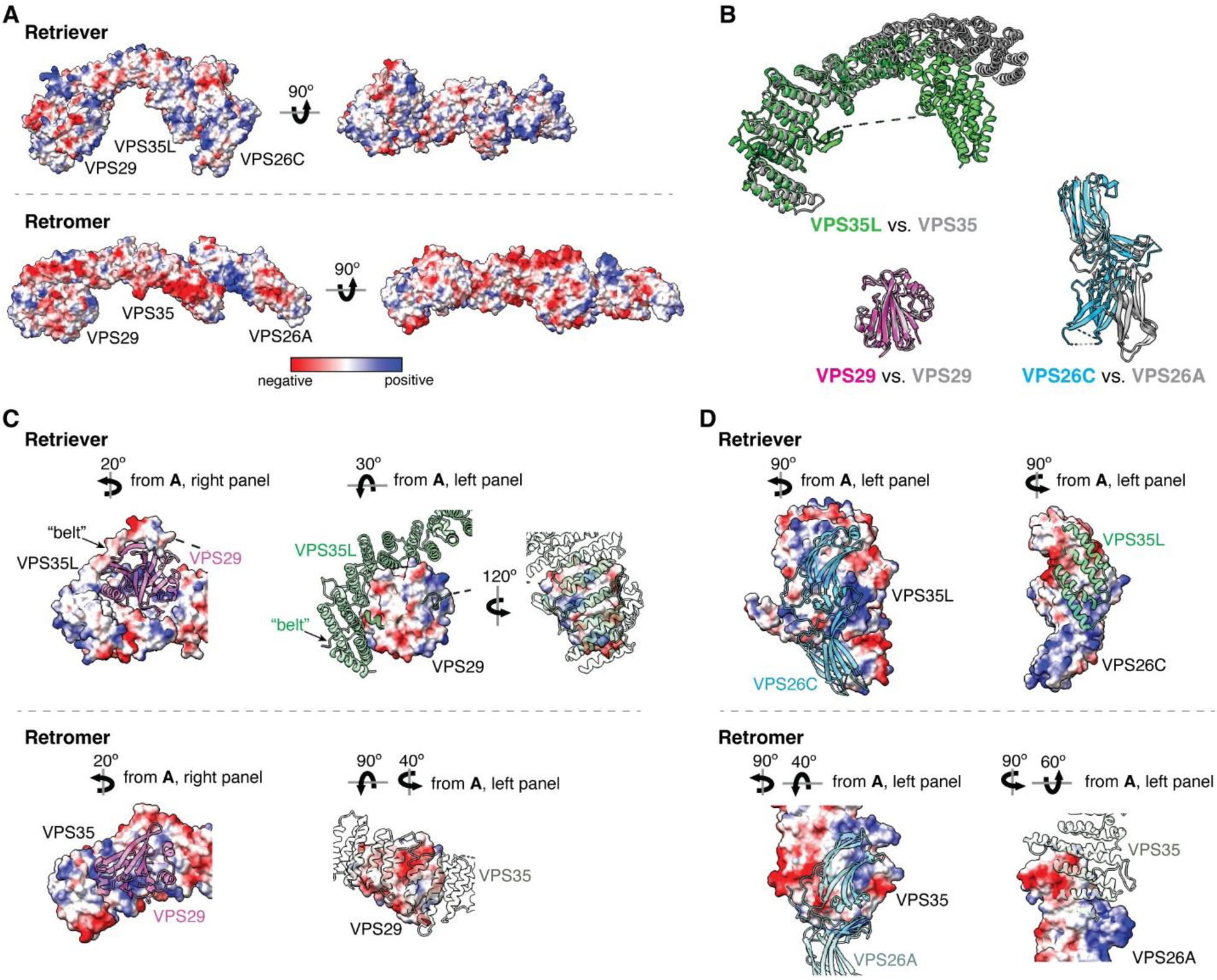
Structural comparison between Retriever and Retromer. **(A)** Surface representation of electrostatic potentials of Retriever vs. Retromer (PDB: 7U6F). **(B)** Superimpose of individual subunits from Retriever (colored) vs. Retromer (gray). **(C)** Intermolecular interface between VPS35L and VPS29 vs. VPS35 and VPS29, shown as surface representations of electrostatic potentials. (D) same as in (C), with VPS35L and VPS26C vs. VPS35 and VPS26A.

**Extended Data Fig. 3.**
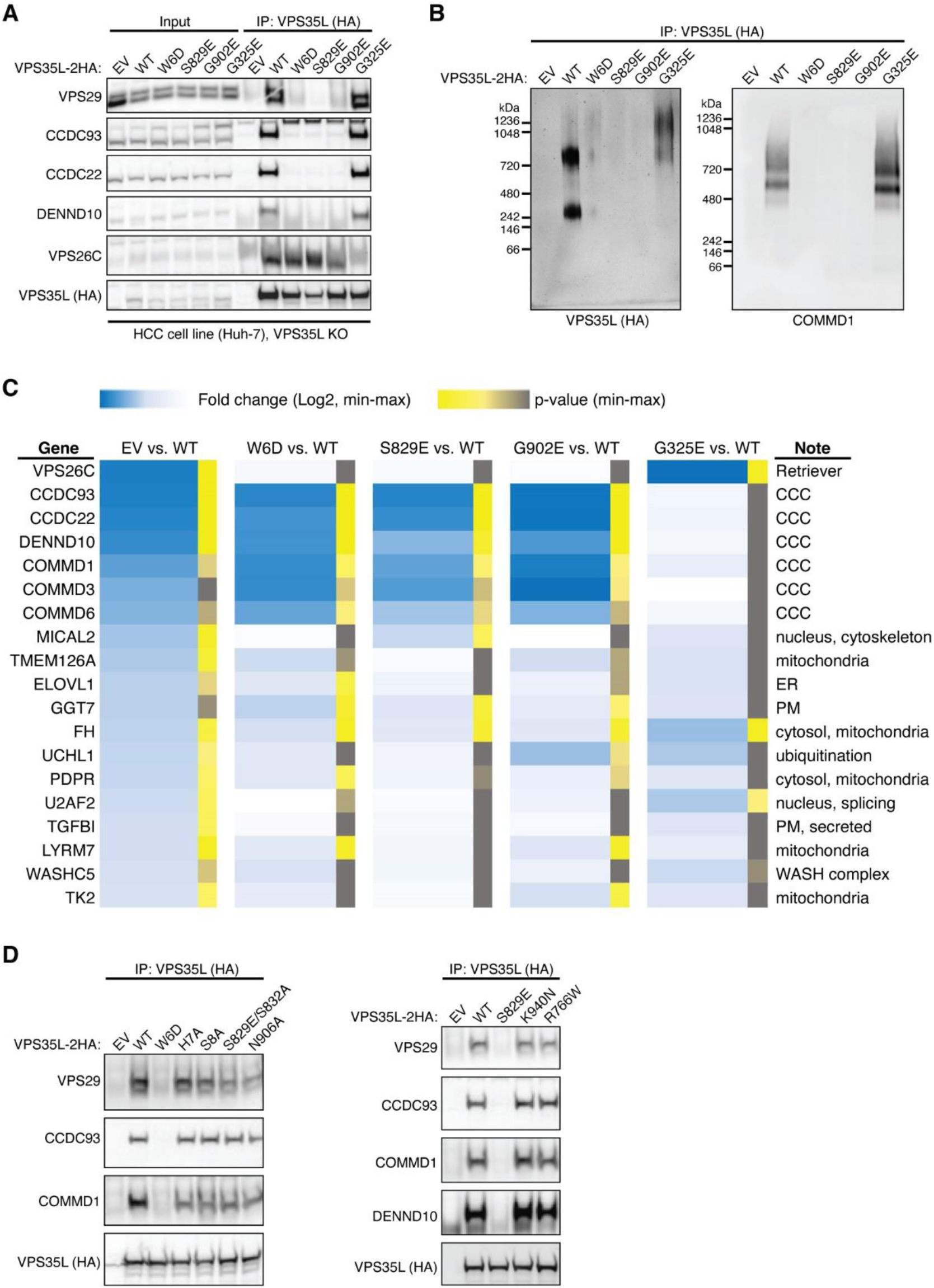
Cellular and proteomic analysis of VPS35L mutants. **(A)** Huh-7 hepatocellular carcinoma cells carrying the indicated mutations in VPS35L (EV, empty vector). Immunoprecipitation of VPS35L followed by western blot for the indicated proteins is shown. **(B)** Immunoprecipitation of VPS35L followed by competitive elution of native complexes using HA peptide, and separation of the complexes in blue native gels. After transfer, the complexes were immunoblotted with the indicated antibodies. **(C)** Heatmap representation of protein-protein interaction results using proteomics. VPS35L was immunoprecipitated from the indicated Huh-7 stable cell lines (in triplicate samples) and the results are expressed as fold compared to Huh-7 control cells (darker blue depicts greater fold difference). Statistical significance is indicated in color scale (yellow indicating p<0.05, and grey indicating p>0.05). (D) Immunoprecipitation of VPS35L carrying indicated point mutations expressed in HEK293T cells and immunoblotting for the indicated proteins.

**Extended Data Fig. 4.**
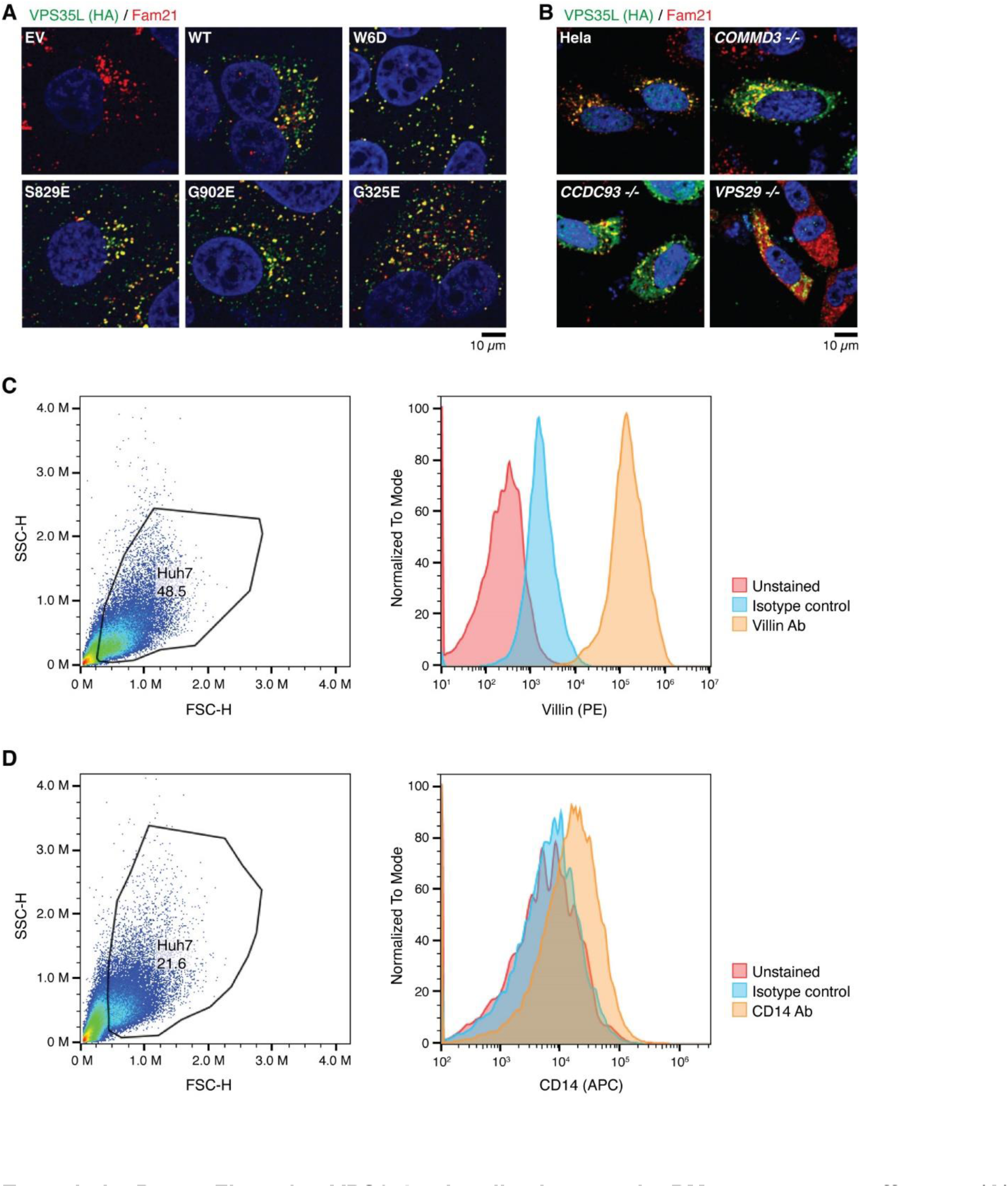
VPS35L localization and PM proteome effects. **(A)** Immunofluorescence staining for VPS35L (green channel, using HA antibody), FAM21 (red channel), and nuclei (DAPI, blue channel) in the indicated stable Huh-7 cell lines. **(B)** Immunofluorescence staining for VPS35L (green channel, using HA antibody), FAM21 (red channel), and nuclei (DAPI, blue channel) in the indicated HeLa knockout cell lines transfected with HA-VPS35L. **(C-D)** Representative gating and acquisition parameters for Villin and CD14 staining by flow cytometry.

**Extended Data Fig. 5.**
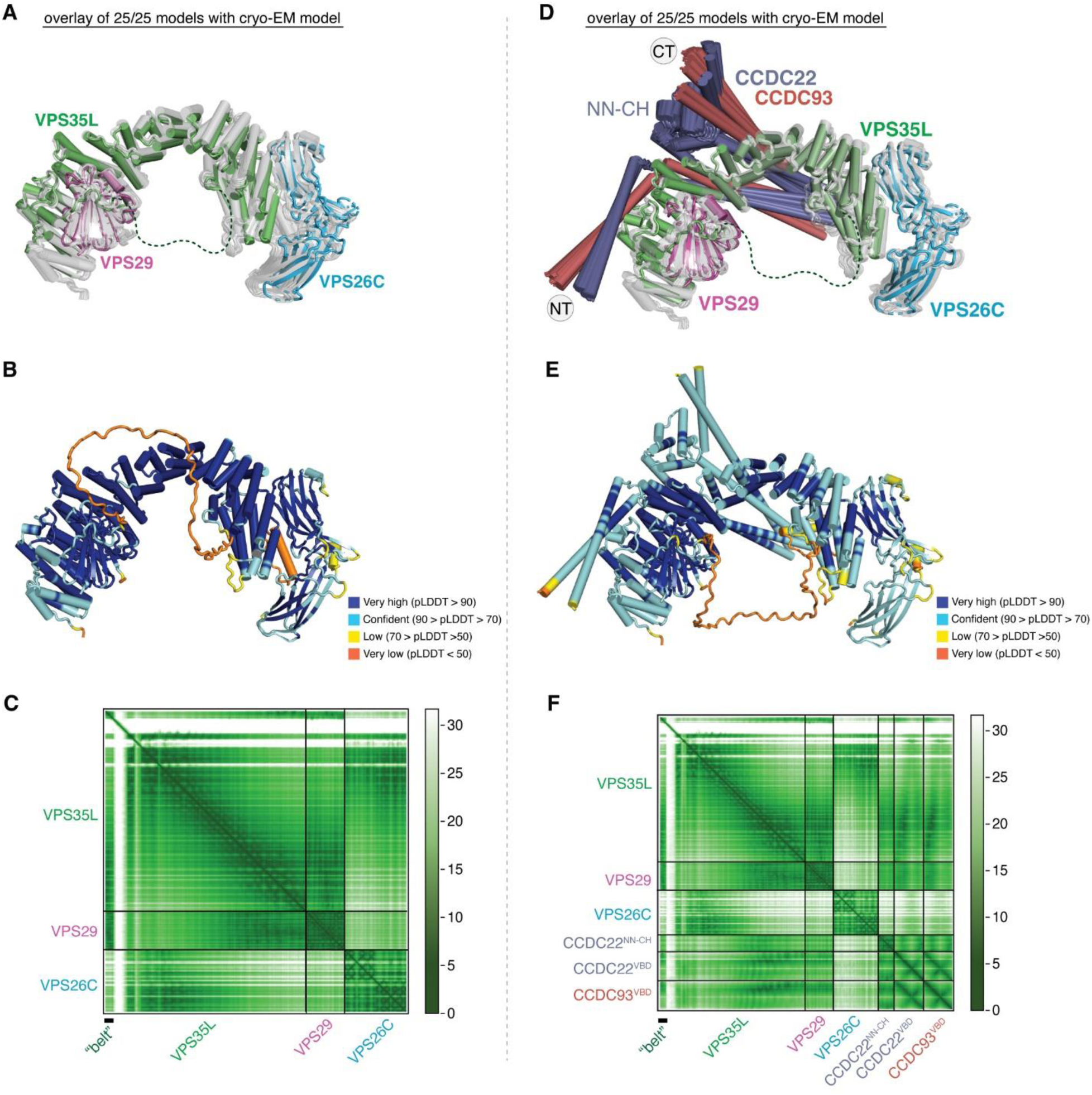
AlphaFold Multimer prediction of CCDC22-CCDC93 binding to Retriever. **(A & D)** Overlay of all 25 AlphaFold Multimer models of Retriever alone **(A)** or CCDC22-CCDC93-Retriever **(D)** with the cryo-EM model of Retriever. AFM models of Retriever are grey. **(B & E)** Representative AFM models colored using pLDDT scores. High scores indicate high reliability in local structure prediction. **(C & F)** PAE score matrix of the AFM model shown in (B & E). Low scores (deep color) indicate high reliability in the relative position in the 3D space. Boundaries of protein sequences and important structure regions are indicated.

**Extended Data Fig. 6.**
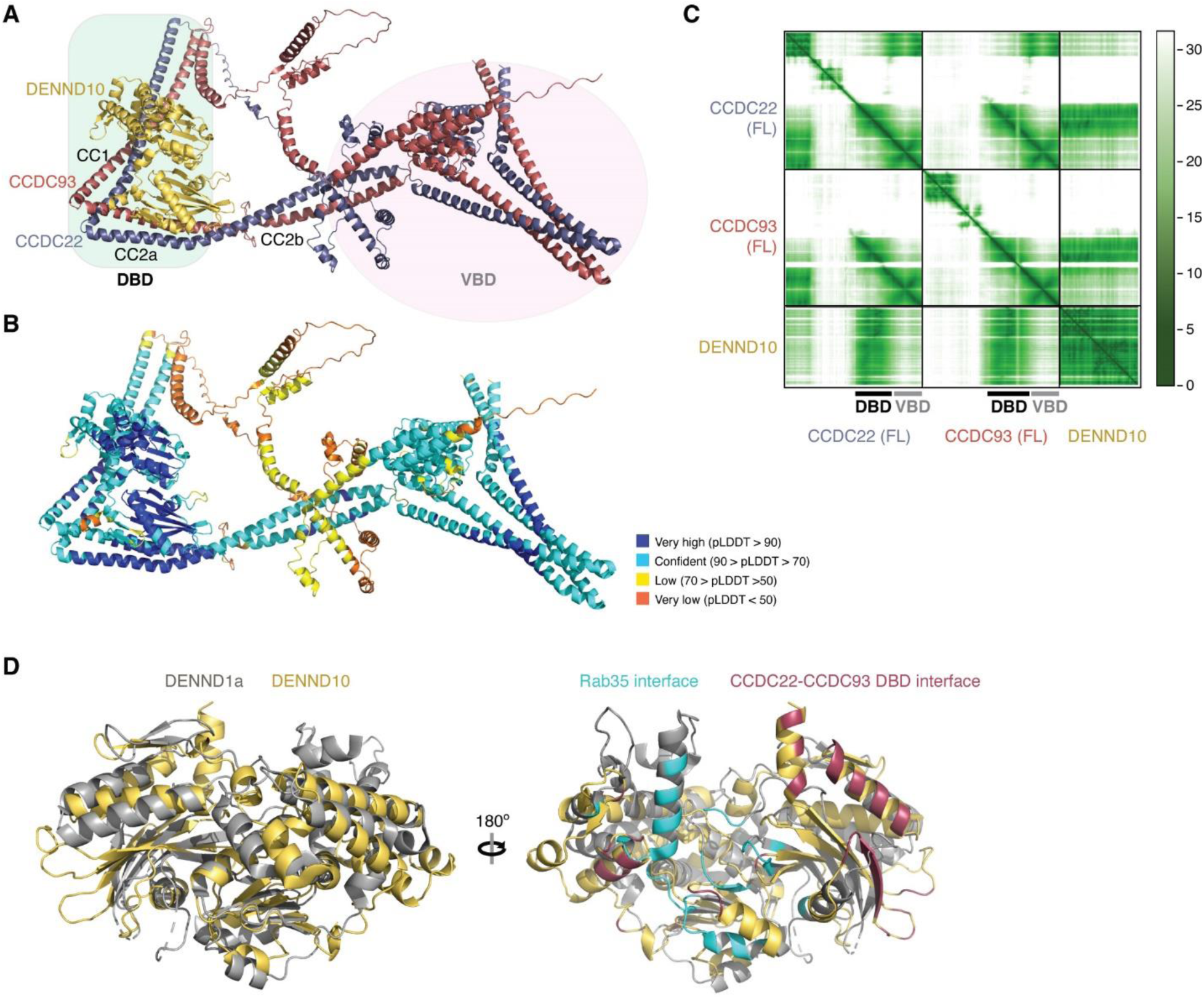
AlphaFold Multimer prediction of CCDC22-CCDC93 binding to DENND10. **(A)** AlphaFold Multimer prediction of DENND10 binding to full-length (FL) CCDC22-CCDC93. **(B)** Representative AFM models colored using pLDDT scores. **(C)** PAE score matrix of the AFM model shown in (B). Boundaries of protein sequences and important structure regions are indicated. **(D)** Superimpose of the AFM model of DENND10 with the crystal structure of DENND1a (PDB: 6EKK). Rab35 binding surface of DENND1a and CCDC22-CCDC93 binding surface of DENN10 are colored to show the partial overlap of the two surfaces.

**Extended Data Fig. 7.**
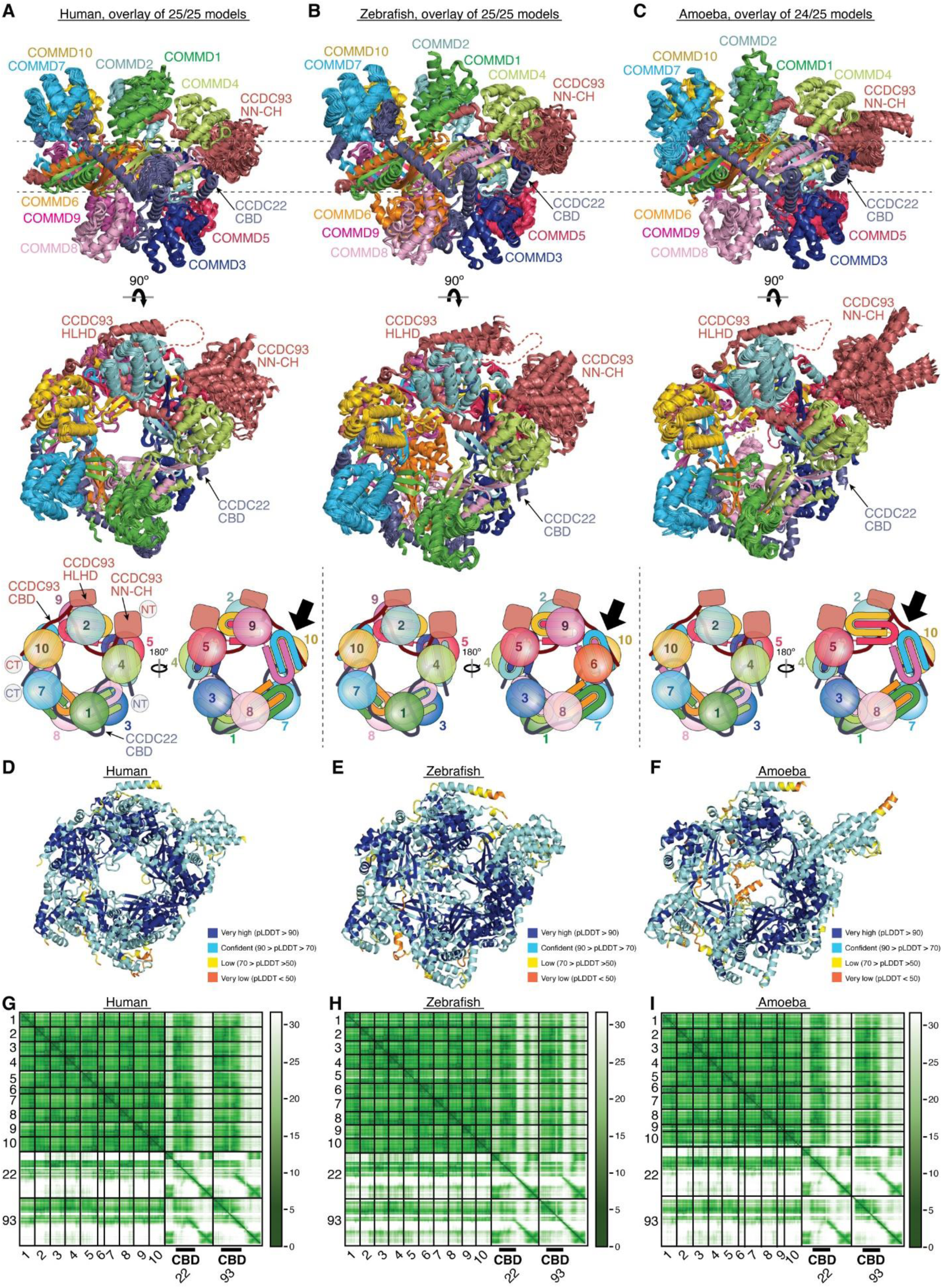
AlphaFold Multimer prediction of CCDC22-CCDC93 binding to COMMD. **(A-C)** Overlays of AlphaFold Multimer models and schematic showing CCDC22-CCDC93 binding to COMMD decamer ring for proteins from Human (A), Zebrafish (B), and Amoeba (*Dictyostelium*) (C). **(D-F)** Representative AFM models colored using pLDDT scores. **(G-I)** PAE score matrices of the AFM models shown in (D-F). Boundaries of protein sequences and important structure regions are indicated.

**Extended Table 1:** Proteomic analysis of complexes resolved by blue native electrophoresis. CRISPR/Cas9 knockout cells stably rescued with HA-tagged COMMD1 (C1) or HA-tagged VPS26C (26C), were used to immunoisolate CCC and Retriever. The complexes were eluted under native conditions (with HA peptide), resolved in blue native gels, and corresponding bands were subjected to proteomics with iBaqq quantification. Average ratios for each protein against VPS35L, CCDC22, CCDC93, COMMD4 and COMMD9 were calculated to estimate molar ratios.

| 720 MW complex |  | Spectral counts / MW<br>(sample name) |  |  | Ratio to<br>VPS35L | Ratio to<br>CCDC22 | Ratio to<br>COMMD9 |
| --- | --- | --- | --- | --- | --- | --- | --- |
| MW | Protein | Control | COMMD1 IP | VPS26C IP |  |  |  |
| 109,563 | VPS35L | 1822 | 1748200 | 1422000 | 1.000 | 0.446 |  |
| 33,010 | VPS26C | 0 | 530920 | 457950 | 0.312 | 0.139 |  |
| 20,496 | VPS29 | 75264 | 138070 | 144540 | 0.089 | 0.040 |  |
| 40,513 | DENND10 | 3720.5 | 792910 | 521280 |  | 0.185 |  |
| 70,756 | CCDC22 | 2345.7 | 4090000 | 3015300 |  | 1.000 |  |
| 73,198 | CCDC93 | 0 | 2588900 | 2013300 |  | 0.648 |  |
| 21,819 | COMMD9 | 0 | 2003200 | 1968900 |  | 0.559 | 1.000 |
| 22,966 | COMMD10 | 0 | 2415900 | 1909600 |  | 0.609 | 1.089 |
| 21,764 | COMMD4 | 0 | 1150600 | 1127600 |  | 0.321 | 0.574 |
| 24,670 | COMMD5 | 0 | 1075200 | 1264900 |  | 0.329 | 0.589 |
| 22,151 | COMMD3 | 0 | 1529200 | 1065500 |  | 0.365 | 0.653 |
| 21,178 | COMMD1 | 0 | 1899100 | 1641000 |  | 0.498 | 0.891 |
| 22,745 | COMMD2 | 0 | 1247500 | 1061900 |  | 0.325 | 0.581 |
| 21,090 | COMMD8 | 0 | 751900 | 596320 |  | 0.190 | 0.339 |
| 22,540 | COMMD7 | 0 | 1003500 | 700460 |  | 0.240 | 0.429 |
| 9,638 | COMMD6 | 0 | 1581400 | 1202800 |  | 0.392 | 0.701 |

| 480 MW complex |  | Spectral counts / MW<br>(sample name) |  |  | Ratio to<br>VPS35L | Ratio to<br>CCDC22 | Ratio to<br>COMMD4 |
| --- | --- | --- | --- | --- | --- | --- | --- |
| MW | Protein | Control | COMMD1 IP | VPS26C IP |  |  |  |
| 109,563 | VPS35L | 6365.7 | 852460 | 915390 | 1.000 | 0.120 |  |
| 33,010 | VPS26C | 0 | 361700 | 336020 | 0.395 | 0.048 |  |
| 20,496 | VPS29 | 67894 | 229980 | 392090 | 0.352 | 0.042 |  |
| 40,513 | DENND10 | 0 | 1706100 | 660430 |  | 0.161 |  |
| 70,756 | CCDC22 | 0 | 8984600 | 5699900 |  | 1.000 |  |
| 73,198 | CCDC93 | 0 | 4502000 | 2461300 |  | 0.474 |  |
| 21,764 | COMMD4 | 0 | 3421700 | 2212600 |  | 0.384 | 1.000 |
| 21,819 | COMMD9 | 0 | 4004700 | 2489000 |  | 0.442 | 1.153 |
| 22,151 | COMMD3 | 0 | 4270600 | 2278200 |  | 0.446 | 1.162 |
| 24,670 | COMMD5 | 0 | 2685500 | 1786300 |  | 0.305 | 0.794 |
| 22,745 | COMMD2 | 0 | 2876700 | 1956600 |  | 0.329 | 0.858 |
| 21,178 | COMMD1 | 0 | 4035300 | 2739700 |  | 0.461 | 1.202 |
| 22,966 | COMMD10 | 0 | 4248800 | 2980500 |  | 0.492 | 1.283 |
| 21,090 | COMMD8 | 0 | 1549600 | 1319200 |  | 0.195 | 0.509 |
| 22,540 | COMMD7 | 0 | 2559000 | 1340400 |  | 0.266 | 0.692 |
| 9,638 | COMMD6 | 0 | 3698000 | 2411000 |  | 0.416 | 1.084 |

| 240 MW complex |  | Spectral counts / MW<br>(sample name) |  |  | Ratio to<br>VPS35L | Ratio to<br>CCDC93 | Ratio to<br>COMMD9 |
| --- | --- | --- | --- | --- | --- | --- | --- |
| MW | Protein | Control | COMMD1 IP | VPS26C IP |  |  |  |
| 109,563 | VPS35L | 0 | 2020800 | 1207400 | 1.000 | 0.632 |  |
| 33,010 | VPS26C | 0 | 1428200 | 720550 | 0.666 | 0.421 |  |
| 20,496 | VPS29 | 90357 | 1113600 | 1333900 | 0.758 | 0.479 |  |
| 40,513 | DENND10 | 0 | 690080 | 223770 |  | 0.179 |  |
| 73,198 | CCDC93 | 0 | 3679300 | 1427000 |  | 1.000 |  |
| 70,756 | CCDC22 | 0 | 1688300 | 309660 |  | 0.391 |  |
| 21,819 | COMMD9 | 0 | 2943600 | 1454200 |  | 0.861 | 1.000 |
| 24,670 | COMMD5 | 0 | 2450000 | 972230 |  | 0.670 | 0.778 |
| 22,966 | COMMD10 | 0 | 3433200 | 1536200 |  | 0.973 | 1.130 |
| 21,764 | COMMD4 | 0 | 1215600 | 120040 |  | 0.262 | 0.304 |
| 22,540 | COMMD7 | 0 | 1375800 | 592580 |  | 0.385 | 0.448 |
| 21,178 | COMMD1 | 0 | 1019100 | 148690 |  | 0.229 | 0.266 |
| 9,638 | COMMD6 | 0 | 1086100 | 75972 |  | 0.228 | 0.264 |
| 22,151 | COMMD3 | 0 | 782380 | 100480 |  | 0.173 | 0.201 |
| 21,090 | COMMD8 | 0 | 810520 | 111830 |  | 0.181 | 0.210 |
| 22,745 | COMMD2 | 0 | 739430 | 127670 |  | 0.170 | 0.197 |

**Extended Table 2:** Cancer associated mutations ranked by predicted severity. Mutations downloaded from the COSMIC database were analyzed using SNAP2. Shown here are mutations with high likelihood of functional impairment (score>50, left), as well as mutations with moderate likelihood of functional impairment (score1-49, right).

| Position | CDS Mutation | AA Mutation | SNAP2 score | Occurrence |
| --- | --- | --- | --- | --- |
| **Mutations with high likelihood of functional impairment (SNAP2 score > 50)** |  |  |  |  |
| 87 | c.259G>T | p.D87Y | 56 | 1 |
| 117 | c.351G>T | p.W117C | 67 | 1 |
| 121 | c.361C>T | p.R121W | 79 | 1 |
| 121 | c.362G>A | p.R121Q | 50 | 1 |
| 128 | c.382T>C | p.Y128H | 59 | 1 |
| 132 | c.394G>A | p.E132K | 53 | 2 |
| 159 | c.475C>T | p.R159W | 55 | 1 |
| 161 | c.482G>A | p.R161Q | 53 | 1 |
| 181 | c.542A>C | p.Q181P | 58 | 1 |
| 226 | c.676T>G | p.F226V | 52 | 1 |
| 227 | c.680A>G | p.Y227C | 81 | 2 |
| 229 | c.685A>C | p.S229R | 73 | 1 |
| 236 | c.706G>T | p.D236Y | 81 | 1 |
| 246 | c.736T>C | p.Y246H | 54 | 3 |
| 248 | c.742C>G | p.R248G | 84 | 2 |
| 280 | c.839G>T | p.W280L | 85 | 1 |
| 283 | c.849G>C | p.K283N | 66 | 1 |
| 288 | c.863G>T | p.R288M | 74 | 1 |
| 293 | c.878G>T | p.R293I | 89 | 2 |
| 297 | c.891G>T | p.E297D | 71 | 1 |
| 299 | c.896C>T | p.S299F | 63 | 1 |
| 318 | c.953G>T | p.R318L | 90 | 1 |
| 318 | c.953G>A | p.R318Q | 86 | 1 |
| 324 | c.971G>C | p.R324T | 86 | 1 |
| 325 | c.974G>A | p.G325E | 94 | 1 |
| 329 | c.985C>T | p.P329S | 77 | 1 |
| 336 | c.1007G>A | p.R336H | 55 | 1 |
| 360 | c.1078G>T | p.D360Y | 63 | 1 |
| 391 | c.1171T>C | p.Y391H | 68 | 1 |
| 436 | c.1306C>T | p.R436W | 83 | 3 |
| 466 | c.1397G>T | p.R466L | 61 | 1 |
| 469 | c.1406G>A | p.G469E | 92 | 1 |
| 469 | c.1406G>T | p.G469V | 86 | 1 |
| 507 | c.1519G>A | p.E507K | 55 | 1 |
| 516 | c.1546C>G | p.H516D | 71 | 1 |
| 582 | c.1744C>T | p.R582W | 86 | 2 |
| 582 | c.1745G>T | p.R582L | 75 | 1 |
| 582 | c.1745G>A | p.R582Q | 58 | 4 |
| 590 | c.1769T>C | p.M590T | 52 | 1 |
| 610 | c.1829T>A | p.L610H | 76 | 1 |
| 636 | c.1908G>C | p.L636F | 58 | 1 |
| 651 | c.1951G>A | p.E651K | 77 | 1 |
| 665 | c.1995C>G | p.C665W | 55 | 1 |
| 682 | c.2044G>C | p.A682P | 64 | 2 |
| 702 | c.2105G>A | p.R702Q | 56 | 1 |
| 704 | c.2111G>A | p.C704Y | 92 | 1 |
| 714 | c.2141C>T | p.S714F | 77 | 1 |
| 721 | c.2162G>A | p.R721H | 64 | 2 |
| 742 | c.2224G>T | p.D742Y | 85 | 1 |
| 742 | c.2225A>C | p.D742A | 66 | 1 |
| 766 | c.2296C>T | p.R766W | 60 | 2 |
| 769 | c.2306A>T | p.E769V | 67 | 1 |
| 787 | c.2359C>A | p.P787T | 76 | 1 |
| 807 | c.2419G>T | p.D807Y | 57 | 1 |
| 811 | c.2432A>G | p.E811G | 61 | 1 |
| 846 | c.2536A>G | p.N846D | 79 | 1 |
| 851 | c.2551G>A | p.G851R | 74 | 1 |
| 856 | c.2568C>A | p.F856L | 55 | 1 |
| 857 | c.2570T>C | p.L857P | 62 | 1 |
| 902 | c.2705G>A | p.G902E | 81 | 1 |
| **Mutations with moderate likelihood of functional impairment (SNAP2 score > 0 & <50)** |  |  |  |  |
| 33 | c.98A>G | p.H33R | 11 | 1 |
| 39 | c.115A>C | p.T39P | 17 | 1 |
| 55 | c.164C>A | p.S55Y | 31 | 1 |
| 74 | c.220G>A | p.D74N | 11 | 2 |
| 75 | c.223G>T | p.G75W | 30 | 1 |
| 102 | c.304C>T | p.R102C | 4 | 1 |
| 113 | c.337G>T | p.D113Y | 43 | 2 |
| 124 | c.372C>G | p.I124M | 24 | 1 |
| 129 | c.385A>G | p.T129A | 28 | 2 |
| 131 | c.392C>T | p.T131I | 30 | 3 |
| 141 | c.422G>C | p.G141A | 3 | 2 |
| 177 | c.531G>C | p.L177F | 23 | 1 |
| 181 | c.541C>A | p.Q181K | 48 | 2 |
| 203 | c.609G>T | p.Q203H | 23 | 2 |
| 206 | c.618G>T | p.K206N | 29 | 1 |
| 222 | c.665G>T | p.S222I | 40 | 1 |
| 234 | c.702C>G | p.I234M | 5 | 1 |
| 244 | c.730C>A | p.L244I | 3 | 1 |
| 271 | c.811G>A | p.D271N | 40 | 2 |
| 281 | c.841T>G | p.F281V | 35 | 2 |
| 292 | c.874C>T | p.P292S | 12 | 4 |
| 304 | c.912C>G | p.N304K | 24 | 1 |
| 322 | c.966G>A | p.M322I | 34 | 1 |
| 343 | c.1028G>C | p.G343A | 44 | 1 |
| 357 | c.1070A>C | p.N357T | 7 | 1 |
| 368 | c.1103T>G | p.I368S | 25 | 1 |
| 394 | c.1180G>A | p.A394T | 36 | 1 |
| 400 | c.1199A>C | p.Q400P | 25 | 1 |
| 403 | c.1208C>T | p.S403F | 18 | 1 |
| 431 | c.1292T>C | p.V431A | 22 | 1 |
| 436 | c.1307G>A | p.R436Q | 33 | 1 |
| 441 | c.1322C>T | p.A441V | 44 | 1 |
| 456 | c.1367A>G | p.E456G | 23 | 1 |
| 466 | c.1397G>A | p.R466Q | 47 | 1 |
| 488 | c.1462G>A | p.E488K | 19 | 1 |
| 537 | c.1609C>T | p.R537C | 34 | 1 |
| 571 | c.1711C>A | p.P571T | 4 | 1 |
| 575 | c.1725G>T | p.M575I | 9 | 1 |
| 599 | c.1795G>A | p.E599K | 20 | 1 |
| 604 | c.1811C>A | p.P604Q | 30 | 1 |
| 634 | c.1901C>T | p.S634L | 17 | 1 |
| 635 | c.1903T>C | p.Y635H | 7 | 1 |
| 645 | c.1934C>T | p.S645F | 32 | 4 |
| 664 | c.1990T>C | p.F664L | 16 | 2 |
| 689 | c.2066T>C | p.M689T | 42 | 1 |
| 693 | c.2077C>A | p.H693N | 25 | 1 |
| 693 | c.2077C>T | p.H693Y | 17 | 1 |
| 730 | c.2190G>T | p.Q730H | 30 | 1 |
| 753 | c.2258C>T | p.P753L | 1 | 1 |
| 754 | c.2261A>C | p.E754A | 32 | 1 |
| 756 | c.2267C>T | p.P756L | 41 | 1 |
| 789 | c.2365C>T | p.H789Y | 48 | 1 |
| 819 | c.2455C>T | p.R819C | 14 | 1 |
| 845 | c.2534C>T | p.S845F | 45 | 2 |
| 881 | c.2641G>T | p.D881Y | 39 | 1 |
| 886 | c.2656C>T | p.R886C | 27 | 1 |
| 905 | c.2713C>T | p.R905C | 5 | 3 |
| 924 | c.2771G>A | p.G924D | 33 | 1 |
| 927 | c.2779G>A | p.D927N | 8 | 2 |
| 930 | c.2789C>T | p.T930I | 13 | 1 |
| 940 | c.2820G>T | p.K940N | 11 | 1 |
| 951 | c.2852C>T | p.T951M | 7 | 3 |
| 951 | c.2852C>G | p.T951R | 6 | 1 |
| **Mutations with low likelihood of functional impairment (SNAP2 score < 0)** |  |  |  |  |
| 2 | c.5C>A | p.A2D | -24 | 2 |
| 2 | c.5C>T | p.A2V | -94 | 1 |
| 3 | c.7G>A | p.V3I | -94 | 1 |
| 11 | c.32G>A | p.R11K | -13 | 2 |
| 19 | c.56C>T | p.S19L | -33 | 1 |
| 20 | c.59G>A | p.C20Y | -69 | 1 |
| 31 | c.91G>A | p.D31N | -51 | 1 |
| 38 | c.114A>G | p.I38M | -70 | 2 |
| 44 | c.132G>C | p.K44N | -49 | 1 |
| 51 | c.151A>G | p.K51E | -41 | 1 |
| 52 | c.154G>A | p.G52R | -7 | 1 |
| 56 | c.167C>T | p.S56F | 0 | 2 |
| 59 | c.176C>T | p.S59F | -4 | 1 |
| 60 | c.179C>T | p.S60F | -11 | 1 |
| 65 | c.193G>A | p.V65M | -71 | 1 |
| 66 | c.196G>A | p.V66M | -51 | 1 |
| 66 | c.196G>T | p.V66L | -55 | 1 |
| 71 | c.212G>C | p.S71T | -75 | 1 |
| 72 | c.214G>A | p.V72I | -73 | 2 |
| 73 | c.217C>T | p.L73F | -14 | 2 |
| 73 | c.217C>A | p.L73I | -21 | 1 |
| 88 | c.263C>T | p.P88L | -50 | 1 |
| 89 | c.265G>A | p.A89T | -88 | 1 |
| 89 | c.265G>T | p.A89S | -92 | 1 |
| 94 | c.280G>A | p.A94T | -39 | 1 |
| 97 | c.290G>A | p.S97N | -31 | 1 |
| 98 | c.293C>G | p.S98C | -55 | 1 |
| 102 | c.305G>A | p.R102H | -8 | 3 |
| 103 | c.309T>A | p.D103E | -79 | 1 |
| 108 | c.323C>T | p.S108F | -32 | 3 |
| 112 | c.335C>T | p.S112L | -59 | 1 |
| 125 | c.373C>A | p.L125I | -25 | 1 |
| 167 | c.501C>A | p.D167E | -52 | 1 |
| 171 | c.511G>T | p.G171C | 0 | 1 |
| 175 | c.523G>A | p.E175K | -9 | 1 |
| 201 | c.602C>T | p.S201L | -45 | 1 |
| 250 | c.749T>C | p.F250S | -45 | 1 |
| 251 | c.752C>T | p.S251F | -11 | 1 |
| 257 | c.770G>A | p.R257H | -9 | 2 |
| 259 | c.775G>A | p.V259I | -90 | 5 |
| 300 | c.900C>G | p.I300M | -57 | 1 |
| 305 | c.914A>G | p.K305R | -82 | 1 |
| 309 | c.926A>G | p.K309R | -69 | 1 |
| 310 | c.929C>T | p.T310M | -32 | 1 |
| 344 | c.1031T>C | p.M344T | -15 | 1 |
| 346 | c.1036G>A | p.V346M | -34 | 2 |
| 347 | c.1039G>A | p.A347T | -90 | 1 |
| 352 | c.1054G>C | p.E352Q | -47 | 1 |
| 355 | c.1064A>G | p.N355S | -56 | 1 |
| 364 | c.1091C>T | p.T364M | -18 | 1 |
| 372 | c.1115C>T | p.T372M | -8 | 1 |
| 378 | c.1132G>T | p.V378L | -67 | 1 |
| 389 | c.1166C>A | p.P389H | -50 | 1 |
| 399 | c.1197C>A | p.F399L | -91 | 2 |
| 406 | c.1216G>T | p.A406S | -66 | 2 |
| 412 | c.1235C>T | p.T412I | -41 | 2 |
| 422 | c.1265G>A | p.G422E | -51 | 2 |
| 432 | c.1296G>T | p.M432I | -48 | 1 |
| 441 | c.1321G>A | p.A441T | -28 | 1 |
| 442 | c.1325C>A | p.T442K | -59 | 2 |
| 453 | c.1357G>C | p.E453Q | -90 | 1 |
| 463 | c.1387C>G | p.L463V | -62 | 2 |
| 467 | c.1399T>A | p.S467T | -82 | 1 |
| 479 | c.1435G>A | p.E479K | -61 | 1 |
| 480 | c.1439G>A | p.S480N | -93 | 1 |
| 481 | c.1442A>T | p.D481V | -9 | 1 |
| 487 | c.1460A>C | p.N487T | -7 | 1 |
| 525 | c.1573G>A | p.V525I | -82 | 1 |
| 533 | c.1597A>G | p.M533V | -9 | 1 |
| 563 | c.1687C>A | p.L563I | -71 | 1 |
| 588 | c.1763G>A | p.C588Y | -44 | 1 |
| 597 | c.1791A>C | p.Q597H | -19 | 1 |
| 600 | c.1798C>T | p.P600S | -19 | 1 |
| 626 | c.1877T>A | p.L626H | -54 | 1 |
| 639 | c.1916G>T | p.G639V | -22 | 1 |
| 644 | c.1930G>A | p.V644I | -61 | 1 |
| 648 | c.1943G>A | p.R648H | -16 | 1 |
| 659 | c.1977G>T | p.E659D | -61 | 1 |
| 680 | c.2039G>A | p.R680Q | -70 | 2 |
| 716 | c.2147C>T | p.A716V | -93 | 1 |
| 722 | c.2164C>G | p.L722V | -27 | 2 |
| 736 | c.2208G>T | p.Q736H | -69 | 1 |
| 753 | c.2257C>G | p.P753A | -35 | 1 |
| 766 | c.2297G>A | p.R766Q | -20 | 1 |
| 767 | c.2299C>T | p.P767S | -72 | 3 |
| 768 | c.2303C>T | p.S768L | -34 | 1 |
| 771 | c.2313C>A | p.F771L | -11 | 1 |
| 782 | c.2345C>T | p.T782I | -1 | 1 |
| 785 | c.2353A>T | p.I785L | -88 | 1 |
| 792 | c.2375A>G | p.H792R | -74 | 2 |
| 797 | c.2389C>G | p.L797V | -62 | 2 |
| 800 | c.2399A>C | p.E800A | -49 | 1 |
| 814 | c.2441G>A | p.S814N | -2 | 1 |
| 815 | c.2443G>A | p.D815N | -13 | 1 |
| 816 | c.2446G>A | p.E816K | -60 | 2 |
| 816 | c.2448G>C | p.E816D | -93 | 1 |
| 824 | c.2470G>A | p.V824I | -57 | 1 |
| 824 | c.2471T>C | p.V824A | -66 | 2 |
| 830 | c.2488G>A | p.A830T | -68 | 2 |
| 852 | c.2555G>T | p.G852V | -3 | 1 |
| 866 | c.2597C>T | p.T866M | -20 | 4 |
| 869 | c.2606C>T | p.A869V | -74 | 1 |
| 873 | c.2619G>T | p.E873D | -71 | 2 |
| 884 | c.2650C>A | p.L884M | -70 | 1 |
| 891 | c.2671G>A | p.G891S | -63 | 1 |
| 905 | c.2714G>A | p.R905H | -22 | 2 |
| 908 | c.2722A>C | p.K908Q | -22 | 2 |
| 914 | c.2741T>C | p.V914A | -16 | 1 |
| 922 | c.2765G>A | p.R922K | -93 | 1 |
| 924 | c.2770G>A | p.G924S | -26 | 3 |
| 925 | c.2774G>A | p.C925Y | -70 | 2 |
| 932 | c.2795T>C | p.V932A | -17 | 1 |

## Source Data

**Source data western blots:** All uncropped western blot images are provided here with the corresponding figure annotation.

**Source data image quantification files** (Fig. 4)

**Source data FACS quantification files** (Fig. 4)

