## Supplementary material for "Structural Organization of the Retriever-CCC Endosomal Recycling Complex": Tables with reagents and materials used for the study

### Supplementary information

**Supplementary Tables 1-5:** These tables contain information about constructs and reagents used for this study.

#### Supplementary Table 1. DNA constructs for recombinant protein production used in this study

| Name | Description | Source/reference | Identifier |
|---|---|---|---|
| **Individual proteins/subunits** | | | |
| VPS35L | VPS35L (1-963, full-length) in pAV5a vector | This study | pDB51 |
| VPS26C | VPS26C (1-297, full-length) in pAV5a vector | This study | pDB48 |
| VPS29-His$_6$ | VPS29-Tev-(GGS)$_2$-His$_6$ in pAV5a vector | This study | pDB47 |
| MBP-CCDC22 NN-CH-VBD | MBP-Tev-CCDC22 (1-118)-(GGSK)$_6$-CCDC22 (436-727) in pMalC2Tev vector | This study | pDB79 |
| MBP-CCDC93 VBD | MBP-Tev-hCCDC93 (442-631) in pMalC2Tev vector | This study | pDB80 |
| MBP-CCDC22 DBD | MBP-Tev-CCDC22 (280-446) in pMalC2Tev vector | This study | pDB81 |
| MBP-CCDC93 DBD | MBP-Tev-CCDC93 (305-433) in pMalC2Tev vector | This study | pDB82 |
| His$_6$-DENND10 | His$_6$-Tev-DENND10 (1-357, full-length) in pHisTev vector | This study | pDB97, pDK380 |
| MBP-CCDC22 NN-CH-VBD$^{R490D}$ | MBP-Tev-CCDC22 (1-118)-(GGSK)$_6$-CCDC22 (436-727)$^{R490D}$ in pMalC2Tev vector | This study | pDB102 |
| MBP-CCDC22 NN-CH-VBD$^{V501R}$ | MBP-Tev-CCDC22 (1-118)-(GGSK)$_6$-CCDC22 (436-727)$^{V501R}$ in pMalC2Tev vector | This study | pDB103 |
| MBP-CCDC22 NN-CH-VBD$^{R490D/V501R}$ | MBP-Tev-CCDC22 (1-118)-(GGSK)$_6$-CCDC22 (436-727)$^{R490D/V501R}$ in pMalC2Tev vector | This study | pDB104 |
| MBP-CCDC93 VBD$^{R483E}$ | MBP-Tev-CCDC93 (442-631)$^{R483E}$in pMalC2Tev vector | This study | pDB84 |
| MBP-CCDC93 VBD$^{A492W}$ | MBP-Tev-CCDC93 (442-631)$^{A492W}$in pMalC2Tev vector | This study | pDB105 |
| MBP-CCDC93 VBD$^{R483E/A492W}$ | MBP-Tev-CCDC93 (442-631)$^{R483E/A492W}$ in pMalC2Tev vector | This study | pDB106 |
| MBP-CCDC22 DBD$^{A411D/A418D/E422R}$ | MBP-Tev-CCDC22 (280-446)$^{A411D/A418D/E422R}$ in pMalC2Tev vector | This study | pDB107 |
| MBP-CCDC22 DBD$^{R425D/R433D/R436D}$ | MBP-Tev-CCDC22 (280-446)$^{R425D/R433D/R436D}$ in pMalC2Tev vector | This study | pDB108 |
| MBP-CCDC93 DBD$^{E410R}$ | MBP-Tev-CCDC93 (305-433)$^{E410R}$ in pMalC2Tev vector | This study | pDB86 |
| MBP-CCDC93 DBD$^{F403D}$ | MBP-Tev-CCDC93 (305-433)$^{F403D}$ in pMalC2Tev vector | This study | pDB109 |

| | | | |
|---|---|---|---|
| MBP-CCDC93 DBD$^{E410R/F403D}$ | MBP-Tev-CCDC93 (305-433)$^{E410R/F403D}$ in pMalC2Tev vector | This study | pDB110 |
| His$_6$-DENND10$^{W30D}$ | His$_6$-Tev-DENND10$^{W30D}$ in pHisTev vector | This study | pDB111 |
| His$_6$-DENND10$^{Y32D}$ | His$_6$-Tev-DENND10$^{Y32D}$ in pHisTev vector | This study | pDB112 |

**Supplementary Table 2. Sequences of recombinant proteins used in this study.**

Only sequences in the final product (i.e., after protease cleavage to remove the affinity tag) are shown and are annotated by corresponding colors.

**>VPS35L**
MAVFPWHSRNRNYKAEFASCRLEAVPLEFGDYHPLKPITVTESKTKKVNRKGSTSSTSSSSSSSVVDPLSSVLDGTDPLSMFAATADPAALAAAM
DSSRRKRDRDDNSVVGSDFEPWTNKRGEILARYTTTEKLSINLFMGSEKGKAGTATLAMSEKVRTRLEELDDFEEGSQKELLNLTQQDYVNRIEE
LNQSLKDAWASDQKVKALKIVIQCSKLLSDTSVIQFYPSKFVLITDILDTFGKLVYERIFSMCVDSRSVLPDHFSPENANDTAKETCLNWFFKIA
SIRELIPRFYVEASILKCNKFLSKTGISECLPRLTCMIRGIGDPLVSVYARAYLCRVGMEVAPHLKETLNKNFFDFLLTFKQIHGDTVQNQLVVQ
GVELPSYLPLYPPAMDWIFQCISYHAPEALLTEMMERCKKLGNNALLLNSVMSAFRAEFIATRSMDFIGMIKECDESGFPKHLLFRSLGLNLALA
DPPESDRLQILNEAWKVITKLKNPQDYINCAEVWVEYTCKHFTKREVNTVLADVIKHMTPDRAFEDSYPQLQLIIKKVIAHFHDFSVLFSVEKFL
PFLDMFQKESVRVEVCKCIMDAFIKHQQEPTKDPVILNALLHVCKTMHDSVNALTLEDEKRMLSYLINGFIKMVSFGRDFEQQLSFYVESRSMFC
NLEPVLVQLIHSVNRLAMETRKVMKGNHSRKTAAFVRACVAYCFITIPSLAGIFTRLNLYLHSGQVALANQCLSQADAFFKAAISLVPEVPKMIN
IDGKMRPSESFLLEFLCNFFSTLLIVPDHPEHGVLFLVRELLNVIQDYTWEDNSDEKIRIYTCVLHLLSAMSQETYLYHIDKVDSNDSLYGGDSK
FLAENNKLCETVMAQILEHLKTLAKDEALKRQSSLGLSFFNSILAHGDLRNNKLNQLSVNLWHLAQRHGCADTRTMVKTLEYIKKQSKQPDMTHL
TELALRLPLQTRT

**>VPS26C**
MGTALDIKIKRANKVYHAGEVLSGVVVISSKDSVQHQGVSLTMEGTVNLQLSAKSVGVFEAFYNSVKPIQIINSTIEMVKPGKFPSGKTEIPFEF
PLHLKGNKVLYETYHGVFVNIQYTLRCDMKRSLLAKDLTKTCEFIVHSAPQKGKFTPSPVDFTITPETLQNVKERALLPKFLLRGHLNSTNCVIT
QPLTGELVVESSEAAIRSVELQLVRVETCGCAEGYARDATEIQNIQIADGDVCRGLSVPIYMVFPRLFTCPTLETTNFKVEFEVNIVVLLHPDHL
ITENFPLKLCRI

**>VPS29-Tev-(GGS)$_2$-His$_6$** (corresponding to Q9UBQ0-2, isoform 2 in Uniprot)
MAGHRLVLVLGDLHIPHRCNSLPAKFKKLLVPGKIQHILCTGNLCTKESYDYLKTLAGDVHIVRGDFDENLNYPEQKVVTVGQFKIGLIHGHQVI
PWGDMASLALLQRQFDVDILISGHTHKFEAFEHENKFYINPGSATGAYNALETNIIPSFVLMDIQASTVVTYVYQLIGDDVKVERIEYKKPENLY
FQGGGSGGSHHHHHH

**>MBP-Tev-CCDC22 NN-CH (1-118)-(GGSK)$_6$-CCDC22 VBD(436-727)**
MKIEEGKLVIWINGDKGYNGLAEVGKKFEKDTGIKVTVEHPDKLEEKFPQVAATGDGPDIIFWAHDRFGGYAQSGLLAEITPDKAFQDKLYPFTW
DAVRYNGKLIAYPIAVEALSLIYNKDLLPNPPKTWEEIPALDKELKAKGKSALMFNLQEPYFTWPLIAADGGYAFKYENGKYDIKDVGVDNAGAK
AGLTFLVDLIKNKHMNADTDYSIAEAAFNKGETAMTINGPWAWSNIDTSKVNYGVTVLPTFKGQPSKPFVGVLSAGINAASPNKELAKEFLENYL
LTDEGLEAVNKDKPLGAVALKSYEEELAKDPRIAATMENAQKGEIMPNIPQMSAFWYAVRTAVINAASGRQTVDEALKDAQTNSSSNNNNNNNNN
NLGIEGRISEFENLYFQGHMEEADRILIHSLRQAGTAVPPDVQTLRAFTTELVVEAVVRCLRVINPAVGSGLSPLLPLAMSARFRLAMSLAQACM
DLGYPLELGYQNFLYPSEPDLRDLLLFLAERLPTDASEDADQGGSKGGSKGGSKGGSKGGSKGGSKRKLQDCRELESSRRLAEIQELHQSVRAAA
EEARRKEEVYKQLMSELETLPRDVSRLAYTQRILEIVGNIRKQKEEITKILSDTKELQKEINSLSGKLDRTFAVTDELVFKDAKKDDAVRKAYKY
LAALHENCSQLIQTIEDTGTIMREVRDLEEQIETELGKKTLSNLEKIREDYRALRQENAGLLGRVREA

**>MBP-Tev-CCDC93 VBD (442-631)**
MKIEEGKLVIWINGDKGYNGLAEVGKKFEKDTGIKVTVEHPDKLEEKFPQVAATGDGPDIIFWAHDRFGGYAQSGLLAEITPDKAFQDKLYPFTW
DAVRYNGKLIAYPIAVEALSLIYNKDLLPNPPKTWEEIPALDKELKAKGKSALMFNLQEPYFTWPLIAADGGYAFKYENGKYDIKDVGVDNAGAK
AGLTFLVDLIKNKHMNADTDYSIAEAAFNKGETAMTINGPWAWSNIDTSKVNYGVTVLPTFKGQPSKPFVGVLSAGINAASPNKELAKEFLENYL
LTDEGLEAVNKDKPLGAVALKSYEEELAKDPRIAATMENAQKGEIMPNIPQMSAFWYAVRTAVINAASGRQTVDEALKDAQTNSSSNNNNNNNNN
NLGIEGRISEFENLYFQGHMTLTSAMTHDEDLDRRYNMEKEKLYKIRLLQARRNREIAILHRKIDEVPSRAELIQYQKRFIELYRQISAVHKETK
QFFTLYNTLDDKKVYLEKEISLLNSIHENFSQAMASPAARDQFLRQMEQIVEGIKQSRMKMEKKKQENKMRRDQLNDQYLELLEKQRLYFKTVKE
FKEEGRKNEMLLSKVKAKAS

**>MBP-Tev-CCDC22 DBD (280-446)**
MKIEEGKLVIWINGDKGYNGLAEVGKKFEKDTGIKVTVEHPDKLEEKFPQVAATGDGPDIIFWAHDRFGGYAQSGLLAEITPDKAFQDKLYPFTW
DAVRYNGKLIAYPIAVEALSLIYNKDLLPNPPKTWEEIPALDKELKAKGKSALMFNLQEPYFTWPLIAADGGYAFKYENGKYDIKDVGVDNAGAK
AGLTFLVDLIKNKHMNADTDYSIAEAAFNKGETAMTINGPWAWSNIDTSKVNYGVTVLPTFKGQPSKPFVGVLSAGINAASPNKELAKEFLENYL
LTDEGLEAVNKDKPLGAVALKSYEEELAKDPRIAATMENAQKGEIMPNIPQMSAFWYAVRTAVINAASGRQTVDEALKDAQTNSSSNNNNNNNNN
NLGIEGRISEFENLYFQGHMGAGAKTGAPKGSRFTHSEKFTFHLEPQAQATQVSDVPATSRRPEQVTWAAQEQELESLREQLEGVNRSIEEVEAD
MKTLGVSFVQAESECRHSKLSTAEREQALRLKSRAVELLPDGTANLAKLQLVVENSAQRVIHLAGQWEKHRVPLLAEYRHLRKLQDCRELES

**>MBP-Tev-CCDC93 DBD (305-433)**
MKIEEGKLVIWINGDKGYNGLAEVGKKFEKDTGIKVTVEHPDKLEEKFPQVAATGDGPDIIFWAHDRFGGYAQSGLLAEITPDKAFQDKLYPFTW
DAVRYNGKLIAYPIAVEALSLIYNKDLLPNPPKTWEEIPALDKELKAKGKSALMFNLQEPYFTWPLIAADGGYAFKYENGKYDIKDVGVDNAGAK
AGLTFLVDLIKNKHMNADTDYSIAEAAFNKGETAMTINGPWAWSNIDTSKVNYGVTVLPTFKGQPSKPFVGVLSAGINAASPNKELAKEFLENYL
LTDEGLEAVNKDKPLGAVALKSYEEELAKDPRIAATMENAQKGEIMPNIPQMSAFWYAVRTAVINAASGRQTVDEALKDAQTNSSSNNNNNNNNN
NLGIEGRISEFENLYFQGHMSPEKLGTSQLHRRKVISLNKQIAQKTKHLEELRASHTSLQARYNEAKKTLTELKTYSEKLDKEQAALEKIESKAD

PSILQNLRALVAMNENLKSQEQEFKAHCREEMTRLQQEIENLKAERAPRGDEKT

**>His6-Tev-DENND10**
MGHHHHHHDYDIPTTENLYFQGSMAAAEVADTQLMLGVGLIEKDTNGEVLWVWCYPSTTATLRNLLLRKCCLTDENKLLHPFVFGQYRRTWFYIT
TIEVPDSSILKKVTHFSIVLTAKDFNPEKYAAFTRILCRMYLKHGSPVKMMESYIAVLTKGICQSEENGSFLSKDFDARKAYLAGSIKDIVSQFG
METVILHTALMLKKRIVVYHPKIEAVQEFTRTLPALVWHRQDWTILHSYVHLNADELEALQMCTGYVAGFVDLEVSNRPDLYDVFVNLAESEITI
APLAKEAMAMGKLHKEMGQLIVQSAEDPEKSESHVIQDIALKTREIFTNLAPFSEVSADGEKRVLNLEALKQKRFPPATENFLYHLAAAEQMLKI

**>MBP-Tev-CCDC22 NN-CH (1-118)-(GGSK)$_6$-CCDC22 VBD (436-727)[R490D]**
MKIEEGKLVIWINGDKGYNGLAEVGKKFEKDTGIKVTVEHPDKLEEKFPQVAATGDGPDIIFWAHDRFGGYAQSGLLAEITPDKAFQDKLYPFTW
DAVRYNGKLIAYPIAVEALSLIYNKDLLPNPPKTWEEIPALDKELKAKGKSALMFNLQEPYFTWPLIAADGGYAFKYENGKYDIKDVGVDNAGAK
AGLTFLVDLIKNKHMNADTDYSIAEAAFNKGETAMTINGPWAWSNIDTSKVNYGVTVLPTFKGQPSKPFVGVLSAGINAASPNKELAKEFLENYL
LTDEGLEAVNKDKPLGAVALKSYEEELAKDPRIAATMENAQKGEIMPNIPQMSAFWYAVRTAVINAASGRQTVDEALKDAQTNSSSNNNNNNNNN
NLGIEGRISEFENLYFQGHMEEADRILIHSLRQAGTAVPPDVQTLRAFTTELVVEAVVRCLRVINPAVGSGLSPLLPLAMSARFRLAMSLAQACM
DLGYPLELGYQNFLYPSEPDLRDLLLFLAERLPTDASEDADQGGSKGGSKGGSKGGSKGGSKGGSKRKLQDCRELESSRRLAEIQELHQSVRAAA
EEARRKEEVYKQLMSELETLPRDVS**D**LAYTQRILEIVGNIRKQKEEITKILSDTKELQKEINSLSGKLDRTFAVTDELVFKDAKKDDAVRKAYKY
LAALHENCSQLIQTIEDTGTIMREVRDLEEQIETELGKKTLSNLEKIREDYRALRQENAGLLGRVREA

**>MBP-Tev-CCDC22 NN-CH (1-118)-(GGSK)$_6$-CCDC22 VBD (436-727)[V501R]**
MKIEEGKLVIWINGDKGYNGLAEVGKKFEKDTGIKVTVEHPDKLEEKFPQVAATGDGPDIIFWAHDRFGGYAQSGLLAEITPDKAFQDKLYPFTW
DAVRYNGKLIAYPIAVEALSLIYNKDLLPNPPKTWEEIPALDKELKAKGKSALMFNLQEPYFTWPLIAADGGYAFKYENGKYDIKDVGVDNAGAK
AGLTFLVDLIKNKHMNADTDYSIAEAAFNKGETAMTINGPWAWSNIDTSKVNYGVTVLPTFKGQPSKPFVGVLSAGINAASPNKELAKEFLENYL
LTDEGLEAVNKDKPLGAVALKSYEEELAKDPRIAATMENAQKGEIMPNIPQMSAFWYAVRTAVINAASGRQTVDEALKDAQTNSSSNNNNNNNNN
NLGIEGRISEFENLYFQGHMEEADRILIHSLRQAGTAVPPDVQTLRAFTTELVVEAVVRCLRVINPAVGSGLSPLLPLAMSARFRLAMSLAQACM
DLGYPLELGYQNFLYPSEPDLRDLLLFLAERLPTDASEDADQGGSKGGSKGGSKGGSKGGSKGGSKRKLQDCRELESSRRLAEIQELHQSVRAAA
EEARRKEEVYKQLMSELETLPRDVSRLAYTQRILEI**R**GNIRKQKEEITKILSDTKELQKEINSLSGKLDRTFAVTDELVFKDAKKDDAVRKAYKY
LAALHENCSQLIQTIEDTGTIMREVRDLEEQIETELGKKTLSNLEKIREDYRALRQENAGLLGRVREA

**>MBP-Tev-CCDC22 NN-CH (1-118)-(GGSK)$_6$-CCDC22 VBD (436-727)[R490D/V501R]**
MKIEEGKLVIWINGDKGYNGLAEVGKKFEKDTGIKVTVEHPDKLEEKFPQVAATGDGPDIIFWAHDRFGGYAQSGLLAEITPDKAFQDKLYPFTW
DAVRYNGKLIAYPIAVEALSLIYNKDLLPNPPKTWEEIPALDKELKAKGKSALMFNLQEPYFTWPLIAADGGYAFKYENGKYDIKDVGVDNAGAK
AGLTFLVDLIKNKHMNADTDYSIAEAAFNKGETAMTINGPWAWSNIDTSKVNYGVTVLPTFKGQPSKPFVGVLSAGINAASPNKELAKEFLENYL
LTDEGLEAVNKDKPLGAVALKSYEEELAKDPRIAATMENAQKGEIMPNIPQMSAFWYAVRTAVINAASGRQTVDEALKDAQTNSSSNNNNNNNNN
NLGIEGRISEFENLYFQGHMEEADRILIHSLRQAGTAVPPDVQTLRAFTTELVVEAVVRCLRVINPAVGSGLSPLLPLAMSARFRLAMSLAQACM
DLGYPLELGYQNFLYPSEPDLRDLLLFLAERLPTDASEDADQGGSKGGSKGGSKGGSKGGSKGGSKRKLQDCRELESSRRLAEIQELHQSVRAAA
EEARRKEEVYKQLMSELETLPRDVS**D**LAYTQRILEI**R**GNIRKQKEEITKILSDTKELQKEINSLSGKLDRTFAVTDELVFKDAKKDDAVRKAYKY
LAALHENCSQLIQTIEDTGTIMREVRDLEEQIETELGKKTLSNLEKIREDYRALRQENAGLLGRVREA

**>MBP-Tev-CCDC93 VBD (442-631)[R483E]**
MKIEEGKLVIWINGDKGYNGLAEVGKKFEKDTGIKVTVEHPDKLEEKFPQVAATGDGPDIIFWAHDRFGGYAQSGLLAEITPDKAFQDKLYPFTW
DAVRYNGKLIAYPIAVEALSLIYNKDLLPNPPKTWEEIPALDKELKAKGKSALMFNLQEPYFTWPLIAADGGYAFKYENGKYDIKDVGVDNAGAK
AGLTFLVDLIKNKHMNADTDYSIAEAAFNKGETAMTINGPWAWSNIDTSKVNYGVTVLPTFKGQPSKPFVGVLSAGINAASPNKELAKEFLENYL
LTDEGLEAVNKDKPLGAVALKSYEEELAKDPRIAATMENAQKGEIMPNIPQMSAFWYAVRTAVINAASGRQTVDEALKDAQTNSSSNNNNNNNNN
NLGIEGRISEFENLYFQGHMTLTSAMTHDEDLDRRYNMEKEKLYKIRLLQARRNREIAILH**E**KIDEVPSRAELIQYQKRFIELYRQISAVHKETK
QFFTLYNTLDDKKVYLEKEISLLNSIHENFSQAMASPAARDQFLRQMEQIVEGIKQSRMKMEKKKQENKMRRDQLNDQYLELLEKQRLYFKTVKE
FKEEGRKNEMLLSKVKAKAS

**>MBP-Tev-CCDC93 VBD (442-631)[A492W]**
MKIEEGKLVIWINGDKGYNGLAEVGKKFEKDTGIKVTVEHPDKLEEKFPQVAATGDGPDIIFWAHDRFGGYAQSGLLAEITPDKAFQDKLYPFTW
DAVRYNGKLIAYPIAVEALSLIYNKDLLPNPPKTWEEIPALDKELKAKGKSALMFNLQEPYFTWPLIAADGGYAFKYENGKYDIKDVGVDNAGAK
AGLTFLVDLIKNKHMNADTDYSIAEAAFNKGETAMTINGPWAWSNIDTSKVNYGVTVLPTFKGQPSKPFVGVLSAGINAASPNKELAKEFLENYL
LTDEGLEAVNKDKPLGAVALKSYEEELAKDPRIAATMENAQKGEIMPNIPQMSAFWYAVRTAVINAASGRQTVDEALKDAQTNSSSNNNNNNNNN
NLGIEGRISEFENLYFQGHMTLTSAMTHDEDLDRRYNMEKEKLYKIRLLQARRNREIAILHRKIDEVPSR**W**ELIQYQKRFIELYRQISAVHKETK
QFFTLYNTLDDKKVYLEKEISLLNSIHENFSQAMASPAARDQFLRQMEQIVEGIKQSRMKMEKKKQENKMRRDQLNDQYLELLEKQRLYFKTVKE
FKEEGRKNEMLLSKVKAKAS

**>MBP-Tev-CCDC93 VBD (442-631)[R483E/A492W]**
MKIEEGKLVIWINGDKGYNGLAEVGKKFEKDTGIKVTVEHPDKLEEKFPQVAATGDGPDIIFWAHDRFGGYAQSGLLAEITPDKAFQDKLYPFTW
DAVRYNGKLIAYPIAVEALSLIYNKDLLPNPPKTWEEIPALDKELKAKGKSALMFNLQEPYFTWPLIAADGGYAFKYENGKYDIKDVGVDNAGAK
AGLTFLVDLIKNKHMNADTDYSIAEAAFNKGETAMTINGPWAWSNIDTSKVNYGVTVLPTFKGQPSKPFVGVLSAGINAASPNKELAKEFLENYL
LTDEGLEAVNKDKPLGAVALKSYEEELAKDPRIAATMENAQKGEIMPNIPQMSAFWYAVRTAVINAASGRQTVDEALKDAQTNSSSNNNNNNNNN
NLGIEGRISEFENLYFQGHMTLTSAMTHDEDLDRRYNMEKEKLYKIRLLQARRNREIAILH**E**KIDEVPSR**W**ELIQYQKRFIELYRQISAVHKETK
QFFTLYNTLDDKKVYLEKEISLLNSIHENFSQAMASPAARDQFLRQMEQIVEGIKQSRMKMEKKKQENKMRRDQLNDQYLELLEKQRLYFKTVKE
FKEEGRKNEMLLSKVKAKAS

**>MBP-Tev-CCDC22 DBD (280-446)[A411D/A418D/E422R]**
MKIEEGKLVIWINGDKGYNGLAEVGKKFEKDTGIKVTVEHPDKLEEKFPQVAATGDGPDIIFWAHDRFGGYAQSGLLAEITPDKAFQDKLYPFTW
DAVRYNGKLIAYPIAVEALSLIYNKDLLPNPPKTWEEIPALDKELKAKGKSALMFNLQEPYFTWPLIAADGGYAFKYENGKYDIKDVGVDNAGAK
AGLTFLVDLIKNKHMNADTDYSIAEAAFNKGETAMTINGPWAWSNIDTSKVNYGVTVLPTFKGQPSKPFVGVLSAGINAASPNKELAKEFLENYL
LTDEGLEAVNKDKPLGAVALKSYEEELAKDPRIAATMENAQKGEIMPNIPQMSAFWYAVRTAVINAASGRQTVDEALKDAQTNSSSNNNNNNNNN
NLGIEGRISEFENLYFQGHMGAGAKTGAPKGSRFTHSEKFTFHLEPQAQATQVSDVPATSRRPEQVTWAAQEQELESLREQLEGVNRSIEEVEAD
MKTLGVSFVQAESECRHSKLSTAEREQALRLKSRAVELLPDGTANLAKLQLVVENS**D**QRVIHL**D**GQW**R**KHRVPLLAEYRHLRKLQDCRELES

**>MBP-Tev-CCDC22 DBD (280-446)[R425D/R433D/R436D]**
MKIEEGKLVIWINGDKGYNGLAEVGKKFEKDTGIKVTVEHPDKLEEKFPQVAATGDGPDIIFWAHDRFGGYAQSGLLAEITPDKAFQDKLYPFTW
DAVRYNGKLIAYPIAVEALSLIYNKDLLPNPPKTWEEIPALDKELKAKGKSALMFNLQEPYFTWPLIAADGGYAFKYENGKYDIKDVGVDNAGAK
AGLTFLVDLIKNKHMNADTDYSIAEAAFNKGETAMTINGPWAWSNIDTSKVNYGVTVLPTFKGQPSKPFVGVLSAGINAASPNKELAKEFLENYL
LTDEGLEAVNKDKPLGAVALKSYEEELAKDPRIAATMENAQKGEIMPNIPQMSAFWYAVRTAVINAASGRQTVDEALKDAQTNSSSNNNNNNNNN

| |
|---|
| NLGIEGRISEFENLYFQGHMGAGAKTGAPKGSRFTHSEKFTFHLEPQAQATQVSDVPATSRRPEQVTWAAQEQELESLREQLEGVNRSIEEVEAD<br>MKTLGVSFVQAESECRHSKLSTAEREQALRLKSRAVELLPDGTANLAKLQLVVENSAQRVIHLAGQWEKH**D**VPLLAEY**D**HL**D**KLQDCRELES |
| **>MBP-Tev-CCDC93 DBD (305-433)**$^{F403D}$<br>MKIEEGKLVIWINGDKGYNGLAEVGKKFEKDTGIKVTVEHPDKLEEKFPQVAATGDGPDIIFWAHDRFGGYAQSGLLAEITPDKAFQDKLYPFTW<br>DAVRYNGKLIAYPIAVEALSLIYNKDLLPNPPKTWEEIPALDKELKAKGKSALMFNLQEPYFTWPLIAADGGYAFKYENGKYDIKDVGVDNAGAK<br>AGLTFLVDLIKNKHMNADTDYSIAEAAFNKGETAMTINGPWAWSNIDTSKVNYGVTVLPTFKGQPSKPFVGVLSAGINAASPNKELAKEFLENYL<br>LTDEGLEAVNKDKPLGAVALKSYEEELAKDPRIAATMENAQKGEIMPNIPQMSAFWYAVRTAVINAASGRQTVDEALKDAQTNSSSNNNNNNNNN<br>NLGIEGRISEFENLYFQGHMSPEKLGTSQLHRRKVISLNKQIAQKTKHLEELRASHTSLQARYNEAKKTLTELKTYSEKLDKEQAALEKIESKAD<br>PSILQNLRALVAMNENLKSQEQE**D**KAHCREEMTRLQQEIENLKAERAPRGDEKT |
| **>MBP-Tev-CCDC93 DBD (305-433)**$^{E410R}$<br>MKIEEGKLVIWINGDKGYNGLAEVGKKFEKDTGIKVTVEHPDKLEEKFPQVAATGDGPDIIFWAHDRFGGYAQSGLLAEITPDKAFQDKLYPFTW<br>DAVRYNGKLIAYPIAVEALSLIYNKDLLPNPPKTWEEIPALDKELKAKGKSALMFNLQEPYFTWPLIAADGGYAFKYENGKYDIKDVGVDNAGAK<br>AGLTFLVDLIKNKHMNADTDYSIAEAAFNKGETAMTINGPWAWSNIDTSKVNYGVTVLPTFKGQPSKPFVGVLSAGINAASPNKELAKEFLENYL<br>LTDEGLEAVNKDKPLGAVALKSYEEELAKDPRIAATMENAQKGEIMPNIPQMSAFWYAVRTAVINAASGRQTVDEALKDAQTNSSSNNNNNNNNN<br>NLGIEGRISEFENLYFQGHMSPEKLGTSQLHRRKVISLNKQIAQKTKHLEELRASHTSLQARYNEAKKTLTELKTYSEKLDKEQAALEKIESKAD<br>PSILQNLRALVAMNENLKSQEQEFKAHCRE**R**MTRLQQEIENLKAERAPRGDEKT |
| **>MBP-Tev-CCDC93 DBD (305-433)**$^{F403D/E410R}$<br>MKIEEGKLVIWINGDKGYNGLAEVGKKFEKDTGIKVTVEHPDKLEEKFPQVAATGDGPDIIFWAHDRFGGYAQSGLLAEITPDKAFQDKLYPFTW<br>DAVRYNGKLIAYPIAVEALSLIYNKDLLPNPPKTWEEIPALDKELKAKGKSALMFNLQEPYFTWPLIAADGGYAFKYENGKYDIKDVGVDNAGAK<br>AGLTFLVDLIKNKHMNADTDYSIAEAAFNKGETAMTINGPWAWSNIDTSKVNYGVTVLPTFKGQPSKPFVGVLSAGINAASPNKELAKEFLENYL<br>LTDEGLEAVNKDKPLGAVALKSYEEELAKDPRIAATMENAQKGEIMPNIPQMSAFWYAVRTAVINAASGRQTVDEALKDAQTNSSSNNNNNNNNN<br>NLGIEGRISEFENLYFQGHMSPEKLGTSQLHRRKVISLNKQIAQKTKHLEELRASHTSLQARYNEAKKTLTELKTYSEKLDKEQAALEKIESKAD<br>PSILQNLRALVAMNENLKSQEQE**D**KAHCRE**R**MTRLQQEIENLKAERAPRGDEKT |
| **>His$_6$-Tev-DENND10 (1-357)**$^{W30D}$<br>MGHHHHHHDYDIPTTENLYFQGSMAAAEVADTQLMLGVGLIEKDTNGEVLWV**D**CYPSTTATLRNLLLRKCCLTDENKLLHPFVFGQYRRTWFYIT<br>TIEVPDSSILKKVTHFSIVLTAKDFNPEKYAAFTRILCRMYLKHGSPVKMMESYIAVLTKGICQSEENGSFLSKDFDARKAYLAGSIKDIVSQFG<br>METVILHTALMLKKRIVVYHPKIEAVQEFTRTLPALVWHRQDWTILHSYVHLNADELEALQMCTGYVAGFVDLEVSNRPDLYDVFVNLAESEITI<br>APLAKEAMAMGKLHKEMGQLIVQSAEDPEKSESHVIQDIALKTREIFTNLAPFSEVSADGEKRVLNLEALKQKRFPPATENFLYHLAAAEQMLKI |
| **>His$_6$-Tev-DENND10 (1-357)**$^{Y32D}$<br>MGHHHHHHDYDIPTTENLYFQGSMAAAEVADTQLMLGVGLIEKDTNGEVLWVWC**D**PSTTATLRNLLLRKCCLTDENKLLHPFVFGQYRRTWFYIT<br>TIEVPDSSILKKVTHFSIVLTAKDFNPEKYAAFTRILCRMYLKHGSPVKMMESYIAVLTKGICQSEENGSFLSKDFDARKAYLAGSIKDIVSQFG<br>METVILHTALMLKKRIVVYHPKIEAVQEFTRTLPALVWHRQDWTILHSYVHLNADELEALQMCTGYVAGFVDLEVSNRPDLYDVFVNLAESEITI<br>APLAKEAMAMGKLHKEMGQLIVQSAEDPEKSESHVIQDIALKTREIFTNLAPFSEVSADGEKRVLNLEALKQKRFPPATENFLYHLAAAEQMLKI |

### Supplementary Table 3. DNA oligos used in this study

| Purpose | Identifier and sequence, all 5' to 3' |
|---|---|
| VPS35L | oDB220120-1, CTCGGTCCGAACTCCTAAAAAACCGCCACCatggccgtctttc, forward primer to extract VPS35L and add annealing arm for SLIC<br>oDB220120-2, GCATGCCTCGAGACTGCAGGCTCTAGATTAggtccttgtttgcagagggag, reverse primer to extract VPS35L and add annealing arm for SLIC<br>oDB220120-14, GGTGGCGGTTTTTTAGGAGTTC, reverse primer to open pAV5a<br>oDB220120-15, TAATCTAGAGCCTGCAGTCTCGAG, forward primer to open pAV5a |
| VPS26C | oDB220120-4, TTTGAAAACCTGTATTTTCAGGGCCATATGGGGACCGCCCTGGACATC, forward primer to extract VPS26C and add annealing arm for SLIC<br>GCTGAAGCTCTGCAGGATATAATCTAGAGCCTGCAGTCTCGAGGCATGC<br>oDB220505-1, GCATGCCTCGAGACTGCAGGCTCTAGATTATATCCTGCAGAGCTTCAGC, reverse primer to extract VPS26C and add annealing arm for SLIC<br>oDB220120-14, GGTGGCGGTTTTTTAGGAGTTC, reverse primer to open pAV5a<br>oDB220120-15, TAATCTAGAGCCTGCAGTCTCGAG, forward primer to open pAV5a |
| VPS29-His$_6$ | oDB220120-11, CTCGGTCCGAACTCCTAAAAAACCGCCACCATGGCTGGGCACAGATTG, forward primer to extract VPS29 and add annealing arm for SLIC<br>oDB220120-12,<br>GATGAGAGCCTCCACTTCCACCGCCCTGAAAATACAGGTTTTCAGGTTTTTTGTATTCGATTCGTTCTACTTTC, forward primer to extract VPS29 and add annealing arm for SLIC<br>oDB220120-13,<br>TTCAGGGCGGTGGAAGTGGAGGCTCTCATCATCATCATCATCATTAATCTAGAGCCTGCAGTCTCGAG, forward primer to open pAV5a and add SLIC annealing arm for VPS29-His6<br>oDB220120-14, GGTGGCGGTTTTTTAGGAGTTC, reverse primer to open pAV5a |
| His$_6$-DENND10 | cbyo-230123-1, ACTACCGAAAACCTGTACTTCCAG, PCR DENND10opti BamHI fw<br>Cbyo-230123-2, TAAGTGCTCGAGTTAAATCTTCAGCATC, PCR DENND10opti XhoI bw |
| MBP-CCDC22 Head-VBD$^{R490D}$ | Cbyo-230502-1, CTGGCATATACCCAGCGTATTCTG, Aliblunt for CCDC22_VBDopti R490D FW |

| | |
|---|---|
| | Cbyo-230502-2, ATCGCTAACATCACGAGGCAGGGTTTC, Aliblunt for CCDC22_VBDopti R490D BW |
| MBP-CCDC22 Head-VBD$^{V501R}$ | Cbyo-230502-3, CGTGGCAATATTCGCAAGCAGAAAGAAG, Aliblunt for CCDC22_VBDopti V501R FW<br>Cbyo-230502-4, AATTTCCAGAATACGCTGGGTATATGC, Aliblunt for CCDC22_VBDopti V501R BW |
| MBP-CCDC22 Head-VBD$^{R490D/V501R}$ | Cbyo-230502-5, TTAGCTTACACGCAACGCATCTTAGAGATCCGTGGCAATATTCGCAAGCAGAAAGAAG, Aliblunt for CCDC22_VBDopti R490D/V501R FW<br>Cbyo-230502-2, ATCGCTAACATCACGAGGCAGGGTTTC, Aliblunt for CCDC22_VBDopti R490D BW |
| MBP-CCDC93 VBD$^{A492W}$ | Cbyo-230502-6, TGGGAACTGATTCAGTATCAGAAACGTTTTATCGAAC, Aliblunt for CCDC93_VBDopti A492W FW<br>Cbyo-230502-7, ACGGCTCGGAACTTCATCAATTTTAC, Aliblunt for CCDC93_VBDopti A492W BW |
| MBP-CCDC93 VBD$^{R483E/A492W}$ | Cbyo-230502-6, TGGGAACTGATTCAGTATCAGAAACGTTTTATCGAAC, Aliblunt for CCDC93_VBDopti A492W FW<br>Cbyo-230502-8, ACGGCTCGGAACTTCATCAATTTTC, Aliblunt for CCDC93_VBDopti R483E/A492W BW |
| MBP-CCDC22 DBD$^{A411D/A418D/E422R}$ | Cbyo-230502-9, TTAGATGGACAATGGCGTAAACATCGTGTTCCGCTGCTGG, Aliblunt for CCDC22_DBDopti A411D/A418D/E422R FW<br>Cbyo-230502-10, GTGGATGACGCGTTGATCGCTATTTTCAACAACCAGCTGCAG, Aliblunt for CCDC22_DBDopti A411D/A418D/E422R BW |
| MBP-CCDC22 DBD$^{R425D/R433D/R436D}$ | Cbyo-230502-11, GAGTACGATCACTTAGATAAACTGCAGGATTGTCGTGAACTGG, Aliblunt for CCDC22_DBDopti R425D/R433D/R436D FW<br>Cbyo-230502-12, GGCTAATAATGGTACGTCATGTTTTTCCCACTGACCTGCCAG, Aliblunt for CCDC22_DBDopti R425D/R433D/R436D BW |
| MBP-CCDC93 DBD$^{F403D}$ | Cbyo-230502-13, TAAAGCCCATTGTCGTGAAGAAATGAC, Aliblunt for CCDC93_DBDopti F403D FW<br>Cbyo-230502-14, TCCTCTTGCTCTTGGCTTTTCAGATTTTCATTC, Aliblunt for CCDC93_DBDopti F403D BW |
| MBP-CCDC93 DBD$^{E410R/F403D}$ | Cbyo-230502-15, TAAAGCCCATTGTCGTGAACGTATGAC, Aliblunt for CCDC93_DBDopti E410R/F403D FW<br>Cbyo-230502-14, TCCTCTTGCTCTTGGCTTTTCAGATTTTCATTC, Aliblunt for CCDC93_DBDopti F403D BW |
| His$_6$-DENND10$^{W30D}$ | Cbyo-230502-16, ATCCGAGCACCACCGCAACAC, Aliblunt for DENND10 W30D FW<br>Cbyo-230502-17, AACAATCAACCCACAGAACTTCACCATTGGTATC, Aliblunt for DENND10 W30D BW |
| His$_6$-DENND10$^{Y32D}$ | Cbyo-230502-16, ATCCGAGCACCACCGCAACAC, Aliblunt for DENND10 W30D FW<br>Cbyo-230502-18, CACACCAAACCCACAGAACTTCAC, Aliblunt for DENND10 Y32D BW |

### Supplementary Table 4: Guide RNA sequences used for CRISPR/Cas9

| **Crispr gRNA** | | | |
|---|---|---|---|
| ***Gene target (Human)*** | ***Guide construct*** | ***Target sequence*** | ***Reference*** |
| *VPS35L* | Guide 1 | TGATGAATCTGGTTTCCCCA | This paper |
| *VPS35L* | Guide 2 | CACTAAGCTGAAGAACCCAC | Singla et al, 2019 |
| *VPS29* | Guide 1 | TAGCTGGCAAACTGTTGCAC | This paper |
| *VPS29* | Guide 2 | GACTATCTCAAGACTCTGGC | This paper |
| *VPS29* | Guide 3 | GGACATCAAGTTATTCCATG | This paper |

**Supplementary Table 5: Antibodies used in this study**

| **Primary antibodies**<br>WB, western blot; IF, immunofluorescence staining; FC, Flow Cytometry. | | | |
|---|---|---|---|
| **Target** | **Source** (host species) | **Catalog or identifying number** | **Application** |
| CCDC22 | ProteinTech Group (rabbit) | 16636-1-AP | WB |
| CCDC22 | Invitrogen (mouse) | MA5-27399 | WB |
| CCDC93 | ProteinTech Group (rabbit) | 20861-1-AP | WB |
| CD14-APC | BD Biosciences (mouse) | 555399 | FC |
| COMMD1 | Novus Biologicals (mouse) | NBP2-03755 | WB |
| COMMD2 | Custom made (rat) | UT-R 42 (Li et al., 2015) | WB |
| COMMD3 | Custom made (rat) | UT-R 43 (Li et al., 2015) | WB |
| COMMD4 | Custom made (rabbit) | UT-691 (Li et al., 2015) | WB |
| COMMD5 | ProteinTech Group (mouse) | 67043-1-IG | WB |
| COMMD6 | Custom made (rabbit) | 051-AP (Starokadomskyy et al., 2013) | WB |
| COMMD7 | Custom made (rat) | UT-R 47 (Li et al., 2015) | WB |
| COMMD8 | Custom made (guinea pig) | UT-GP 135 (Li et al., 2015) | WB |
| COMMD9 | Custom made (rabbit) | 192-AP (Starokadomskyy et al., 2013) | WB |
| COMMD10 | Santa Cruz (mouse) | sc-398798 | WB |
| DENND10 | Custom made (rabbit) | 95-110 (Singla et al., 2019) | WB |
| FAM21 | Custom made (rabbit) | MC2188 (Gomez and Billadeau, 2009) | IF |
| FLAG | Sigma (mouse) | F1804 | WB |
| HA | Biolegend (mouse) | 901502 | WB, IF |
| HA | Cell Signaling (mouse) | 2999S | WB |
| Integrin-β1 | Santa Cruz (mouse) | sc-53711 | IF |
| LAMP1 | Abcam (rabbit) | Ab24170 | IF |
| Villin | ProteinTech Group (rabbit) | 16488-1-AP | FC |
| VPS26A | Abcam (rabbit) | Ab23892 | WB |
| VPS26C | Millipore (rabbit) | ABN87 | WB |
| VPS29 | GeneTex (rabbit) | GTX104768 | WB |
| VPS35 | Abcam (goat) | Ab10099 | WB |
| VPS35L | ThermoFisher Scientific (rabbit) | PA5-28553 | WB |
| **Secondary antibodies used for immunofluorescence** | | | |
| **Fluorophore** | **Source** (target species) | **Catalog number** | |
| Alexa 488 | Invitrogen (mouse) | A11029 | |
| Alexa 555 | Invitrogen (rabbit) | A21428 | |
| Phalloidin 488 | Invitrogen | A12379 | |
